# Analysis of long-range contacts across cell types outlines a core sequence determinant of 3D genome organisation

**DOI:** 10.1101/2025.03.16.643527

**Authors:** Liezel Tamon, Zahra Fahmi, James Ashford, Rosana Collepardo-Guevara, Aleksandr B. Sahakyan

## Abstract

The sequence-driven organising principles of the 3D genome are crucial for interpreting the core effects of genomic variation and for understanding the evolution of genome organisation and function. We investigated these by isolating and analysing cell-type-persistent contacts, heavily dependent on the similarly cell-type-persistent genomic sequence. We stratified long-range contacts from a diverse group of human tissues and cell lines based on contact persistence, c_p_, reflecting their presence across cell or tissue types, presenting them as an atlas of contacts and the cell-type invariant (CETI) hubs they form across human chromosomes. Our survey of more than 300 chromatin and genome features revealed their association with c_p_, contrasting variable from persistent contacts in terms of co-localisation with genes, 3D architectural domains, epigenetic and sequence elements. We found persistent contacts to be predominantly comprised of AT-rich sequences and related to heterochromatin. A key outcome is finding a link between the experimental genomic contacts and the complementarity between pairs of contacting DNA loci. This work provides evidence for a sequence determinant of genomic contacts contributing to the decoding of the relationship between sequence and structure that is crucial for functional and evolutionary studies concerning the 3D genome organisation.

## Introduction

Within a human cell, a DNA double helix of ∼ 2 m in length is intricately organised inside a nucleus, ∼ 10 μm in diameter, while enabling proper functioning of molecular processes like replication and gene expression. According to length or size, this organisation can broadly be categorised into the **1)** scale of nucleosome- and chromatin-fibre, and of higher-order organisation, consisting of the **2)** intermediate (domain) scale (e.g. loops, topologically associating domains or TADs [1–3], lamina associated domains or LADs and compartments [4]), and the **3)** nuclear scale (e.g. chromosome territories, genome arrangement with respect to nuclear centre/periphery and nuclear bodies).

Key to understanding the relationship between the way the genome is organised and its function is the identification of factors influencing that organisation. Results from biochemical mapping methods, imaging experiments, polymer simulations, and other computational and experimental investigations have shown that a number of factors play a multiplexed role in determining the 3D genome organisation. These factors include the general polymeric nature of the DNA [5], the complex and nonuniform information the chromatin holds in the form of epigenetic modifications and other occupants [6], the molecular processes the chromatin goes through particularly during replication [7] and transcription [8], and the genomic DNA sequence [9–12]. The relationship between genomic sequence and any compounding phenomenon, like 3D genome organisation, remains a subject of much interest, as it can crucially contribute to the interpretation of naturally occurring and *de novo* genomic variations and to the understanding of the sequence-structural evolution of our genome.

The volume of work targeting the understanding of genomic contacts has steeply risen ever since the first experimental means have been reported [4,13–15] and it has established genome organisation as dynamic and stochastic at all levels (recently reviewed [16]), as opposed to being static and deterministic. The increasing amount of data from 3C and other types of methods in the context of various cell types, states, and species have outlined the multiscale nature of genome organisation [1,3,6,17], and the complex dynamics of contacts therein [16,18]. Genome organisation within a given species varies [19,20] across population [21], depending on a cell type [22], state in a cell cycle [23,24], epigenetic state [25], and throughout differentiation [26,27]. The presence of features and patterns first described using bulk data were validated, but extensive heterogeneity across single cells of the same type and population[28] was revealed, even down to individual alleles [20,29,30].

In this work, we leveraged the available large pool of experimental contact data, particularly through the availability of Hi-C data from a wide selection of human cell lines and primary tissues [22], to better understand the DNA sequence basis of genomic contact formation and 3D genome organisation. In particular, we characterised the contacts across human cell types to isolate and investigate the core, cell-type-persistent or invariant contacts, which are likely enriched in associations with similarly cell-type-invariant factors, i.e. DNA sequence-based features largely common in all the human cells.

## Materials and Methods

### Computational platforms and resources

Computer code was written in R programming language and most computations utilised the in-house high-performance computing facilities at MRC Weatherall Institute of Molecular Medicine, University of Oxford, by employing cluster nodes with 256 GB random access memory, and Intel Xeon E5-2680v3 12-core (24-thread) and Intel Xeon E7-8891v3 10-core (20-thread) processors.

### Statistical analyses

Differences between two groups were analysed using Student’s t-test and Mann-Whitney-Wilcoxon (MWW) tests. Pairwise comparisons of > 2 distributions were done using pairwise implementations of both tests in R. Alternative hypothesis was two-sided by default and the p-values reported were Benjamini-Hochberg adjusted [31]. The alpha (α) or significance level is by default 0.05.

### Contact persistence stratification

The main working dataset of genomic contacts was retrieved from 21 published Hi-C data from 14 primary tissues and 7 cell lines at 40-kb resolution, consolidated and reanalysed in Schmitt et al. [22] (GEO:GSE87112). The processed contact matrices, containing uniquely mapped reads (in correspondence with the authors), and normalised using HiCNorm [32] were used for the stratification. Only long-range contacts were used in our analyses with linear distance or contact gap ≥ 2 Mb. The c_p_ of each long-range contact was calculated as the number of cell type the contact is present in i.e. the contact had a non-zero HiCNorm c_f_ in the Hi-C contact matrix. See **Supplementary Table S5** and **S6** for counts of long-range contacts and unique regions forming these contacts per cell line or tissue, respectively.

### Atlas of persistent substructures in human chromosomes

#### Arc and network diagrams for visualising contacts

Arc and network diagrams were built using R libraries R4RNA [33] and visNetwork, respectively. In the network, a node represents a region or bin of length equal to the Hi-C resolution. Which bins to be represented as black nodes are determined by the set gap between nodes. If the value is 50, the consecutive black nodes would have 50 bins in between them i.e. bin 1 is connected to bin 52 by a grey arrow edge; bin 52 is connected to bin 103 and so on. The length of the edge is scaled by this gap argument; however, since edges behave like a spring, this length is only the value at rest, and it can change when the edge must stretch due to the positioning of contacts. Consequently, the network representation of the persistent substructure may not be proportional to the length of the chromosome, hence the equidistant black nodes become the only distance markers. There are also the orange nodes and edges, which represent the highly persistent contacts. Note that when a contact bin coincides with a black marker node, that node is coloured orange. But this contact region can still be distinguished as a distance marker because, unlike the other contact regions located in between black marker nodes, it is preceded by the arrowhead of an edge.

#### Identification of CEll-Type Invariant (CETI) hubs

CETI hubs are comprised of a central region, 40 kb long in this work, highly interconnected with several other 40-kb regions. Per chromosome, hub centres were chosen based on having extremely long-range persistent contacts and then all the 2-Mb persistent contacts (c_p_ ≥ 19) it participates in were retrieved, forming the hub. We have identified 38 such CETI hubs, with each chromosome contributing at least one hub except for chr. 15, chr. 17 and chr. X. For these chromosomes, it was hard to identify hubs because they barely contain contacts between very distant regions (as filtered in our visualisations) compared to other chromosomes of similar length. **Supplementary File S5** brings detailed information on the 38 CETI hubs.

### Feature association with contact persistence

See **Supplementary Table S7** for the comprehensive list of the sources of around 300 features used for the associations. Unless indicated otherwise, ranges overlapping means having at least 1 shared or common bp.

#### Region-wise association

Chromatin and genomic features, which are characteristics of a region (not by a contact), were associated with the unique contact regions per c_p_ *via* two ways. **1)** The significance of feature enrichment or depletion in a set of unique contact regions was quantified through permutation tests (10,000 iterations) using custom wrapper functions based on the R library regioneR [34]. The unique contact regions were the ones being permuted and the random samples were drawn without replacement from the background set. The association was quantified by calculating **(a)** number of contact regions overlapping with feature ranges, where an overlap of one contact region with multiple regions of a feature was counted only once, and **(b)** total intersection in bp. The same permutation test procedure was used for calculating the significance of enrichment of long genes at unique high-c_p_ contact regions except that the sample statistic was the mean length of genes overlapping. Enrichment and depletion of features were determined at unique 40-kb regions forming the c_p_ = 21 and c_p_ ≥ 19 contacts. The background was the set of all unique long-range contact regions (c_p_ ≥ 1). **2)** The number of unique contact regions per c_p_ that overlap with a feature was calculated using custom R scripts.

#### Contact-wise association

The contact-wise association was done *via* two ways. 1) Per c_p_, we determined the fraction of feature-defined contact types based on whether the two regions of a contact overlap or do not overlap with a feature. 2) Per feature-defined contact type, we calculated the fraction of contacts of given c_p_ (see **Supplementary Fig. S10B,11B**).

### Gene-related analyses

In all analyses involving genes, those with multiple transcripts were represented either by the single longest transcript or, in the case of ties, by the first of the longest transcripts (but preferring coding over non-coding ones). The latter was the case for 962 genes out of 24,910 (∼3.86%) unique genes from the UCSC hg19 annotation table.

#### Expression analysis

The Genotype-Tissue Expression (GTEx) data (E-MTAB-5214) [35] (in TPM) was filtered to contain only genes with expression values in at least 1 tissue. The EMBL-EBI Expression Atlas definition of the expression levels was used in this study – not-expressed (NE): TPM/FPKM < 0.5; low-expressed (LE): TPM/FPKM = [0.5,10]; medium-expressed (ME): TPM/FPKM = (10,1000], high-expressed (HE): TPM/FPKM > 1000 [36]. Another baseline expression dataset (E-MTAB-1733) [37], containing RNA-seq data of coding genes from 27 normal tissues from 95 adult individuals, was subjected to the same pre-processing and used to repeat the analyses (**Supplementary Fig. S15 Set 1**).

#### Functional term enrichment analysis with DAVID

Because DAVID can only be used to process up to 3000 inputs at a single instance, 3 sets containing 2999 genes (in some instances, DAVID maps more than 1 DAVID gene identifier to a gene name) were randomly sampled without replacement from 4209 unique genes co-localising with prime contact (c_p_ = 21) regions. The built-in medium stringency of DAVID functional annotation clustering was applied. The built-in set of Homo sapiens genes was used as background. The top 2 enriched clusters were similar across samples and (**Fig. 3D**) shows results from one sample. Gene count is the number of c_p_ = 21 contact genes associated with each term.

### Replication timing data processing and analysis

The replication timing (RT) data (192 samples) from ReplicationDomain [38] were processed using custom R scripts into a final dataset, wherein a 40-kb region has one average RT value from each of the 61 unique cell lines (50 non-cancer- and 11 cancer-related cell lines). RT measurements were binned at 40 kb to match the contact data and normalised across samples by aligning each sample to a reference set by linear-model fitting. For the association with c_p_, the mean and median of the average values from each group of cell lines were calculated for each 40-kb region. Only regions with data from ≥ 59 cell lines were considered. A region was represented by the mean or median of the RT measurements overlapping with it. The consensus RT for a contact was then calculated as the mean of the two means or two medians from the two contacting regions.

### Somatic cancer SNV data processing and analyses

The dataset (N = 38,428,969) was downloaded from the ICGC Data Portal (Release 28, 2019 March 27) [39] from all human autosomes from 2320 samples. The SNVs were categorised based on their location relative to transcript components according to a hierarchical assignment of SNVs in the following decreasing order of priority, exon > intron > intergenic. To quantify the vulnerability of each contact to SNVs, we first calculated, for each region, the number of mutated sites with at least one mutation (Nmutsite), and the total number of mutations (Nmut). Both metrics were normalised to the number of base pairs that can be mutated depending on the SNV type and location (Nmutsite_norm_ and Nmut_norm_, respectively). The consensus value for a contact was then equal to the mean of the metric values from the two contacting regions.

### Contact sequence complementarity calculation

Sequence complementarity of contacts (c_||_) was estimated using three ways. **1)** Calculating c_||_ *via* global sequence alignment using edit or Levenshtein distance (i.e., minimum number of single-character edits to transform one sequence to another) (c_|_|^align^) was implemented using the open-source C/C++ library, edlib [40]. Substitutions, insertions, or deletions were penalised by 1 regardless of the base identity. **2)** Calculating c_||_ *via* matching of 7-mer counts (c_|_|^k-mer^) involved counting the occurrence of all possible 7-mers in both strands of a region and normalising these counts to the length of the region. The c_|_|^k-mer^ of a contact is then calculated as the sum of the absolute differences between the 7-mer normalised counts of pair of regions in contact. **3)** Calculating c_||_ *via* crude estimation of hybridisation energy of regions in contacts (c_||_^G^) was possible with published free energy parameters for unique, perfectly matched DNA triplets [41]. Free energy parameters for 7-mers were derived from the triplet energy parameters by sliding a 3-bp window along the 7-mers and averaging the parametric values of triplets present. The c_||_^G^ of a contact is then given by the sum of the products of the 7-mer parametric values with the matching 7-mer counts of regions in contact. All complementarity values were normalised to the length of contact regions. The c_||_^k-mer^ and c_||_^align^ were negated to be directly proportional to complementarity. See **Supplementary File S1: Section 5.2** for more details.

### Shuffling of contact regions

Shuffling, performed per chromosome and per c_p_, was done using our general-purpose optimisation library, rOptimus [42], in order to maximise the number of fake/shuffled long-range contacts that would not be present in the real/original set (duplicates also not allowed). See **Supplementary File S1: Section 5.3** for more details.

### Repeat-related analyses

Transposon subfamily sites and copy number ranking were derived from the UCSC hg19 RepeatMasker annotation table. For calculating sequence complementarity using repeat-masked genome, only c_||_^k-mer^ was calculated as detailed above. Contacts formed by regions with > 50% of their sequence masked were excluded. In addition, those involving regions with at least one missing bp in the unmasked genome were excluded also to be consistent with the analysis using the unmasked genome. See **Supplementary Fig. S28** caption for more details.

### Contact map generation based on c_p_, c_f_ and c_||_

The Hi-C_p_ maps include all long-range contacts from all cell types. The other Hi-C_f_ data not part of the main contact dataset were downloaded as .hic from the 4DN portal [43] and the sparse contact matrices were retrieved using the R library strawr. For all contact maps with gradient colouring, the upper and lower limits of the colour scale are the upper and lower whisker values (Q1 − 1.5×IQR and Q3 + 1.5×IQR). Values outside these limits are coloured using the corresponding most extreme colour.

## Results

### Contact persistence to focus on core genomic contacts

We quantified the persistence of genomic contacts across human cell types (**Supplementary File S1**, **Supplementary Table S1**) by integrating long-range contacts (40-kb resolution) with gap between the contacting regions ≥ 2 Mb. The latter threshold was taken to be greater than the size of most TADs [44] (**Supplementary Fig. S1**). This was done considering our focus on core, sequence-driven components of genome organisation, which would benefit from minimising the effect of specific mechanisms, such as the reported role in TAD formation of the CTCF/cohesin-mediated loop extrusion predominantly demonstrated at the submegabase scale [17,45,46] and the similarly, more pronounced driver role of cell-type specific transcription at shorter-scale organisation [8]. The assembly of such contacts was possible by using 21 high-quality, re-analysed Hi-C datasets from varied primary tissues and cell lines generously made available by Ren and co-workers [22] (**Fig. 1A**). With the integration, the original contact matrices, denoted as Hi-C_f_, based on the conventional contact frequency measure, c_f_, were converted into a single map - here termed as Hi-C_p_ (**Fig. 1B**). In a Hi-C_p_ map, each contact is represented by a persistence score (c_p_) from 1 to 21, equal to the number of human cell types it is present in (**Fig 1B,C**). The value of c_p_ is independent of the exact c_f_ value that a given contact is present in the different cell types i.e. a contact in a given cell type contributes to c_p_ as long as cf (HiCNorm-normalised [32]) > 0 for uniquely mapped contacts. We did not apply statistical tests to enrich for interactions that have higher c_f_ than the expected value based on distance or gap between contacting regions, as our goal was not to prioritise the potential for the most functionally relevant contacts. It should also be emphasised that contacts not flagged as “significant” by statistical enrichment tests are not guaranteed to be non-contacts or noise [47]. By design, we wanted to work with all contacts with firm evidence of happening, i.e. a robust chimeric read representing that contact. For this reason, we also ensured that potential artefactual/ambiguously mapping reads were not accounted at all. In addition, working with longer-range contacts (> 2 Mb, greater than the average TAD size determined using various methods [44]) was a way to mitigate inclusion of significant noise from the method, e.g. getting chimeric reads for contacts that are just extremely close to each other linearly. Furthermore, if a given contact was just deceptively passing all our checks being a mere artefact, we then had the voting of all the considered cell types to define the persistence of the contact, hence an artefact will not likely have a high contact persistence (c_p_), hence influence our conclusions (**Supplementary Methods** for extended explanation). The contact persistence scores were also found to be robust against any outlying datasets based on sequencing depth and source similarity (**Supplementary Fig. S2**).

**Figure 1.**
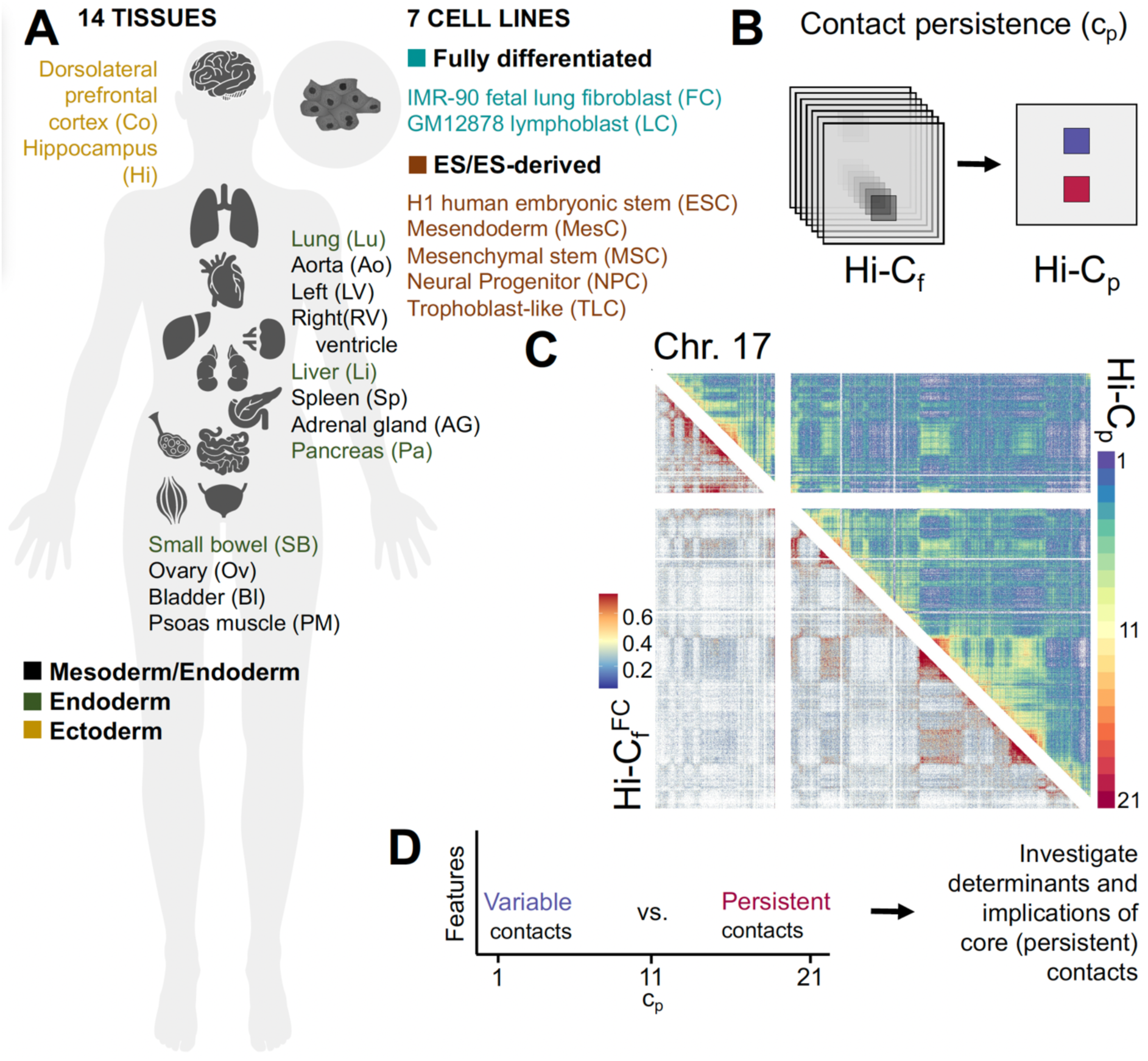
Contact persistence to isolate and investigate the core genomic contacts, their determinants and implications. (**A**) Human tissue and cell line sources of the Hi-C (or Hi-C_f_) datasets [22] used in this study. The icons indicate the sources, and the colours denote tissue origin, cell line source and differentiation state (**Supplementary Table S1**). (**B**) Diagram representation of c_p_ derivation. Hi-C_f_ contacts all in black to indicate that all contacts with HiCNorm c_f_ > 0 (uniquely mapped reads) were considered present in a dataset. (**C**) Chr. 17 FC (IMR-90) Hi-C_f_ compared with Hi-C_p_ of long-range contacts from all cell types. (**D**) The strategy of this study to reveal core determinants of genomic contacts – investigating core, persistent contacts by contrasting with variable ones to reveal their implications and determinants, which are likely to be similarly persistent across cell types i.e. sequence determinants.

Out of the 112,861,349 long-range contacts examined, the proportion of contacts decreases with increasing persistence, with the least and most persistent ones comprising 17.448 % and 0.012 %, respectively (**Supplementary Table S2, Supplementary Fig. S3**). The c_p_ stratification is independent of the exact c_f_, but we did find that persistent contacts tend to have relatively high c_f_ including when accounting for contact gap variation across c_p_ (**Supplementary Fig. S4,5**). Persistent contacts tend to be shorter in range, with median contact gap lengths between 2.4 to 2.7 Mb for c_p_ ≥ 19 contacts (**Supplementary Fig. S6**). Interestingly, in Yang et al. [48], authors reported that short-range contacts, with contact gaps of around 2.5 Mb, primarily have high relative contact frequencies conserved across the lymphoblastoid cells of humans and 3 other primates - chimpanzee, bonobo and gorilla. By directly associating their data with our c_p_ data, we did find that our persistent contacts mostly correspond to the contacts they found to have conserved high-c_f_ pattern (**Supplementary Fig. S7,8**). By contrasting variable and persistent contacts, we investigated the foundations of contact persistence to facilitate the study of the core, invariant determinants (as well as its implications) of higher-order genome organisation (**Fig. 1D**).

In **Fig. 2**, we visualise the persistent contacts as arcs in an arc diagram (**Fig. 2A**) and as edges in a network diagram (**Fig. 2B**), in particular, to highlight outlying persistent contacts that are extremely long-range. For instance, out of its 10,253 c_p_ ≥ 19 contacts, chr. 1 has 303, 90 and 5 highly persistent contacts joining intervals linearly separated by ≥ 8, 26 and 200 Mb distance. The network diagrams also effectively show how high persistence contacts can bring together far-off regions and form clusters of highly interconnected regions that are present amongst most cell types and that we call in this text as CETI (**ce**ll-**t**ype-**i**nvariant) hubs (**Fig. 2B,D**).

**Figure 2.**
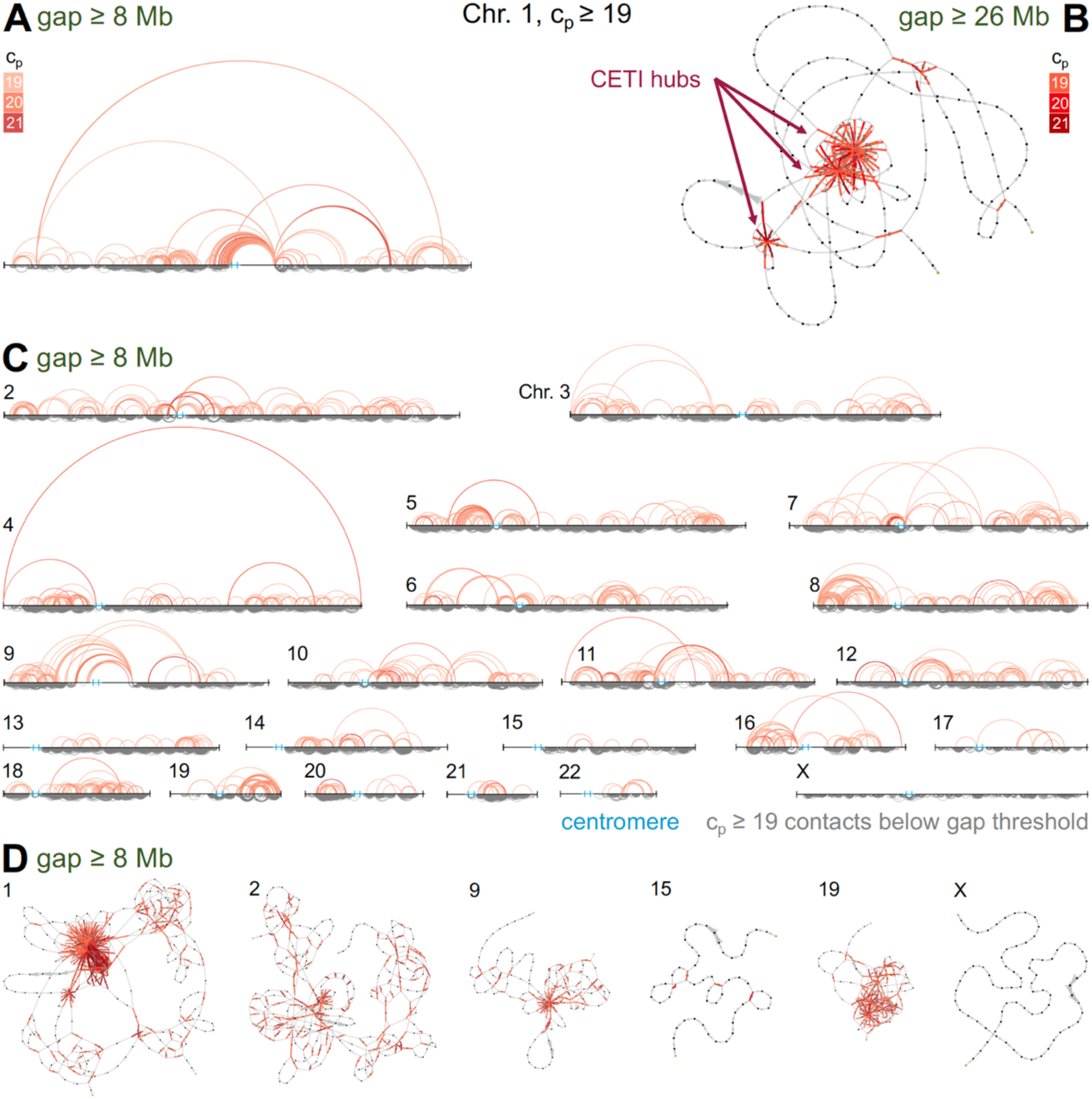
Visualisation of persistent contacts and the chromosome organisation they mediate. Contacts displayed have c_p_ ≥ 19 (c_p_ denoted by arc or edge colour). Centromeric regions are marked by cyan lines and by thick edges on arc and network diagrams, respectively. (**A**) Arc diagram for chr. 1. The upper side has contacts with gap ≥ 200 40-kb bins (or 8 Mb), while the bottom side shows the rest of c_p_ ≥ 19 contacts (≥ 2 Mb). (**B**) Network diagram for chr. 1 formed by c_p_ ≥ 19 contacts with gap ≥ 650 40-kb bins (or 26 Mb). The **ce**ll-**t**ype-**i**nvariant (CETI) hubs of interconnected regions are indicated. (**C**) Arc diagrams corresponding to (A), but for the rest of human autosomes and chr. X. (**D**) Representative network diagrams, analogous to (B), for some chromosomes generated through c_p_ ≥ 19 contacts with gap ≥ 200 40-kb bins (or 8 Mb).

### Feature associations of persistent contacts

To examine the identity, causes and implications of a contact persistent in many cell identities, we looked for significant associations with ∼300 chromatin and genomic features with c_p_ (**Supplementary Fig. S9-11**). We found, at high-c_p_ regions (region-wise), a significant enrichment of heterochromatin-related domains and features i.e. B-compartments and subcompartments, lamina-associated domain (LAD) marker LMNB1 and repressive chromatin mark H3K9me3 (**Fig. 3A** left). At the sequence level, AT-rich features are consistently enriched, particularly A-phased repeats, L1 and L2 isochores, and CpG-depleted prairie sequences [49] (**Fig. 3A** left). Analysis of enriched 7-mers at persistent contact regions show no strong motif, but they tend to contain more A/T over G/C bases (**Supplementary Table S3, Fig. S12, 13**); although, at 40-kb resolution, the differences in the AT content across c_p_ is not drastic (**Supplementary Fig. S14**), with values across c_p_ being close to the recently calculated genome-wide average of 40.9% GC [50]). The enrichment of features is accompanied by depletion of euchromatin-related domains and GC-rich features namely A-compartments and subcompartments, CpG island, and putative G-quadruplex sequences, H1, H2 and H3 isochores, and CpG-dense forest sequences [49] (**Fig. 3A** right). Findings translate contact-wise, with the proportion of contacts associated with the enriched features increasing with c_p_ as demonstrated here for H/L isochores (**Fig. 3B**).

**Figure 3.**
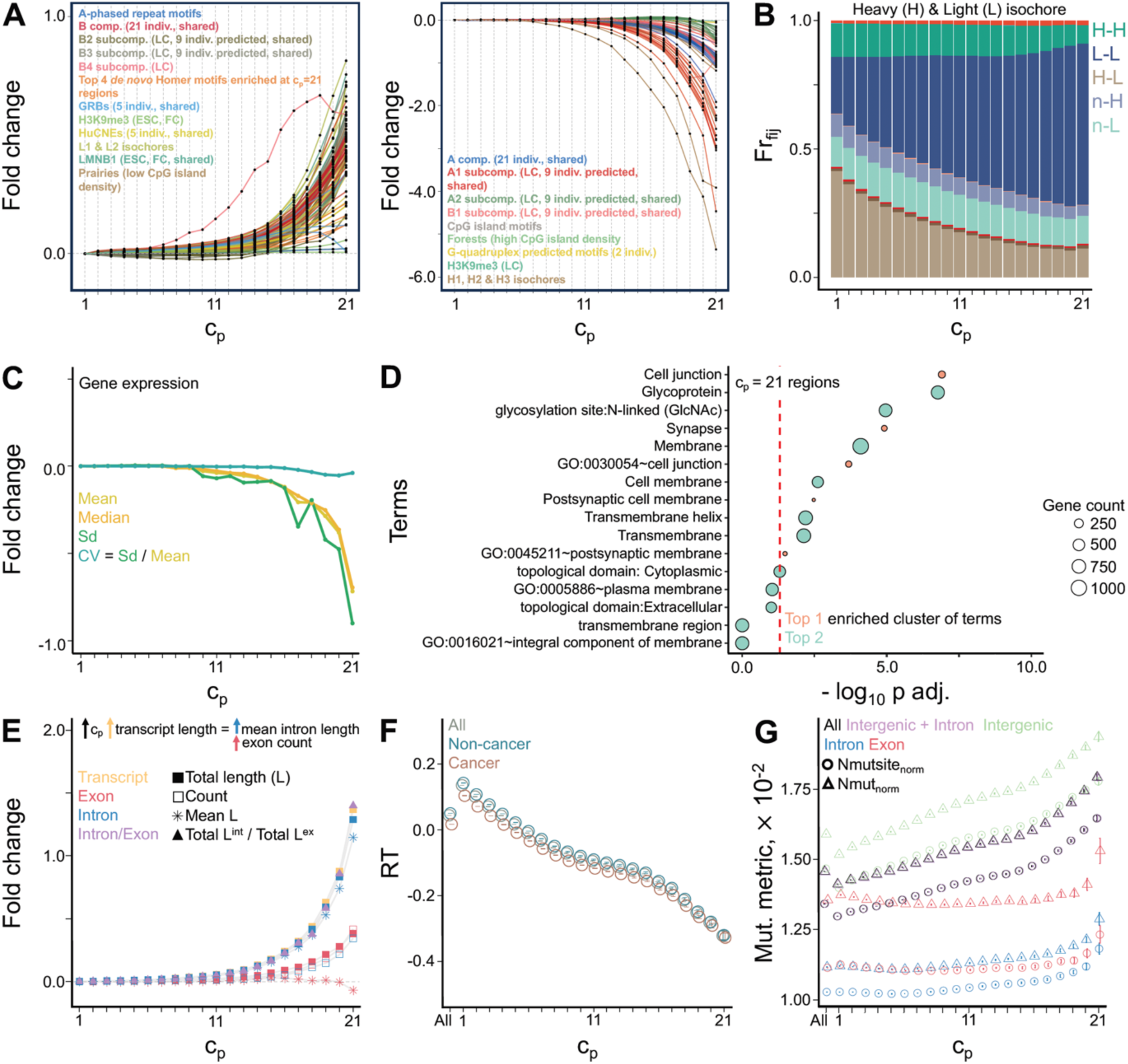
Persistent contacts enriched for contacts with features associated with heterochromatin and preferential AT sequence composition. The log_2_ fold changes are relative to value at c_p_ = 1. The c_p_ = “All” refers to all long-range contacts. (**A**) Fold change of the proportion of unique contact regions across c_p_ overlapping with significantly (left) enriched and (right) depleted features at unique c_p_ = 21 contact regions (p-value < 0.05, permutation test, see **Supplementary Fig. S9** heatmap for complete result of permutation tests and **Supplementary File S2** for data behind the heatmap). Cell-type specific features can have multiple datasets (indicated in parentheses). “Indiv.” refers to individual data, “shared” refers to regions shared by or common to all individual data for that feature, and “predicted” refers to subcompartment regions predicted by SNIPER [53]. (**B**) Fraction of contacts (Fr_fij_) overlapping isochore families across c_p_ (see **Supplementary Fig. S10,11** for similar plots of other enriched features). Only dominant contact types are shown in legend; “n” means no overlap. (**C**) Fold change of the mean of various cross-tissue expression metrics across c_p_ (**Supplementary Fig. S15,16**). The c_p_ ≤ 3 and c_p_ ≥ 19 distributions are significantly different (p-values < 0.002) except for CV (Mann-Whitney-Wilcoxon, MWW, p-value of 0.105). Only genes with data in ≥ 70% of the tissues were considered. (**D**) Top 2 most significant DAVID clusters of enriched functional terms at c_p_ = 21 (**Supplementary Fig. S17**). Red dashed line at log_10_ 0.05 ∼ 1.301. (**E**) Fold change of the mean of derivative lengths and counts of various genic elements across c_p_ (**Supplementary Fig. S18**). Mean length of c_p_ = 21 and c_p_ ≥ 19 genes significantly higher than c_p_ ≥ 1 genes (p-value < 0.0001, permutation test). (**F**) Mean cross-tissue replication timing (log_2_ ratio of early and late signals) across c_p_; error bars at 95% confidence intervals (**Supplementary Fig. S19**). All pairwise comparisons of the distributions are significantly different (p adj. < 0.002). (**G**) Mean somatic cancer single nucleotide variant (SNV) frequency metrics across c_p_ (**Supplementary Fig. S20-22**); error bars at 95% confidence intervals. Nmutsite_norm_ and Nmut_norm_ are the number of mutated sites with at least one mutation, and the total number of mutations, respectively, normalised to the total bp that can be mutated depending on the SNV type and location.

Consistent with enrichment of heterochromatin-related features, we also observed that genes at persistent contacts have a relatively lower expression across tissues -evident in the lower distribution of cross-tissue mean and median values of expression, but with the cross-tissue normalised variation (CV) being not significantly different between variable and persistent gene sets (**Fig. 3C**). Consistently, the fraction of tissues with low expression for a gene increases with c_p_, accompanied by a decrease in fraction of tissues with medium and high expression (**Supplementary Fig. S15**). The c_p_ = 21 contact regions co-localise with genes enriched for terms related to synapse, glycoproteins, and (trans)membrane (**Fig. 3D**). With the most significant cluster of terms showing enrichment of neuronal genes that are known to be long [51] we also investigated gene length variation and found that c_p_ = 21 contact genes are also significantly longer, driven by higher mean intron length and greater count of exons as c_p_ increases (**Fig. 3E**). To further probe the implications of the persistent genome organisation, we associated contact persistence with two associated molecular phenotypes, replication timing (using aggregated data from different cell types) and somatic mutation frequency (using cancer SNV data). Consistent with the enrichment of heterochromatin-related features [52], we found that persistent contacts occur between regions that are late-replicating (**Fig. 3F**) and have higher SNV frequencies (**Fig. 3G**).

### Higher sequence complementarity between persistent contacts

Sequence-specific interactions involving the recognition and association of complementary or homolog nucleic acid sequences are commonplace occurrences in various molecular processes (**Fig. 4A** left). The protein-independent, preferential interaction of identical DNA duplexes (DNA self-assembly) with the help of biological cations has also been demonstrated *in vitro* [54–58], even for DNA occurring in nucleosomes (nucleosome self-assembly) [59] (**Fig. 4A** right, see **Supplementary File S1: Section 5.1** for description of each paper). The sequence identity awareness or recognition involved in these processes, whether happening directly and/or indirectly, prompted us to hypothesise that the degree of sequence similarity or complementarity between regions is associated or could contribute to the tendency of regions to come in contact, regardless of cell type or state. Associating the similarity of sequences with contact persistence, we have taken a general approach, defining the similarity to be independent of any specific chromatin feature or genomic pattern, by using different measures of complementarity between sequences, c_||_. These measures were derived from **1)** matching of short-span 7-mer counts between sequences in contact (c_||_^k-mer^) (**Fig. 4B**), **2)** long-span global sequence alignment using edit distance (c_||_^align^) (**Fig. 4C**), and **3)** approximate estimation of hybridisation free energy (c_||_^G^, decreasing trend means increasing c_||_). Remarkably, for these three metrics, we observe a stepwise increase in the complementarity of sequences with rising persistence (**Fig. 4B,C**). Furthermore, when shuffling the contacting DNA regions within a c_p_ category to form fake contacts not actually paired in that c_p_ category, fake contacts showed significantly lower c_||_ distribution of fake compared to that of real ones, more pronounced for high-c_p_ values (**Fig. 4B,C**). This suggested that the degree of complementarity is specific to real pairs of sequences in contact and not solely dependent on a single sequence motif or pattern present in all persistent regions. The observation holds valid even when limiting the drastic contact gap variation across c_p_ by observing c_||_ trends at narrower ranges of contact gaps (**Fig. 4D**), and when completely removing the effect of gap or distance (**Supplementary Fig. S23**), which may independently impact contact formation.

**Figure 4.**
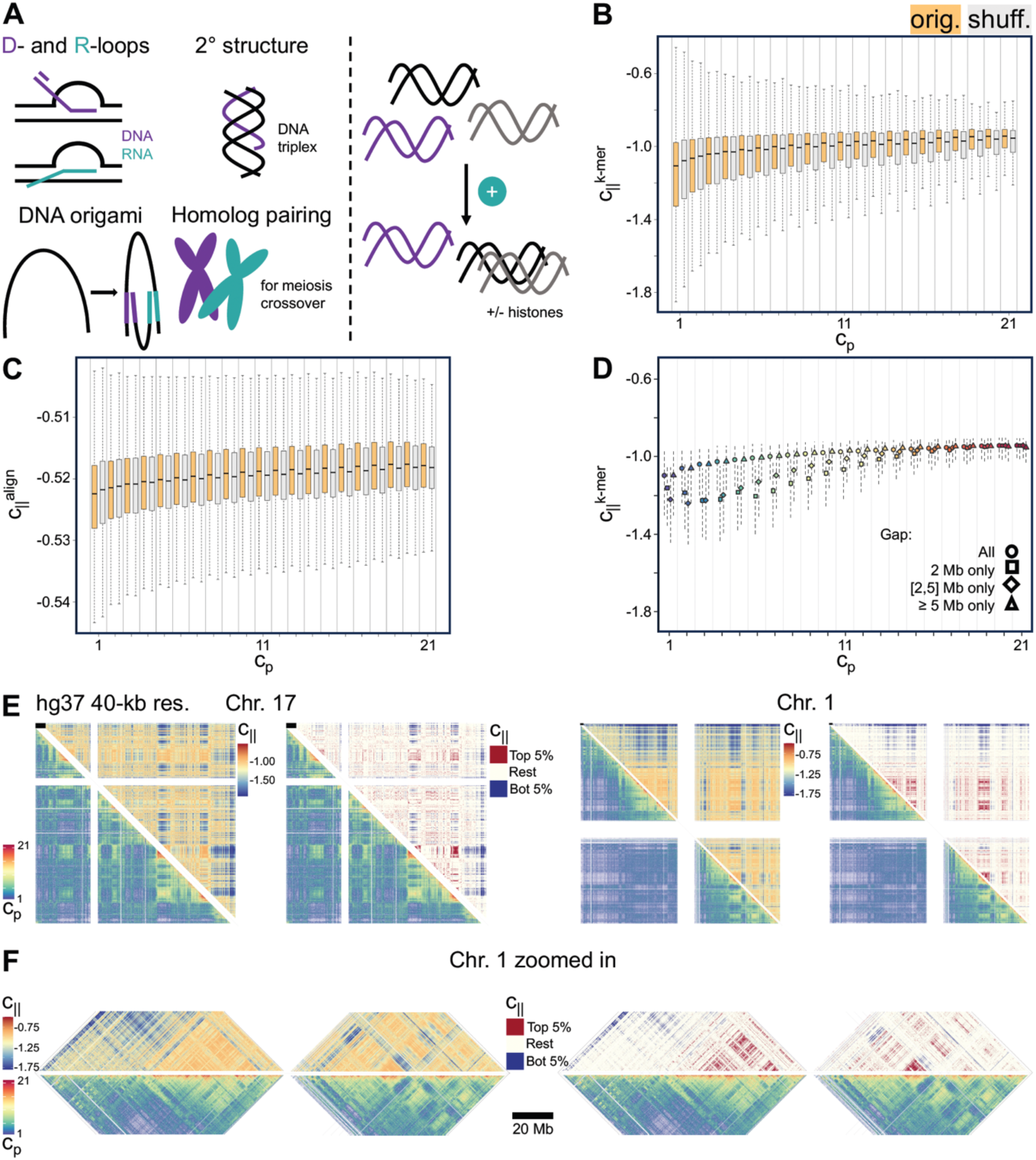
Higher sequence complementarity between persistent contacts. (**A**) Studies on the preferential association of identical DNA duplexes *in vitro* [54–59] (right) motivate the implication of sequence complementarity, c_||_, in contact formation, reinforced by many known sequence-dependent processes (left), involving single- and double-stranded nucleic acids (e.g. D-[60] and R-loop [61,62] formation, triplex formation [63,64], DNA origami nanotechnology [65] and homolog pairing [66]). Sequence complementarity of original (orig.) contacts calculated based on (**B**) short-span k-mer matching (c_||_^k-mer^) and (**C**) long-span global alignment using edit distance (c_||_^align^); c_||_ of shuffled (shuff.) contacts in grey. Pairwise comparisons of orig. distributions between any two neighbouring c_p_ values show significant differences (p adj. < 0.01) except for some comparisons between the highly persistent contacts (c_p_ ≥ 17). Each orig. vs. shuff. distribution pair also significantly different (p adj. < 0.001). (**Supplementary Fig. S24-27**, **Supplementary Table S4**) (**D**) c ^k-mer^ across c_p_ at specific contact gap ranges (**Supplementary Fig. S28**). Shown are medians of distributions and the dashed segments extend to the 25th and 75th percentiles. Pairwise comparisons of the 5 most persistent and 5 most variable distributions show significant differences (p adj. < 0.05). (**E**) Hi-C_p_ vs. Hi-C ^k-mer^ using absolute and binned values of c_||_ (top and bottom 5% contacts based on c_||_ values in red and blue, respectively; the rest in beige (see **Supplementary File S4** for other chromosomes). Braces highlight some areas similar between contact maps. Black scale bars at the top left corner are 4 Mb long. (**F**) Zoom in on chr. 1 highlighted regions in (**E**).

We thus found a link between the Hi-C-inferred contact persistence and a simple metric, c_||_, calculable between any two sequences, without any training or assumption borrowed from experimental data. Consistent with the observed positive correlation between c_||_ and c_p_, the complementarity-based map, Hi-C_||_ (generated by directly plotting the calculated c_||_ values and binned at a matching resolution), shows areas of both high and low signal like those in the Hi-C_p_ (**Fig. 4E,F**, highlighted with braces). Consequently, similar areas of matching high and low signals can also be seen when comparing c_||_ with c_f_ data from human embryonic and lymphoblastoid cell lines (**Fig. 5A,B**), and *Drosophila* embryonic and differentiated (neuronal) cell types (**Fig. 5C**) at different resolutions (**Supplementary File S3** for complete set of contacts maps across chromosomes using data from different human and *Drosophila* cell lines). These similarities are emphasised in the versions of the hybrid contact maps, where c_||_ is binned to distinguish the top and bottom 5% contacts based on value.

**Figure 5.**
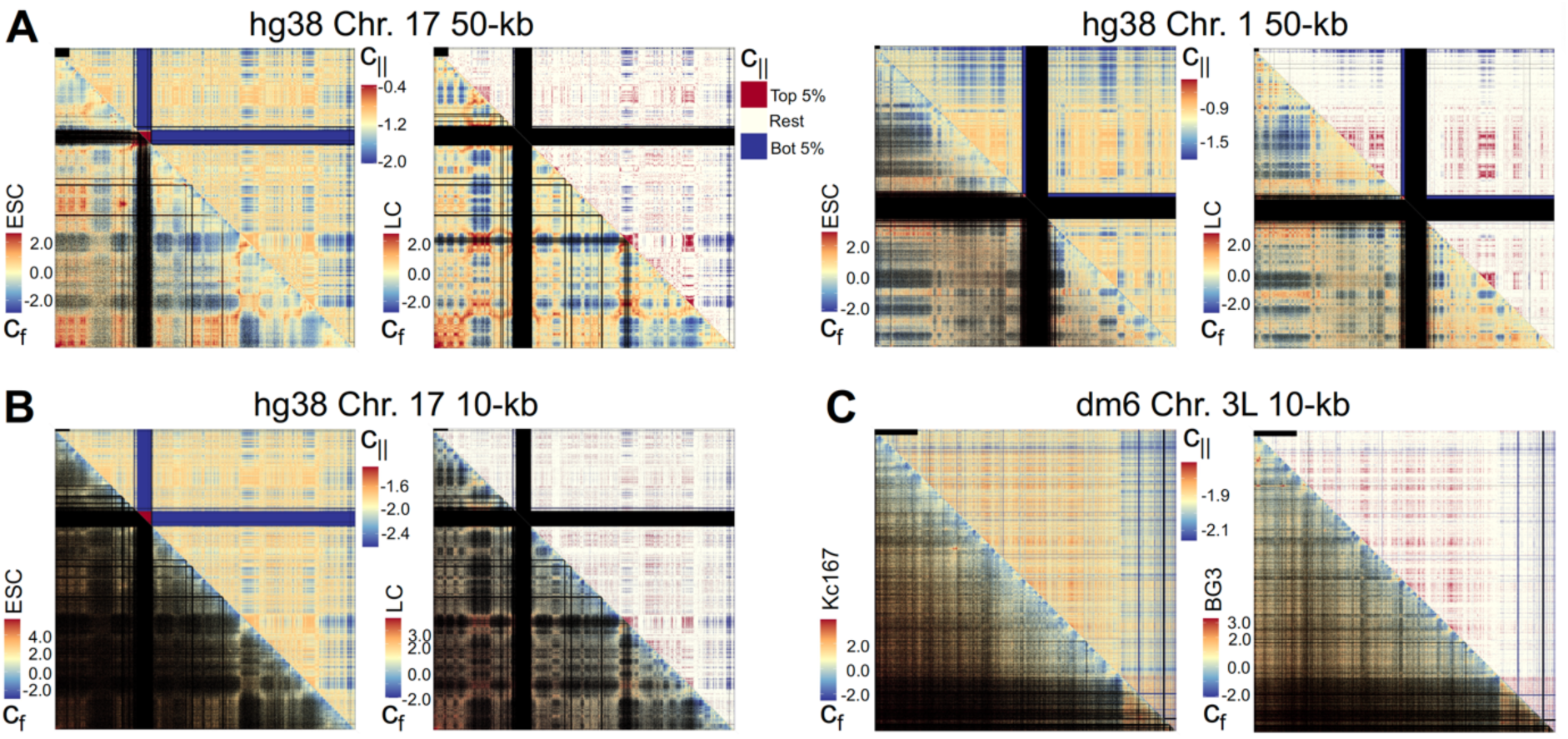
Genome-wide c_II_ values recapitulate some Hi-C features shown for Human and *Drosophila*. Hi-C_f_ vs. Hi-C ^k-mer^ using log_2_ observed over expected c_f_, and absolute and binned values of c (top and bottom 5% contacts based on c_||_ values in red and blue, respectively; the rest in beige. Black scale bars at the top left corner are 4 Mb long. (**A-B**) Human (hg38) cell types: ESC (H1-hESC) embryonic and LC (GM12878) lymphoblastoid cells; (**C**) *Drosophila* (dm6) cell types: Kc167 embryonic and BG3 neuronal cells. See **Supplementary File S3** for other chromosomes.

### Repeat in the observed sequence complementarity of contacts

Based on the DNA/nucleosome self-assembly phenomenon [54–59], repetitive elements, representing similar sequences, have been proposed to mediate a sequence-dependent phase separation of the genome [67]. Consistently, a combination of computational [68–70] and experimental [71] findings show that the self-clustering of repeat families is indeed correlated with genome organisation. Given these, we determined whether transposons, which have brought similar sequences to different parts of the genome, could be reinforcers of the observed higher complementarity at persistent contacts by using metrics to measure site distribution of a subfamily between two contacting regions. In **Fig. 6**, we highlight 4 candidate subfamilies, MIRb, MIR, L2a and L2c, likely to be most influential to the observed higher complementarity of persistent contacts (green text in **Fig. 6**). These subfamilies, among the ones with the highest copy numbers, have at least 1 shared number of sites in more than 50% of persistent contacts (c_p_ ≥ 19) (**Fig. 6A**), and higher median site skew at persistent compared with variable contacts (c_p_ ≤ 3) (**Fig. 6B**). Based on the site distribution metrics, their sites tend to be more distributed between persistent contact regions even relative to other high-copy-number subfamilies like AluJb and AluSx (∼2000 sites more than L2c). MIRb was the only subfamily that had higher median shared number at persistent contacts (**Fig. 6C**). MIRb does have the highest copy number among the subfamilies, but it is not the only one that could have 2 shared sites because the other 3 subfamilies had median site total of ≥ 4 sites (**Fig. 6D**) at persistent contacts. MIR, L2a and L2c have mean shared numbers at persistent contacts significantly higher than the variable counterparts. As for the rest of the subfamilies, since only contacts with at least 1 site could be considered in the site skew calculation, having a median site skew of 1 means that most of the (long-range, intra-chromosomal) contacts these subfamilies have inserted on have no shared site.

**Figure 6.**
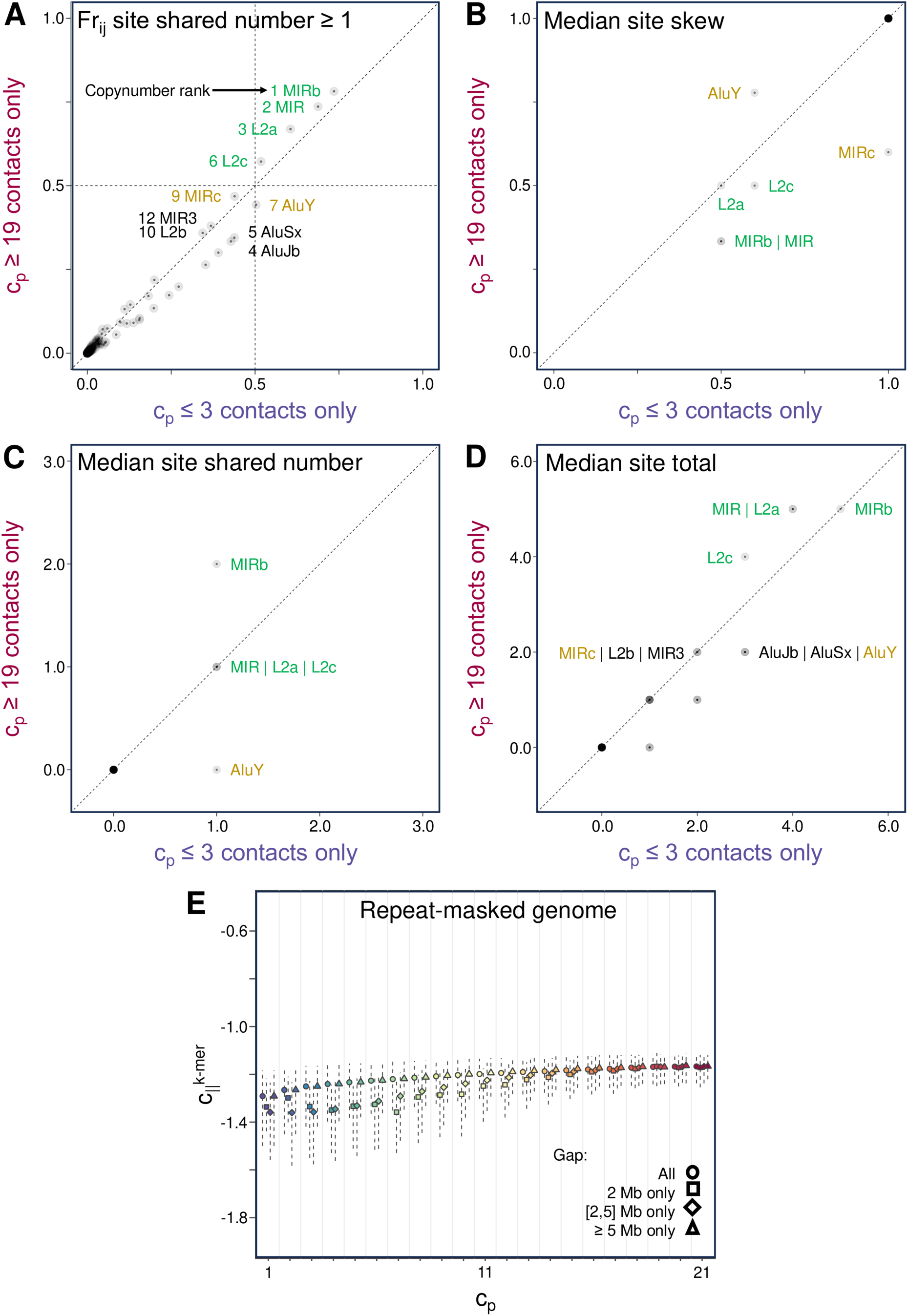
Repeat contribution to the observed sequence complementarity of contacts. Transposon subfamily site distribution at persistent (c_p_ ≥ 19) vs. variable contacts (c_p_ ≤ 3). Given *I* and *j* forming a contact, and *N_i_* and *N_j_* being the number of subfamily sites in *i* and *j*, **A** shows fraction of contacts with at least 1 shared number of site i.e. min(*N_i_*, *N_j_*) = 1. In green text are subfamilies with ≥ 1 shared number at more than 50% of contacts. **B-D** show medians of the distributions at persistent and variable contacts of the following metrics: **B** site skew equal to |N_i_ – N_j_| / (N_i_ + N_j_), **C** shared number of sites i.e. min(*N_i_*, *N_j_*), and **D** total number of sites of a contact i.e. *N_i_* + *N_j_*. In yellow text, are additional subfamilies with median site skew not equal to 1 for both distributions. Only contacts with at least 1 site can have a valid site skew value. All subfamilies from the UCSC hg19 RepeatMasker annotation table were used and each data point, representing a subfamily, is coloured with a transparency level of less than 1, whereby darker areas indicate the overlap of many points. For all named subfamilies, the variable and persistent distributions of the metrics are significantly different (p-values < 0.03). **E** c ^k-mer^ across c_p_ at specific contact gap ranges using repeat-masked genome (**Supplementary Fig. S28**).

Given that repeats represent a significant amount of sequence, about half of the human genome, it is not surprising that repeats are key contributors to the observed complementarity phenomenon. Finding that some high-copy-number subfamilies, particularly the ancient MIRb and MIR transposons as well as L2a and L2c, tend to have more shared sites that are more distributed at persistent contacts support this and is consistent with earlier aforementioned studies. It should be noted, however, that the site distribution metrics do not directly measure the individual and collective contribution of subfamilies to the total degree of complementarity of contacts based on the length and sequence of the remnants. Interestingly, when we used the repeat-masked genome to analyse a smaller subset of long-range contacts, whose regions were mostly devoid of annotated repeats, even the unmasked portions of persistent contacts in that subset were found to be more similar than that of variable contacts, suggesting that this higher complementarity at the given resolution is a characteristic of persistent contacts that the annotated simple and interspersed repeat sites could not solely account for.

## Discussion

In this study, we aimed to contribute to the understanding of the sequence basis of 3D genome organisation. A sequence determinant learned from one or a few cell types could potentially be applicable to any other type for being a cell-context-invariant feature. However, because a phenotype or phenomenon, such as regions being in contact, is evidently dictated by the interplay of cell-invariant sequence and cell-specific epigenetic drivers, the significant association of a contact with a specific sequence motif or pattern in one or few cell types may not hold in other cell types. This underscores the advantage of our strategy to isolate and investigate the cell-context-persistent contacts (or any phenotype), in this case cell-type persistent ones, for studying the sequence basis of genomic contacts, as well as the contribution of epigenetic features that behave consistently across cell contexts, explored here through comprehensive associations of various chromatin features using data from different cell types.

We have found that persistent contacts are predominantly associated with B-compartment/heterochromatin features across different cell types and AT-rich sequence features, which indeed are less dictated by the cell identity, i.e. vary less across cell types (particularly, constitutive heterochromatin regions), in contrast to A-compartment active regions. This observation is also linked to the proposed model that B compartmentalisation, which was not linked to any sequence pattern except for preferring AT-rich regions, could be the “default” state of sequences [72], unless sequences become A-compartment material by e.g. having active TSS sites [72], which are prone to cell-type specificity depending on the required gene expression profile. Also, with AT-rich duplexes found to interact more favourably than GC-rich ones *in vitro* [73], AT-rich features enriched at contacts, could contribute to persistence across cell states, demonstrating how a non-specific or core sequence effect can have influence on which regions preferentially interact. Also, this tight clustering of AT-rich sequences in, generally, heterochromatin regions, could account for some chromosomes, e.g. chromosomes 1 and 9, which are known to have a large portion of constitutive heterochromatin, containing prominent CETI hubs i.e. large clusters of persistent contacts particularly extremely long-range ones (**Fig. 2**, **Supplementary File S5**).

The mechanisms involved in the preferential interaction of similar regions, correlated with genome organisation, are yet to be elucidated in detail but have been described in terms of direct and indirect mechanisms. The direct recognition of sequence identity is supported by *in vitro* [55–57,59,73] and *in vivo* (in *Neurospora crassa* fungus) [74] studies. In addition, similar DNA sequences could have similar proteins and RNAs co-localised on them, which could be the ones driving the preferential interaction or phase separation [71,75]. Both mechanisms could potentially contribute, and the combination and identity of factors may vary for certain subsets of contacts depending on whichever sequence or epigenetic features are present. Our analyses do not thoroughly characterise specific combinations of mechanisms at play, but the association studies have identified a general characteristic, sequence complementarity between regions, that is most pronounced for contacts persistent across different cell contexts. Hence, it is a potential core sequence determinant of genome organisation, which contributes to the “default” tendency of interactions between regions in any cell context, much like the observed favourable association of AT-rich duplexes [73]. The implication of sequence complementarity in genome organisation along with findings from the characterisation of the persistent contacts also corroborate with associations previously brought up elsewhere, providing insight on the relationship between genome organisation and function. For instance, as summarised in [76], pairing of homologous DNA duplexes have been proposed to have a role in initiating heterochromatin formation and transcription silencing [77], in mediating cytosine methylation [78], which produces mutation hotspots across the genome, and in generating supercoiling, which could affect topoisomerase-dependent long genes [79].

Findings presented here prompt further computational and experimental investigations. The contribution of sequence to genome organisation could be quantitatively assessed, both at long and shorter scales, by comparing models of genome organisation based on the complementarity of sequences (and other sequence-derived features such as k-mer contents calculated in this study and the recently reported quantum mechanical properties of genomic sequence [80]) with experimental data not only from different cell types but also from other layers e.g. across cell cycle and species. These models can be generated through molecular simulations, as we have attempted (**Supplementary Fig. S29,30**). Quantifying the similarity of regions using the complementarity measures, which goes beyond defining similarity based on a particular feature like repeat site content, could enable dissection of how the degree and pattern of similarity or homology between regions could influence genome organisation, which are relevant based on *in silico* [81], *in vitro* [82] and *in vivo* [74] mechanistic studies of DNA duplex association [76], and phase separation mechanisms [83]. Finally, experimental investigations are crucial, particularly, to validate and disentangle the potential direct and/or indirect contributions of genomic sequence to the 3D organisation.

This study therefore contributes to the understanding of the relationship between genome sequence and structure by implicating a single parameter, sequence complementarity, as a core factor contributing to the formation of genomic contacts. Along with other works that aimed to understand the encoding of the structure into the sequence (reviewed in [9–12]), this study shows that the complementarity between different parts of the genome may play a role in this encoding, and suggests that organising mechanisms, such as phase separation, are not agnostic to the underlying DNA sequence. Consequently, the DNA could be involved in both direct and indirect manner, and we encourage experimental validation that will help delineate the contribution of sequence to genome organisation, in conjunction with earlier-characterised protein and epigenetic determinants.

## Data Availability

The computer code is available through the GitHub repository: https://github.com/SahakyanLab/GenomicContactDynamics.

This study is a purely computational work that relied on these publicly available datasets and software:

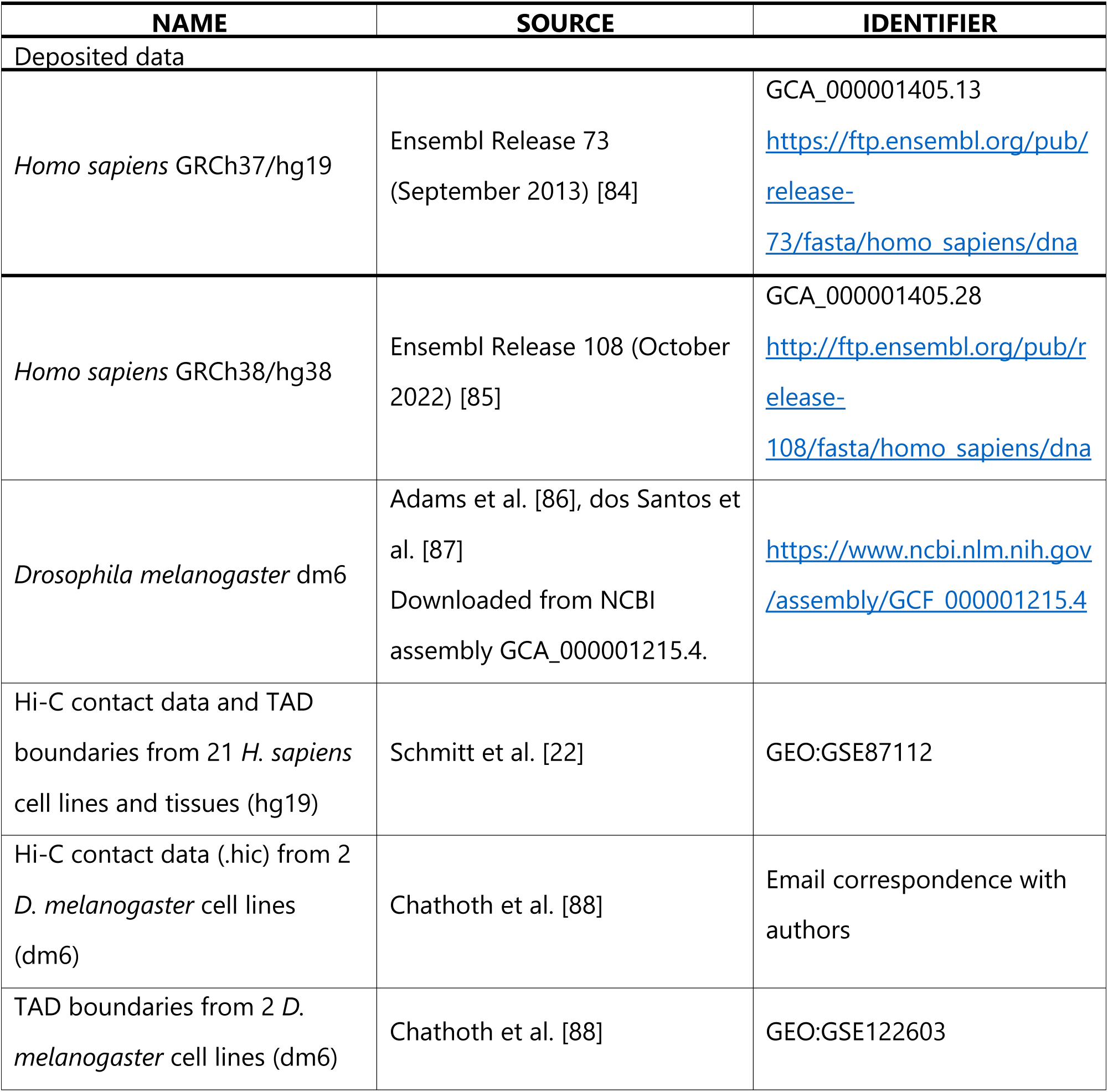
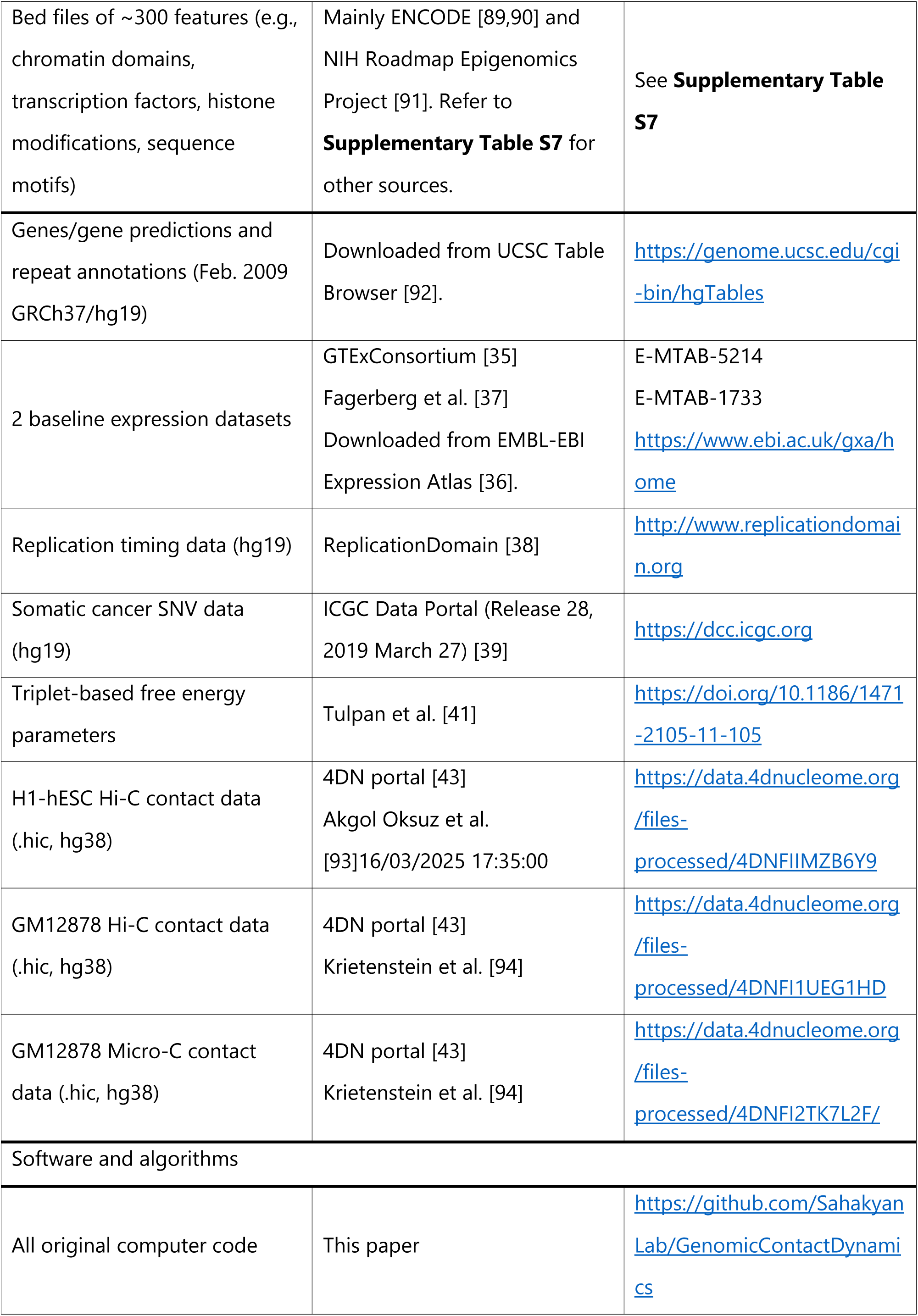
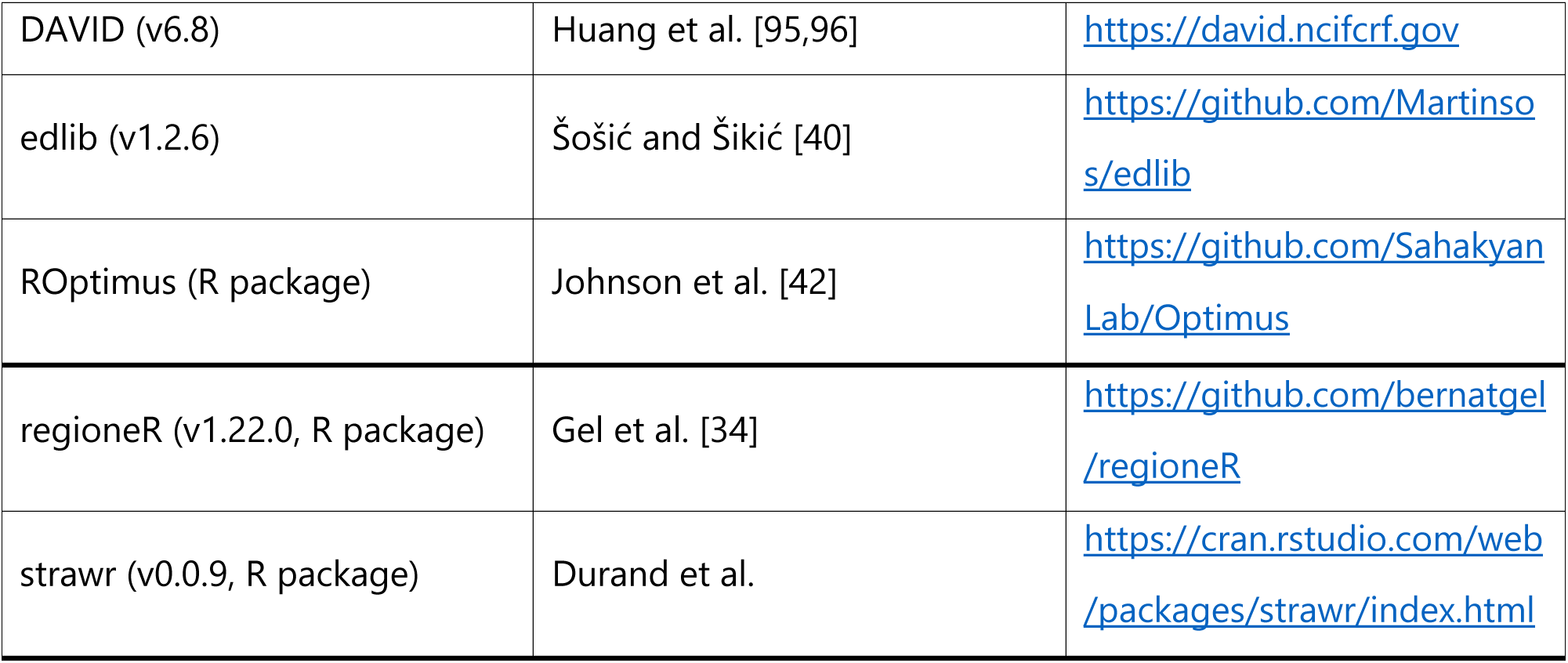

## Supplementary Data

Supplementary Data are available at NAR Genomics & Bioinformatics online.

## Acknowledgements

L.T. is grateful to the Jardine Foundation for supporting her DPhil studies. J.A. is thankful to the MRC for funding his DPhil through a WIMM Studentship. The Sahakyan laboratory including this project has been supported by the UK Medical Research Council (MRC) for the MRC Strategic Alliance Funding (MC_UU_12025). UCSF Chimera, used here for 3D genome model visualisation, was developed by the Resource for Biocomputing, Visualization, and Informatics at the University of California, San Francisco, with support from NIH P41-GM103311. The cell-MDCK icon by DBCLS (grey-scaled in this paper) in Fig 1A of this study is licensed under CC-BY 4.0 Unported and was retrieved from bioicons (https://bioicons.com).

## Author Contributions

Liezel Tamon: Conceptualisation, Formal analysis, Methodology, Software, Validation, Visualisation, Writing – original draft. Aleksandr B. Sahakyan: Conceptualisation, Funding acquisition, Methodology, Resources, Software, Supervision, Writing – original draft. Zahra Fahmi: Formal analysis, Methodology, Writing – review & editing. Rosana Collepardo-Guevara: Funding acquisition, Methodology, Resources, Supervision. James Ashford: Software, Writing – review & editing.

## Funding

This research has been supported by the UK Medical Research Council (MRC) for the MRC Strategic Alliance Funding (MC_UU_12025).

## Conflict of Interest

None declared.

## Notes

https://github.com/SahakyanLab/GenomicContactDynamics

### Competing Interest Statement

The authors have declared no competing interest.

