## Supplementary File S1 for "Analysis of long-range contacts across cell types outlines a core sequence determinant of 3D genome organisation"

### Supplementary Information

8 Appendix 46

9 Bibliography 50

### 3 Contact persistence to focus on core genomic contacts

#### 3.1 Contact Persistence, $c_p$

***Table 1. The 7 cell-line and 14 primary-tissue sources of the main Hi-C dataset used in this study.*** *The Hi-C data were consolidated and processed in Schmitt et al. [1] (GEO:GSE87112). The origin of the cell lines and primary tissues, their abbreviations (Abbr.) as used throughout this work, and the publications that generated the data are listed. Source numbers refer to* ***1)*** *Schmitt et al. [1],* ***2)*** *Leung et al. [2],* ***3)*** *Jin et al. [3],* ***4)*** *Selvaraj et al. [4] and* ***5)*** *Dixon et al. [5].*

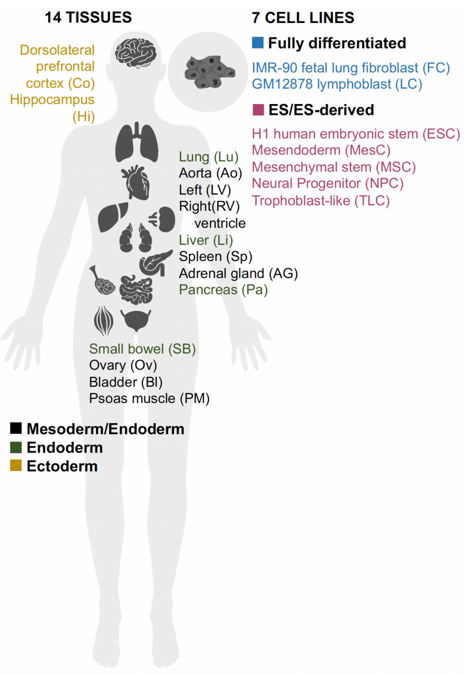

| Type / Origin | Cell line / Primary tissue | Abbr. | Source |
|---|---|---|---|
| Ectoderm | Dorsolateral prefrontal cortex | Co | 1 |
| | Hippocampus | Hi | 1 |
| Endoderm | Liver | Li | 2 |
| | Lung | Lu | 1 |
| | Pancreas | Pa | 1 |
| | Small bowel | SB | 1 |
| Mesoderm / Endoderm | Aorta | Ao | 2 |
| | Adrenal gland | AG | 1 |
| | Bladder | Bl | 1 |
| | Left ventricle | LV | 2 |
| | Ovary | Ov | 1 |
| | Psoas muscle | PM | 1 |
| | Right ventricle | RV | 1 |
| | Spleen | Sp | 1 |
| Fully differentiated | IMR-90 fetal lung fibroblast cell line | FC | 3 |
| | GM12878 lymphoblast cell line | LC | 4 |
| ES/ES-derived | H1 human embryonic stem cell line | ESC | 5 |
| | Mesendoderm cell | MesC | 5 |
| | Mesenchymal stem cell | MSC | 5 |
| | Neural Progenitor cell | NPC | 5 |
| | Trophoblast-like cell | TLC | 5 |

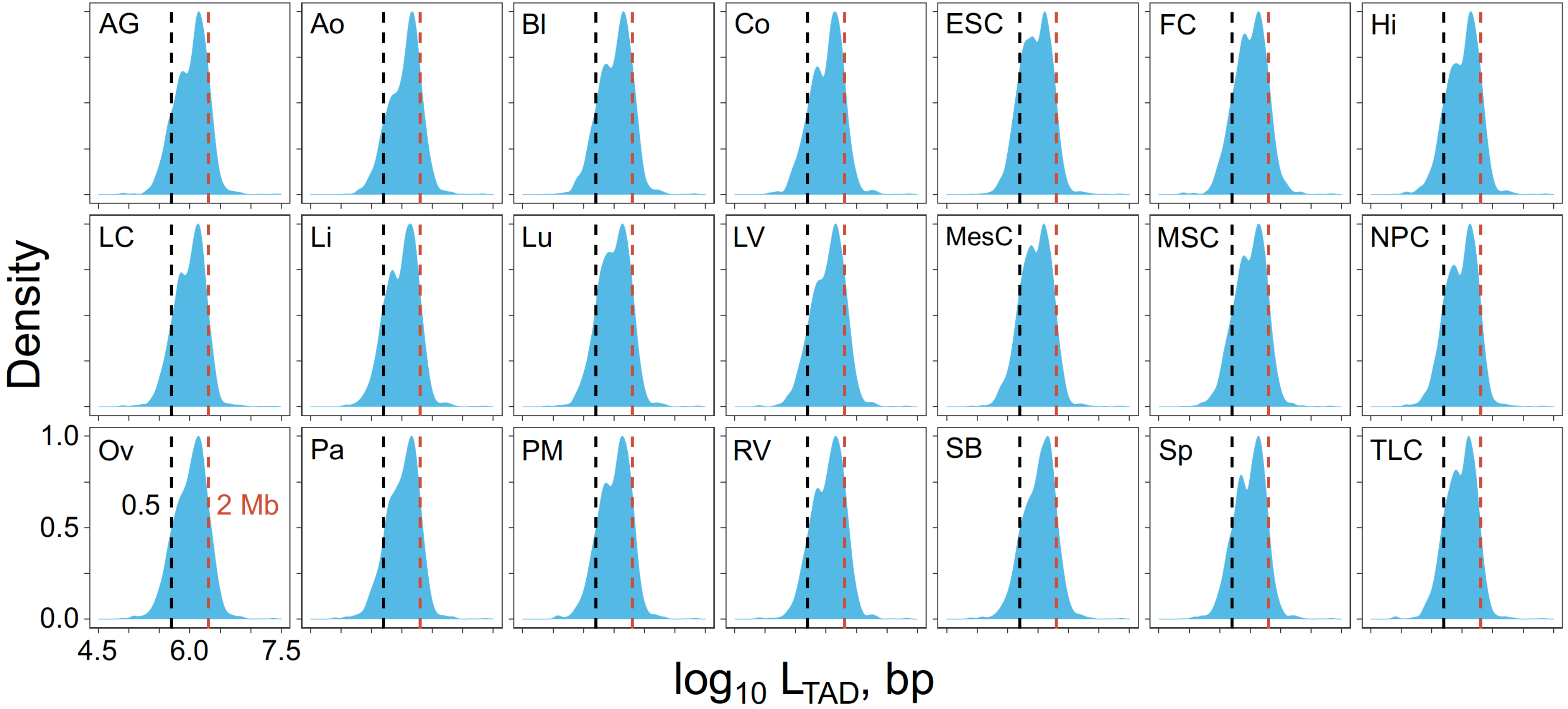

***Figure 1. TAD size distributions in the 21 Hi-C dataset.*** *$\mathrm{Log}_{10}$-transformed TAD sizes were calculated from the TAD boundary coordinates provided in Schmitt et al. [1] as determined via the insulation score method [6] for the same 21 Hi-C dataset. TAD size was defined as the total number of base pairs between the boundaries, excluding any base pair from the 40-kb boundary region. TAD counting started from the most upstream boundary given. Dashed lines mark the two minimum contact gaps, 2 and 0.5 Mb, used for selecting the long-range contacts.*

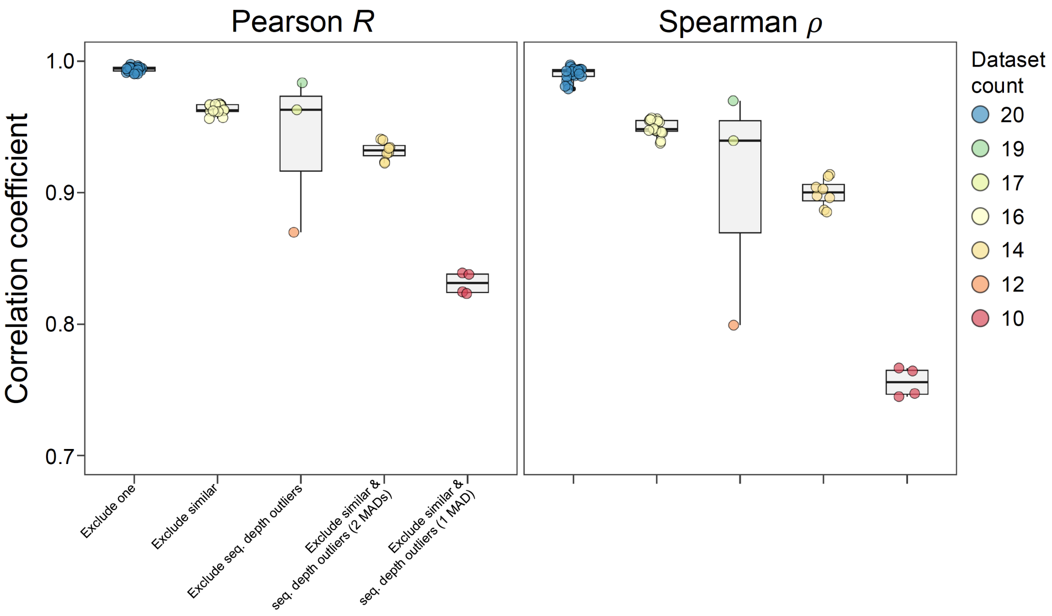

***Figure 2. Correlation of original $c_p$ with recalculated $c_p$ from leaving out contact datasets.*** *Pearson and Spearman coefficients of correlation of original $c_p$ with recalculated $c_p$ after **1)** excluding 1 dataset at a time ("Exclude one"), **2)** excluding 5 datasets at a time ("Exclude similar") to not have multiple datasets from a group of related cell/tissue types as described below, **3)** excluding datasets with outlying sequencing (seq.) depth based on median absolute deviation (MAD) ("Exclude seq. depth outliers"), and excluding similar datasets and outlying datasets based on **4)** 2 MADs ("Exclude similar & seq. depth outliers (2 MADs)") and **5)** 1 MAD ("Exclude similar & seq. depth outliers (1 MAD)"). All correlation tests were significant (Mann-Whitney-Wilcoxon / MWW test, BH $P$ value $<$ 0.0001). The colour of points indicate the count of datasets used to recalculate the $c_p$.*

**Leave-out analysis.** Leave-one-out analysis was performed to test the robustness of contact persistence score to the set of Hi-C datasets. Contact persistence was recalculated after removing 1 dataset from the 21, and then correlated with the original $c_p$ values. This was repeated, removing each of the 21 dataset at a time. Another type of leave-out analysis was done excluding multiple (6) datasets at a time to make sure that no closely related tissue/cell types are included at one time. Leave-out-analysis was performed taking 1 representative cell/tissue type from each group and taking the rest that do not belong in a group (i.e. excluding 6 datasets at a time). The following cell/tissue types were grouped together as nervous tissue = {"Dorsolateral prefrontal cortex", "Hippocampus"}, muscle tissue = {"Left ventricle", "Right ventricle", "Psoas muscle"}, and multipotent precursors = {"Mesendoderm cell", "Mesenchymal stem cell", "Neural Progenitor cell"}. We repeated this for each possible combination of representatives per group. Before correlation, the recalculated and original $c_p$ values are scaled by the total number of datasets. Correlation test was done as described in **Supplementary Methods**.

**Identification of outlier datasets.** Datasets with outlying sequencing depth were determined based on median absolution deviation (MAD) using `isOutlier(type = "both")` from the R library `scuttle` [7] specifying the number of MADs to use.

***Table 2. Proportion of long-range contacts per** $c_p$ as percentage (%) of the total number of long-range contacts. See **Table 5** for the absolute counts. The 2- and 0.5-Mb sets include contacts with contact gap $\geq$ 2 and 0.5 Mb, respectively.*

| $c_p$ | **2 Mb** 112,861,349 | **0.5 Mb** 115,402,630 |
|---|---|---|
| | % | % |
| 1 | 17.45 | 17.07 |
| 2 | 17.05 | 16.68 |
| 3 | 14.54 | 14.22 |
| 4 | 11.66 | 11.42 |
| 5 | 9.18 | 8.99 |
| 6 | 7.21 | 7.07 |
| 7 | 5.67 | 5.57 |
| 8 | 4.44 | 4.37 |
| 9 | 3.44 | 3.40 |
| 10 | 2.62 | 2.61 |
| 11 | 1.97 | 1.98 |
| 12 | 1.45 | 1.50 |
| 13 | 1.06 | 1.15 |
| 14 | 0.77 | 0.89 |
| 15 | 0.55 | 0.71 |
| 16 | 0.38 | 0.58 |
| 17 | 0.26 | 0.50 |
| 18 | 0.16 | 0.43 |
| 19 | 0.09 | 0.36 |
| 20 | 0.04 | 0.29 |
| 21 | 0.01 | 0.19 |

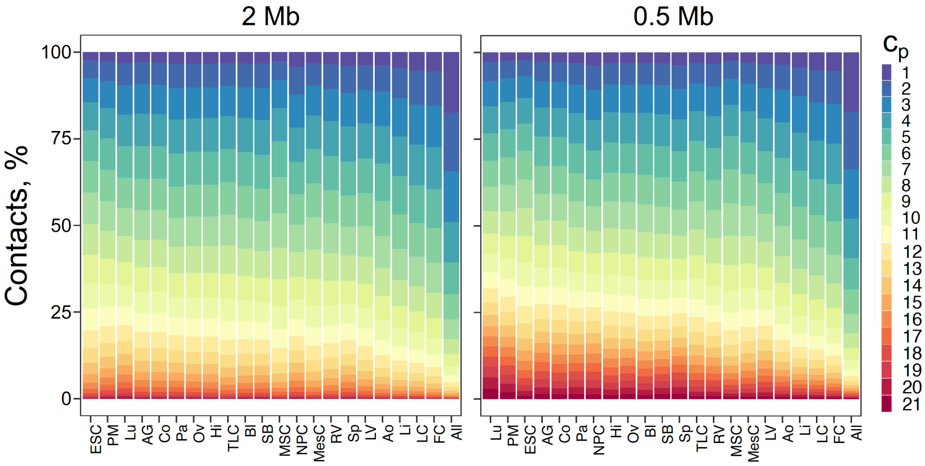

***Figure 3. Proportion of long-range contacts per** $c_p$ **in each of the 21 cell types.** See **Table 5** for the absolute counts. The colour indicates the $c_p$. Refer to **Table 1** for the abbreviations of the cell lines and tissues.*

#### 3.2 Contact persistence, $c_p$, in relation to Hi-C contact frequency, $c_f$

**Log transformation of HiCNorm $c_f$.** To avoid negative values, the HiCNorm $c_f$ values were subtracted by the minimum HiCNorm $c_f$, added by 1.001 and then $\log_{10}$-transformed. The calculation was done per cell type.

**Percentile rank of HiCNorm $c_f$.** The ranks of contacts with non-zero HiCNorm $c_f$ were determined per contact gap for each cell type, taking the maximum ranking in case of ties. The rankings were then converted to percentile rank by dividing by the number of contacts and multiplying by 100. With this method, a percentile rank of 20% means that 80% of contacts have higher $c_f$ value, and a percentile rank of 100% means that no contact exists with a higher $c_f$ value.

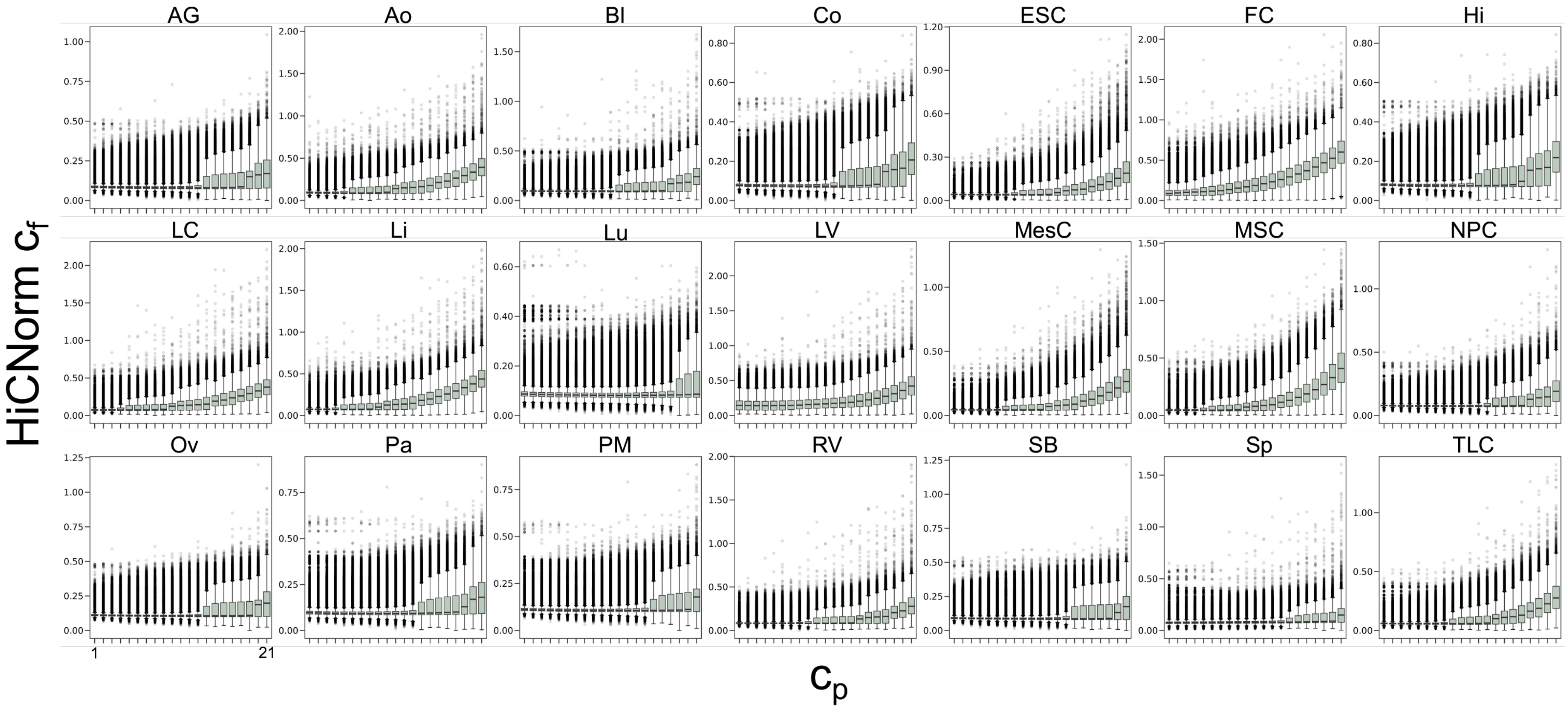

***Figure 4. Contact** $c_f$ **across** $c_p$ are shown (log$_{10}$-transformed* HiCNorm $c_f$*) per cell line and tissue.*

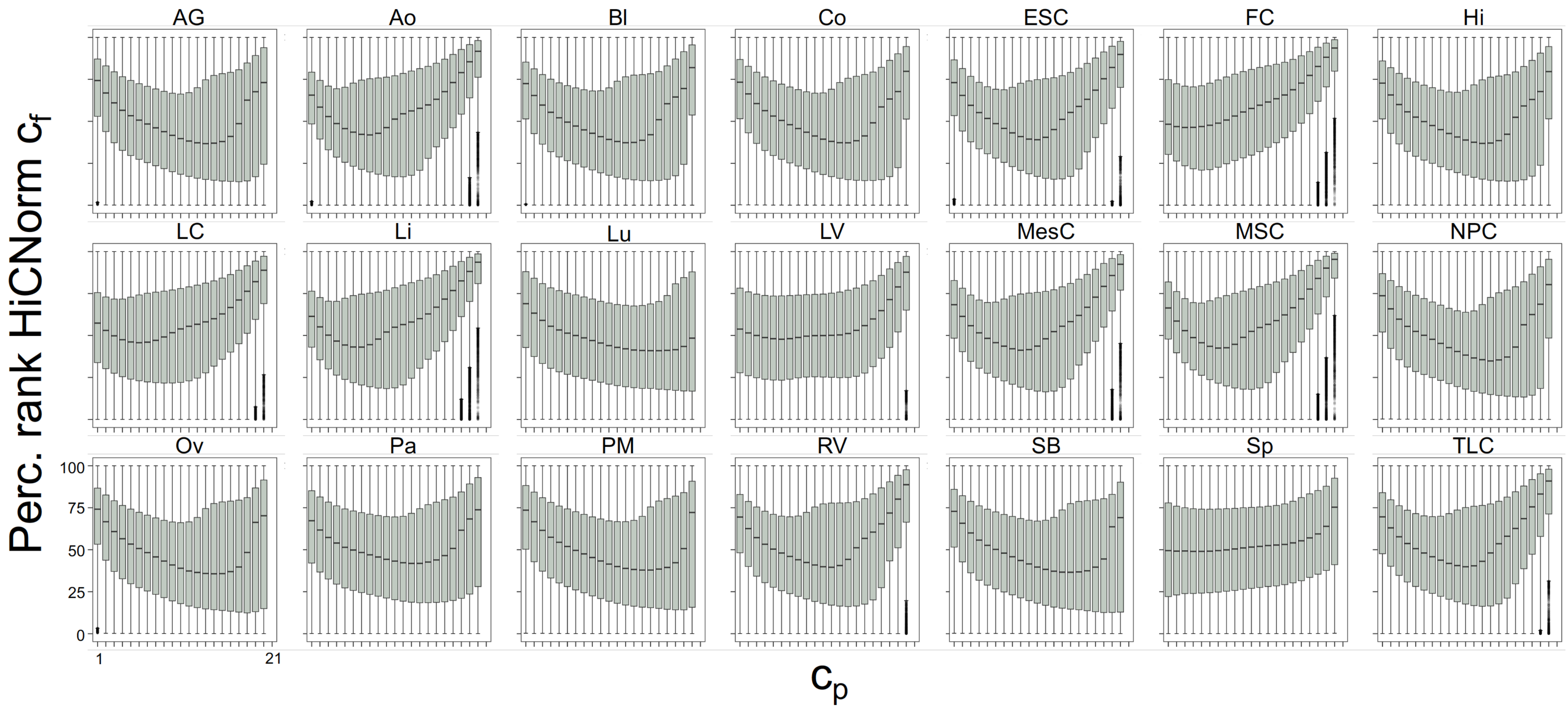

***Figure 5. Percentile ranks of contact $c_f$ per contact gap across $c_p$*** *are shown (HiCNorm $c_f$) per cell line and tissue. The percentile (perc.) rank of a contact $c_f$ was determined relative to the set of contacts of the same contact gap and per cell type.*

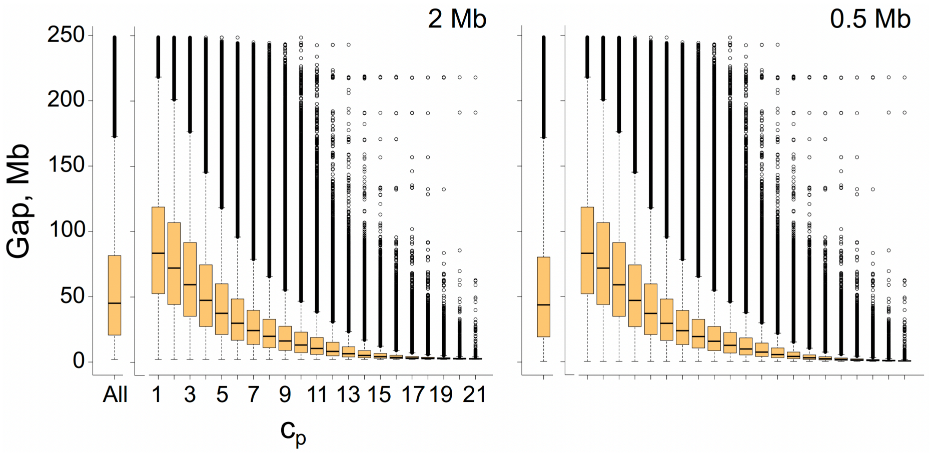

***Figure 6. Contact gaps across*** $c_p$. *Contact gap is the total number or length of intervening base pairs, excluding any base pair from the contacting 40-kb regions. "All" refers to all long-range contacts (*$c_p \geq 1$*).*

##### 3.3 Rationale for the contact persistence calculation

To explain why distance-based statistical enrichment tests was not applied to determine contacts in this study (omitted by design), we would like to revisit the interpretation of the physical actuality and the output of these statistical enrichment tests, which are also discussed in "The Hitchhiker's guide to Hi-C analysis: Practical guidelines" by Bryan Lajioe, Job Dekker and Noam Kaplan [8]. In the excerpt below, they talk about point interactions, which are statistically enriched interactions that have higher interaction frequency than expected (e.g. based on a background model of distance-decay, or comparing the frequency of the given contact with other neighbouring contacts):

"While the biological interpretation of point interactions seems to be straightforward, it is important to consider what such methods find. If we look for interactions that have a higher interaction frequency then what is expected given their distance, we are not evaluating their absolute interaction frequency. For example, consider two loci which are nearby in the genomic sequence, and are thus expected to interact very frequently. Such interactions may be functional and biologically important, but they may not have a much higher interaction frequency than expected by distance, and thus may not be found to be point interactions. Similarly, the expected interaction frequency for loci that are separated by large genomic distances is very low. As a result even a small increase in their interaction frequency can make their interaction statistically significant even though their absolute interaction frequency is still low, implying it occurs in only few cells. Thus, careful biological evaluation is always required in order for interpreting any statistical approach to identifying point interactions."

That said, contacts that are not flagged as "significant" by statistical enrichment tests are not guaranteed to be non-contacts (noise or an artefact of the method). They could still be valid contacts stably occurring in the genome, with their formation additionally facilitated by shorter distance, amongst myriads of other factors.

If we consider this phenomenon at a molecular level, the thermodynamic basis of the inverse distance dependence of genomic contacts boils down to the following fact: the more DNA sequence there is in between the two contacting sites, the more such a contact will contribute to an entropic loss in the genome by restricting a large chunk of DNA in a loop, hence such a contact can break easier, and may need to be rather enthalpically stable to survive the usual dynamics of the DNA polymer (the dynamical motions and pulls from the large loop in between). This exact phenomenon is true for also protein structures, where contact frequencies in large protein contact matrices also exhibit a certain degree of inverse distance rule. This rule, however, in no way makes a given well-characterised contact a “no contact” simply because it happens at a shorter distance range where there are more contacts to be generally expected. A backbone $C_\alpha$ - $C_\alpha$ contact in proteins, for instance, will never be discarded simply because it occurred between amino acids 5 residues apart rather than 12. Those are still valid contacts in the protein that contribute into the 3D structure and function of the protein. As another analogy, in some way, such a distance-based significance filtering will resemble the long-standing G-quadruplex structure (analogy of our contact) versus G content (analogy of our distance) debate in genome composition. At some point in past, attempt was made to focus on only the G-quadruplex structures that would not be the direct result of the high G content of a DNA sequence, as high G content can non-specifically result in G-quadruplex motifs. This type of significance filtering would focus our attention to G-quadruplexes that happen in the chunks of DNA with low G content, where it is less likely to have repeated tracks of Gs for the G-quadruplex formation, hence, if found, those should be more significant. With this, we may or may not zoom onto G-quadruplexes that were functionally selected or evolutionarily maintained, but this would certainly exclude all the most functionally relevant G-quadruplexes present in our genome that are actually located in G-rich gene promoter regions and regulate gene expression (including the famous G-quadruplex of the c-myc oncogene), even if the Nature’s way to form those is through maintaining the high G-content in the promoters.

We do think though that these statistical enrichment tests are very important particularly for certain applications. For example, some Hi-C-driven machine learning works performed the distance normalisation, but for the actual benefit of first eliminating the most obvious general trend (the “carrier wave”) for the contact prediction, to focus on the predictions of the perturbations, which was proven to improve the results by doing essentially a single pass of gradient boosting (a known phenomenon in machine learning), hence such a normalisation has a practical role there, rather than phenomenological. Some biological works perform such distance normalisation if wanting to look at functional enrichment of contacts. The basis here is that contacts that happen at the regions where there should be far fewer overall contacts should be extremely stable for that contact to survive, hence such a normalisation will focus on the most “stable” contacts, enriching in potentially more important functional contacts, which can then be experimentally validated for their functional role in certain biological processes. But, as pointed out previously, users of those statistical tests should be aware that such filtering could eliminate other relevant contacts that were just ignored because of lower contact frequency than the expected average value based on distance. In our work, we have not performed such a filtering by design: we want to work with all contacts with evidence of happening, i.e. a robust chimeric read representing that contact, as validated by all the underlying bioinformatics read-quality scores that comprise the pillars of the validity of all the sequencing technologies. That is why we also ensured that potential artefactual/ambiguously mapping reads were not accounted for at all. Working with longer-range contacts ($>$ 2 Mb, way greater than average TAD size [9]) also was a way to mitigate inclusion of significant noise from the method e.g. getting chimeric reads for contacts that are just extremely close to each other linearly. Furthermore, if a given contact was just deceptively passing all our checks being a mere artefact, we then had the voting of all the considered cell types to define the persistence of the contact, hence an artefact will not likely have a high contact persistence ($c_p$), hence influence our conclusions. Our method of identifying cell-type persistent contacts is independent of the actual value of contact frequency but in **Figure 5**, we did additionally inspect the contact frequency values. We converted the contact frequency ($c_f$) values to their percentile rank relative to other contacts with the same distance value and we did find that persistent contacts are not dominated by the lowest-ranked contacts. This means that for a certain distance value, persistent contacts tend to not be the contacts with the lowest contact frequency value.

##### 3.4 Associations with cross-species patterns of organisation

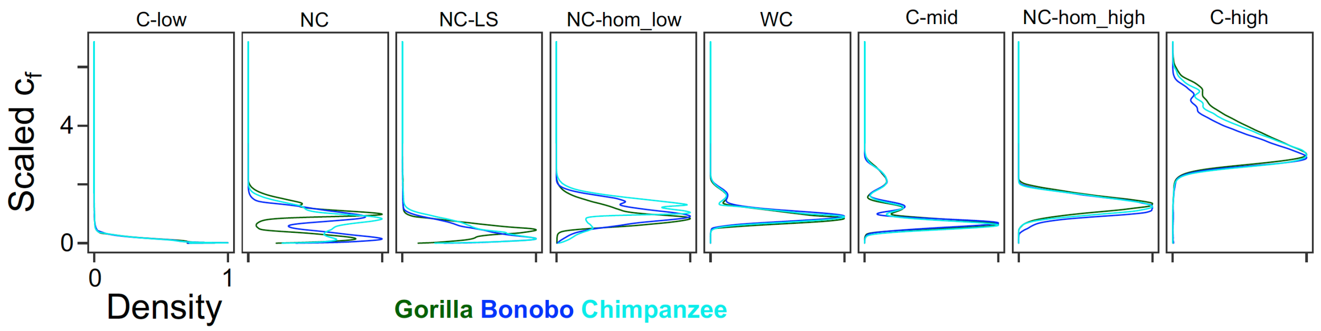

***Figure 7. Distribution of contact $c_f$ across* Phylo-HMRF *states determined based on pattern of $c_f$ across primates*** *reported in Yang et al. [10]. They determined lymphoblastoid cell 50-kb contacts within syntenic regions of 4 primates, human, chimpanzee, bonobo and gorilla (represented by different colours), by aligning the Hi-C reads from all species to the hg38 reference genome. They then categorised the contacts into different states based on the contact $c_f$ pattern across species. Measuring the similarity of distributions in terms of chi-squared distance, conserved states have distances smaller than expected by chance and compared with that of the non-conserved (NC) states. Conserved states as well as non-conserved ones are then differentiated into high, mid or low, based on the scaled value of $c_f$. NC-hom-high and NC-hom-low include contacts with distinct patterns in humans, i.e. they have higher and lower relative/scaled $c_f$ than their counterparts in the other species, respectively. The versions of these two states for the other primates were combined to comprise NC-LS, which are all non-conserved contacts with distinct $c_f$ patterns in each of the non-human primates. This combined 6* Phylo-HMRF *states into one as the distinction between those states is not needed for the analysis here. The human data were not available to be shown.*

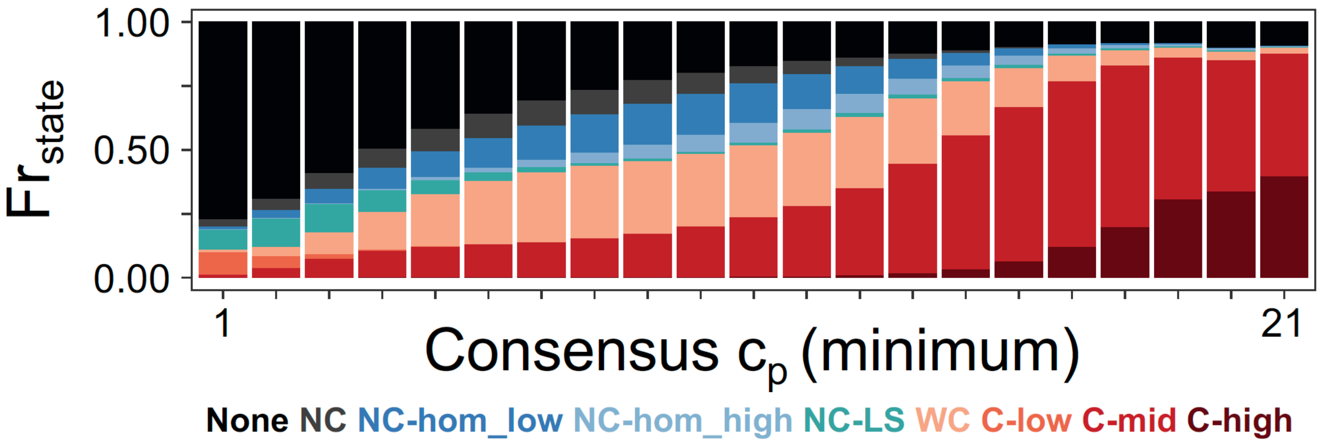

***Figure 8.* Phylo-HMRF *states across $c_p$.*** *Refer to caption of* ***Figure 7*** *for information on the* Phylo-HMRF *states. On the x-axis is the consensus $c_p$ for each hg38 contact. The value is equal to the ceiling of the minimum (for stringency) $c_p$ of hg19 contacts overlapping with the hg38 contact (see* ***Methods*** *for details). Results were found to be consistent when using the other summary statistics (maximum, minimum, median or mode) to calculate consensus $c_p$. Shown are fraction of contacts in a given state (represented by different colours) per ceiling consensus $c_p$ ($Fr_{state}$). In black are contacts with no state data.*

**Consensus $c_p$ for hg38 contacts.** We derived a consensus $c_p$ value for each of the hg38 contacts (50-kb resolution) using our hg19 $c_p$ contact data (40-kb resolution). Each hg38 contact was first converted to hg19 coordinates. Because an hg38, 50-kb region could be converted into multiple hg19 regions, a conversion is only considered valid when each of the two 50-kb regions of an hg38 contact maps to an hg19 region of length $\geq$ 30 kb. The consensus $c_p$ values for each hg38 contact are the mean, median, maximum, minimum and mode of the $c_p$ values of hg19, 40-kb contacts overlapping with the converted contacts. For multi-modal cases, the mean of the modes was calculated. To categorise the hg38 contacts across a $c_p$ axis, the ceiling values of floating point consensus $c_p$s were calculated, referred to in the text as ceiling consensus $c_p$.

### 4 Feature associations of persistent contacts

#### 4.1 Associations between Contact Persistence and Various Chromatin Elements

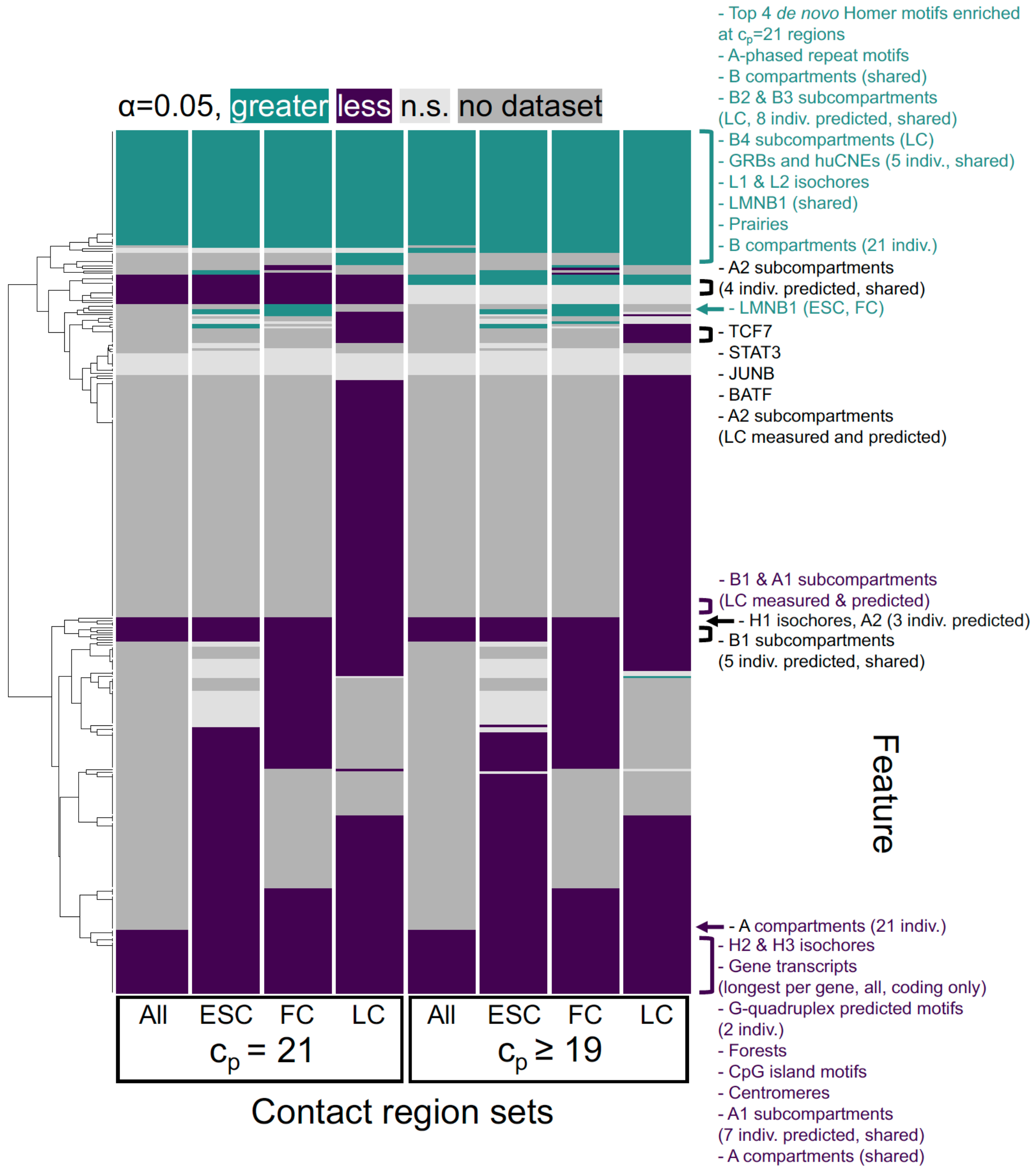

***Figure 9. Enriched and depleted features at persistent contact regions.*** *The set of all long range ($\geq$ 2 Mb) contact regions, i.e. with $c_p \geq 1$ was used as background. When the association involved features specific to a cell type, contact region sets were accordingly filtered to only include contacts present in that cell type. For cell-type-invariant features, the association was done with contact regions from all cell types, as well as with contact regions only from FC (IMR-90), ESC (H1-hESC) or LC (GM12878), to see whether the results are sensitive to the exact way of comparison. Association was considered significant when both the number of contact regions overlapping with feature regions and the total intersection in bp have a p-value $<$ 0.05. The first metric does not consider the portion of overlap, hence the use of the other metric. See attached* ***Supplementary file 2*** *for the data behind the heatmap and* ***Methods*** *for details on the permutation test. "Indiv." refers to individual data per cell type, individual data from different sources for G-quadruplex motifs or individual data conserved in each species for GRBs and huCNEs. "Shared" refers to regions shared by or common to all individual data for that feature. "Predicted" refers to subcompartment regions predicted by SNIPER [11].*

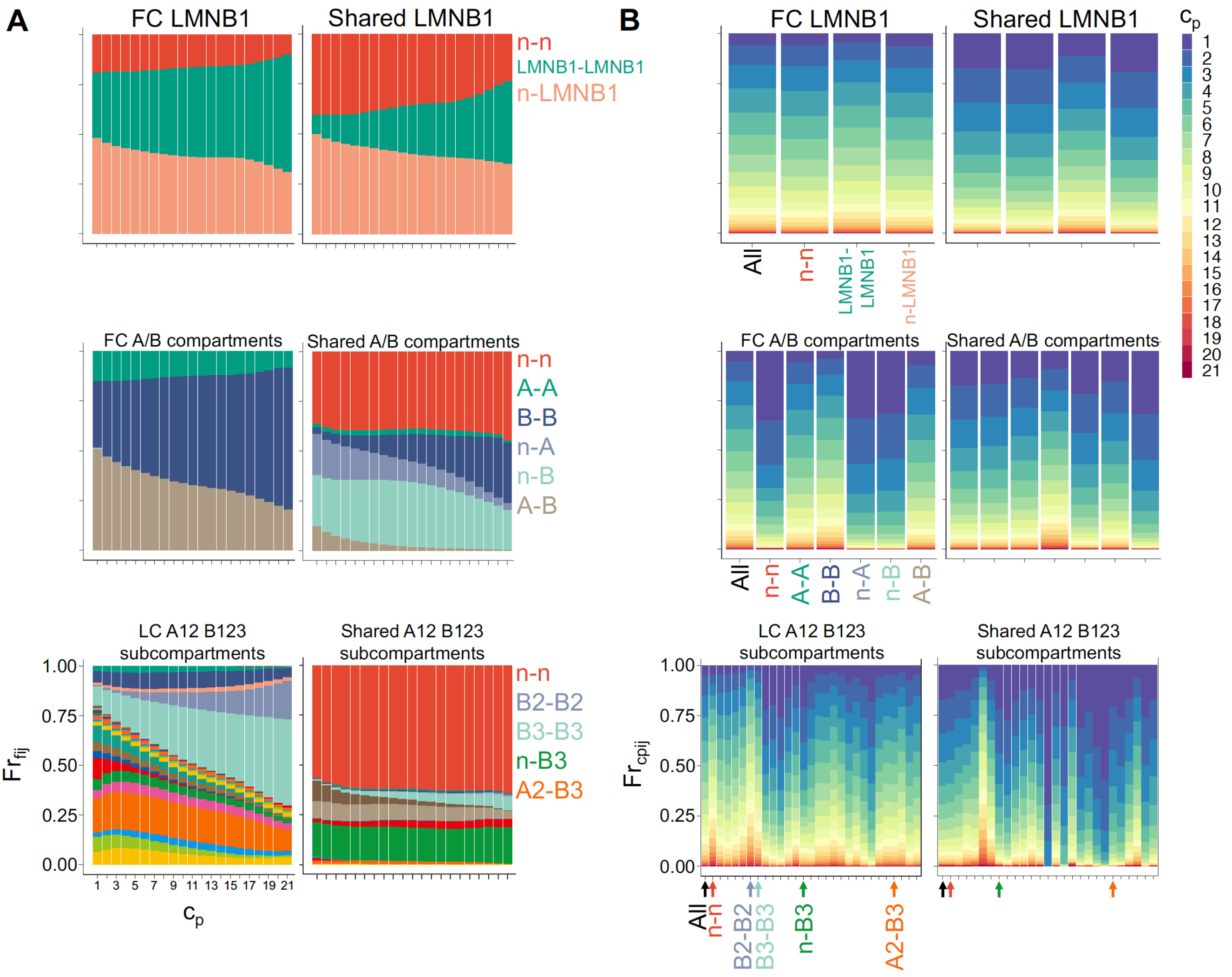

***Figure 10. Proportion of contacts within LADs and compartments across $c_p$.*** *(**A**) shows the fraction of contact types ($Fr_{fij}$) per $c_p$, while (**B**) shows the fraction of contacts of certain $c_p$ ($Fr_{cpij}$) per contact type. The classification of the contacts was determined by the overlaps of each of the two regions forming the contact, hence the region1-region2 type of notation brought in the plots. "n" and "All" mean none (no overlap between region and domain), and all long-range contacts, respectively. The domains and compartments are cell-type-specific features so, for stringency, only contacts present in the specified cell type were used. Features with the affix "shared" were generated by extracting regions shared by all cell lines or tissues with data. Shared LMNB1 regions are those common to FC (IMR-90) and ESC (H1-hESC) cell lines. Shared A/B compartments are regions common to all 21 cell types. Shared subcompartments are regions common to the LC (GM12878) data of Rao et al. [12] and the SNIPER predicted subcompartment data from 9 human cell lines (LC, HAP1, HeLa, HMEC, HSPC, HUVEC, FC, K562, and T cells) [11].*

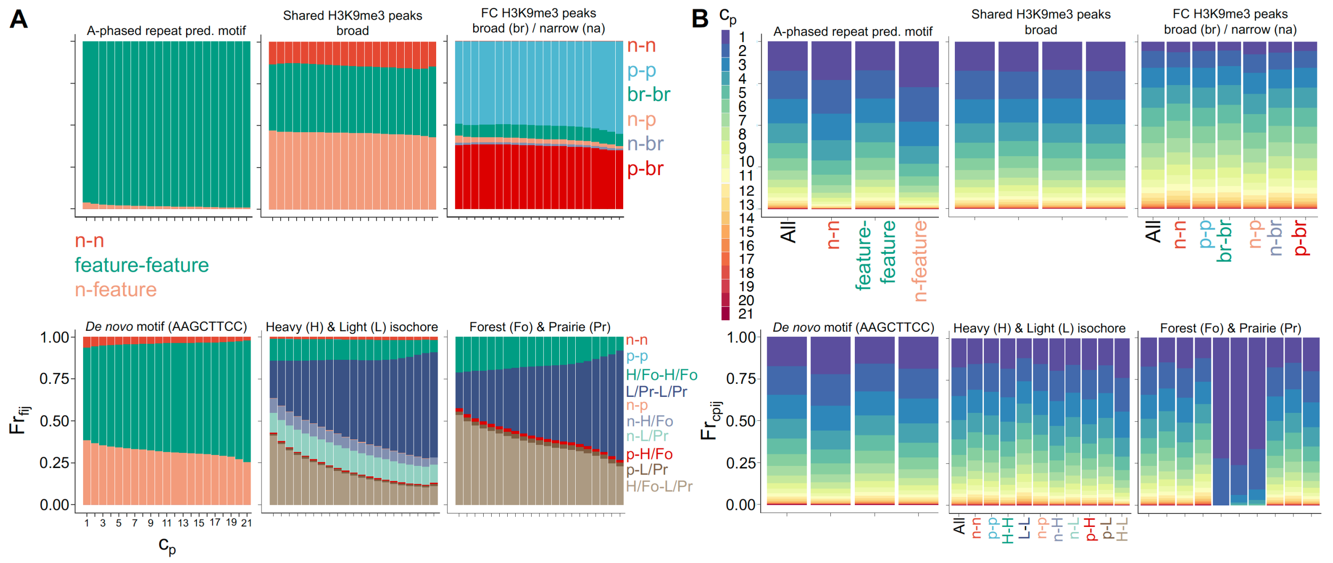

***Figure 11. Contact-wise association across $c_p$ of enriched features at persistent contact regions.*** *(**A**) shows the fraction of feature-defined contact types ($Fr_{fij}$) per $c_p$, while (**B**) shows the fraction of contacts of certain $c_p$ ($Fr_{cpij}$) per contact type. The feature-based classification of the contacts was determined by the overlaps of each of the two regions forming the contact, hence the region1-region2 type of notation brought in the plots. "n" and "p" mean none (no overlap between region and feature) and promiscuous (region overlaps with two types of feature). "All" refers to all long-range contacts. For cell-type-specific H3K9me3 data, only contacts present in the specified cell type were used. Shared H3K9me3 are broad peak regions common to FC (IMR-90), ESC (H1-hESC) and LC (GM12878) cell types.*

**BED files of features.** When necessary, the coordinates of the feature regions were converted to hg19, 1-based coordinates. The raw version of the feature regions had consecutive and overlapping ranges and for certain purposes, `reduce()` from the R library `GenomicRanges` was applied to make a version in which overlapping and consecutive ranges are joined. To derive the isochore data, regions with GC content data from Costantini et al. [13] were classified into the following isochore families based on a modified version of the % GC content ranges of families reported in Jabbari and Bernardi [14]: L1 ($<$ 36.4%), L2 (37.6-39.6%), H1 (42-45%), H2 (46.9-52%), H3 ($>$ 54%). The original ranges were overlapping so to avoid classifying some regions into families with high uncertainty, we modified them to become open, disjoint ranges. **Table 7** has the complete list of features used.

**Region-wise association or enrichment methods.** Chromatin and genomic features, which are characteristics of a region (not by a contact), were associated with the unique contact regions per $c_p$ *via* two ways. **1)** The significance of feature enrichment in a set of unique contact regions was quantified by doing a permutation test (10,000 iterations) using `permTest()` from the R library `regioneR` (the rest of the functions mentioned are from the same library). The contact regions were the ones being permuted and the random samples were drawn without replacement from the background that was the set of all unique long-range contact regions ($c_p \geq 1$) using `resampleRegions()`. The association was quantified by calculating: **a)** number of contact regions overlapping with feature ranges, where an overlap of one contact region with multiple regions of a feature was counted only once (`numOverlaps()`) and **b)** total intersection in bp (`commonRegions()`). The alternative hypothesis was determined in `permTest()` by comparing the mean of the random distribution and the evaluated value. Note that this function also sets a minimum p-value to be returned equal to inverse the number of permutations or 0.0001 in this case. The same permutation test procedure was used for calculating the significance of enrichment of long genes at high-$c_p$ unique contact regions except that the sample statistic was the mean length of genes overlapping. **2)** The number of unique contact regions per $c_p$ that overlap with a feature was calculated using custom R scripts.

**Contact-wise association or enrichment methods.** The contact-wise association was done *via* two ways. **1)** Per $c_p$, we determined the fraction of feature-defined contact types based on whether the two regions of a contact overlap or do not overlap with a feature. When using multiple features to categorise a contact, promiscuous cases may arise i.e. a region overlapping with multiple features. **2)** Per feature-defined contact type, we calculated the fraction of contacts of given $c_p$.

#### 4.2 Differential k-mer composition of contact regions

***Table 3. Proportion of significantly changed 7-mers per $c_p$.*** *Number of significantly changed 7-mers per $c_p$ and the percentage (%) of enriched and depleted 7-mers. The count of all possible 7-mers is $4^7$, i.e. 16,384.*

| $c_p$ | 2 Mb | | | 0.5 Mb | | |
|---|---|---|---|---|---|---|
| | Total sig. k-mer | % Depleted | % Enriched | Total sig. k-mer | % Depleted | % Enriched |
| 1 | 0 | 0 | 0 | 0 | 0 | 0 |
| 2 | 0 | 0 | 0 | 0 | 0 | 0 |
| 3 | 0 | 0 | 0 | 0 | 0 | 0 |
| 4 | 0 | 0 | 0 | 0 | 0 | 0 |
| 5 | 0 | 0 | 0 | 0 | 0 | 0 |
| 6 | 0 | 0 | 0 | 0 | 0 | 0 |
| 7 | 0 | 0 | 0 | 0 | 0 | 0 |
| 8 | 8 | 75 | 25 | 0 | 0 | 0 |
| 9 | 1550 | 46.19 | 53.81 | 0 | 0 | 0 |
| 10 | 6334 | 42.78 | 57.22 | 0 | 0 | 0 |
| 11 | 9272 | 43.83 | 56.17 | 0 | 0 | 0 |
| 12 | 11262 | 46.01 | 53.99 | 0 | 0 | 0 |
| 13 | 12520 | 48.51 | 51.49 | 46 | 86.96 | 13.04 |
| 14 | 13642 | 50.24 | 49.76 | 1250 | 49.12 | 50.88 |
| 15 | 14402 | 51.53 | 48.47 | 5264 | 41.00 | 59.00 |
| 16 | 14844 | 52.26 | 47.74 | 7996 | 43.82 | 56.18 |
| 17 | 15180 | 52.67 | 47.33 | 10192 | 44.15 | 55.85 |
| 18 | 15462 | 53.21 | 46.79 | 11888 | 47.07 | 52.93 |
| 19 | 15618 | 53.45 | 46.55 | 13100 | 49.13 | 50.87 |
| 20 | 15682 | 53.79 | 46.21 | 14072 | 50.51 | 49.49 |
| 21 | 15562 | 53.90 | 46.10 | 14872 | 51.33 | 48.67 |

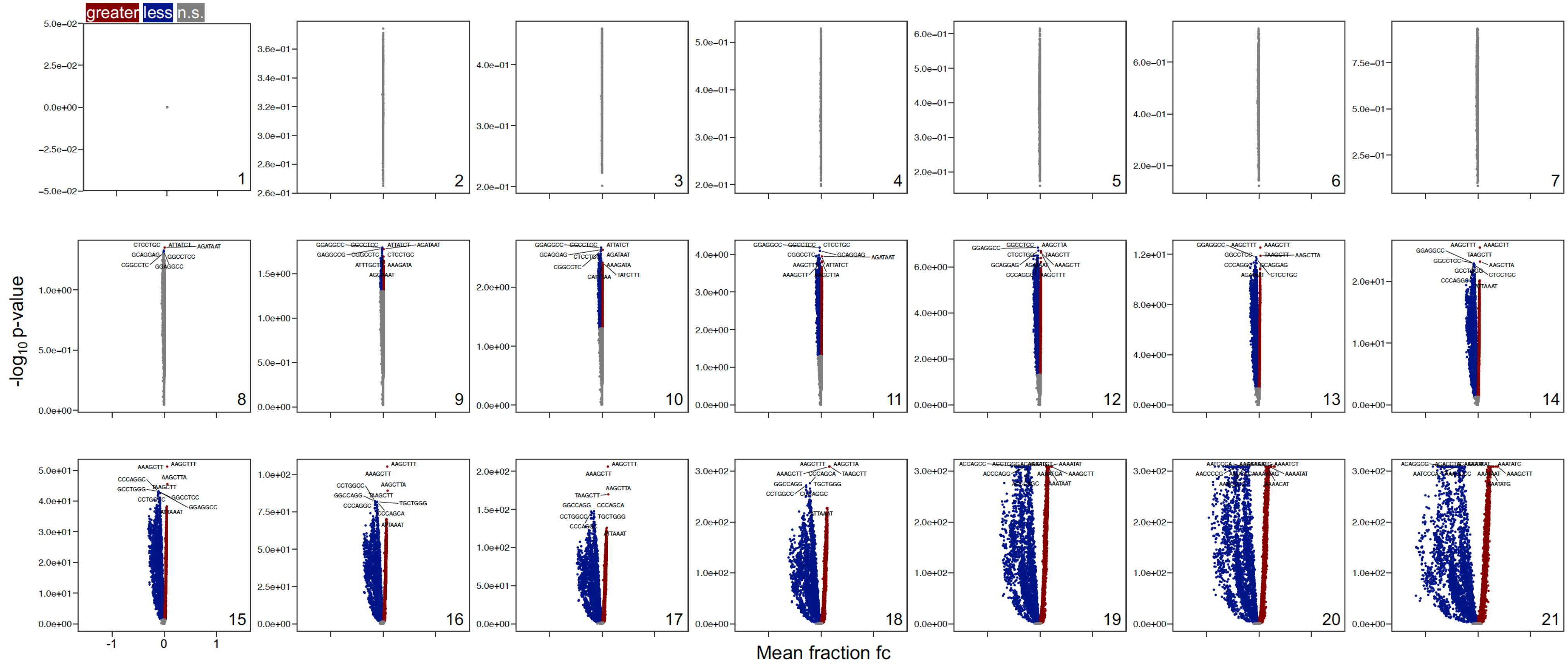

**Figure 12. K-mer enrichment analysis of contact regions across $c_p$.** *Shown are significantly enriched (in red) and depleted (in blue) 7-mers at unique contact regions per $c_p$ (see **Methods** for details). K-mers not significantly changed are in grey. The set of unique contact ($c_p \geq 1$) regions was used as background. The $c_p$ is shown at the bottom right corner.*

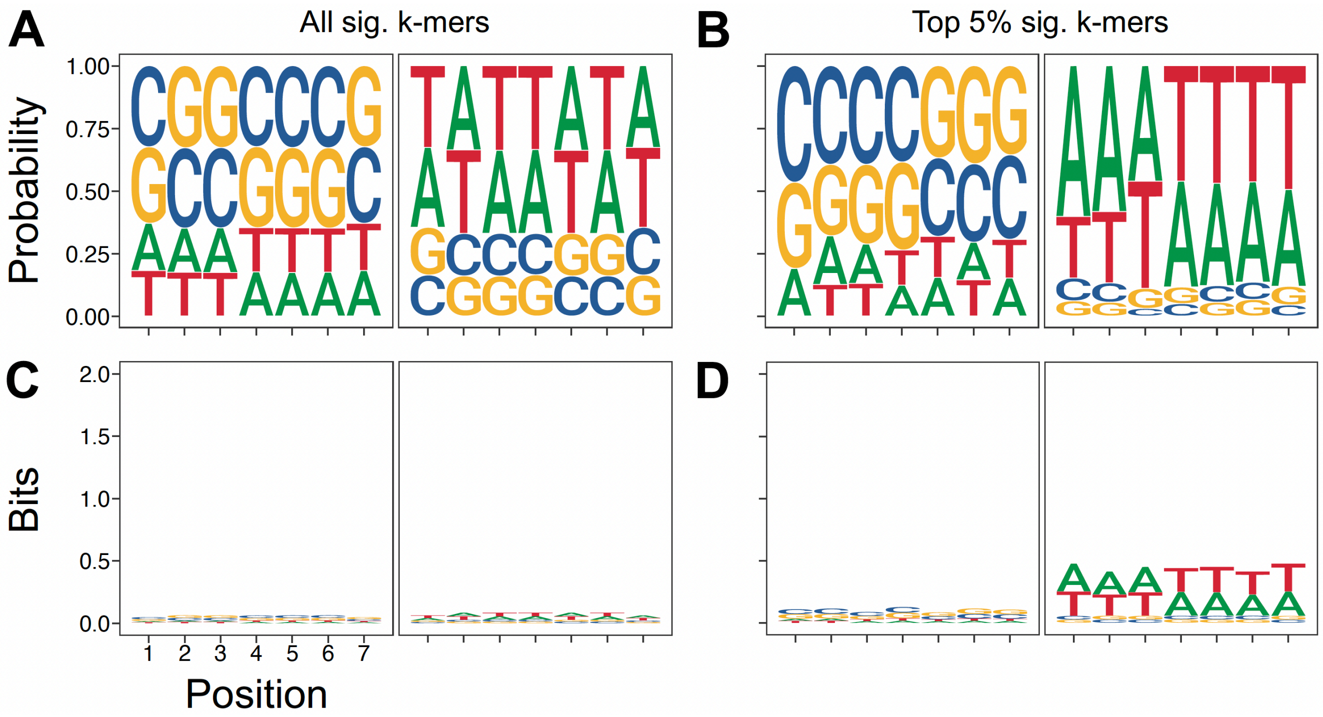

***Figure 13. Sequence logos of significantly enriched and depleted 7-mers at prime contact regions.*** *Plots are in terms of (**A-B**) probability and (**C-D**) information content (in bits) values. Versions were made for (**A** and **C**) "All" and (**B** and **D**) only top 5% of significantly depleted and enriched 7-mers ($\alpha = 0.05$).*

Note that contacts can only have one $c_p$ value, but the regions that form these contacts can be shared across $c_p$s. In fact, as mentioned in the associated studies, unique $c_p = 1$ set of regions are equivalent to the background, i.e. to all regions forming our working set of long-range contacts; hence, its volcano plot shows no significantly changed 7-mers. With increasing $c_p$, significant differences in 7-mer content start to emerge. At prime contact regions, there are 15,562 7-mers that are significantly changed, split into 53.9% depleted and 46.1% enriched ones ($\alpha = 0.05$). The most depleted k-mers are CGCGCGA, TCGCGCG and CGGCCCG with fold changes of ≈ -1.18. The most enriched ones are AAGCTTA, TAAGCTT and AAAGCTT with fold changes of ≈ 0.38.

**K-mer enrichment analysis at contact regions.** The 7-mer contents were calculated as fractional values at each region averaged per $c_p$ category. This was done by first dividing the absolute count of a given 7-mer from both strands (not scaled as in the $c_{||}^{k-mer}$ calculation) by the sum of counts for all possible 7-mers in that region. This sum of counts is constant for a certain k-mer length $k$ ($k = 7$ in this case) and region length $L$ and is given by $(L - k + 1) \times 2$ (multiplied by 2 to account for both strands). Given a 7-bp k-mer and 40-kb region, the sum would be equal to 79,988 bp. This sum was constant in our case because regions with at least one missing base pair were excluded from the analysis. The mean fraction of a given 7-mer per $c_p$ was then derived by taking the mean of its fractional values from all regions belonging to that $c_p$. The mean fraction fold change (fc) of a 7-mer, brought on the $x$ axes in **Figure 12**, is the fc of the mentioned average content relative to that in the control. The p-values, brought in the $y$ axes in **Figure 12**, are from the MWW tests done in a directional manner. If the mean fraction of the sample was greater or less than that of the control, the alternative hypothesis was accordingly taken as greater or less. If mean values were equal, the alternative hypothesis was checked in a "two-sided" manner. The `wilcox.text()` function results in a p-value of 0 in cases where the value was extremely small, beyond the allowed double precision storage limit. For plotting such cases in a log scale, those were converted to the maximum significance -$\log_{10}$-transformed p-value observed for the whole dataset, considering all $c_p$ values. Ultimately, this approach gave us a mean fraction and a p-value for each of the 16,384 7-mers per $c_p$.

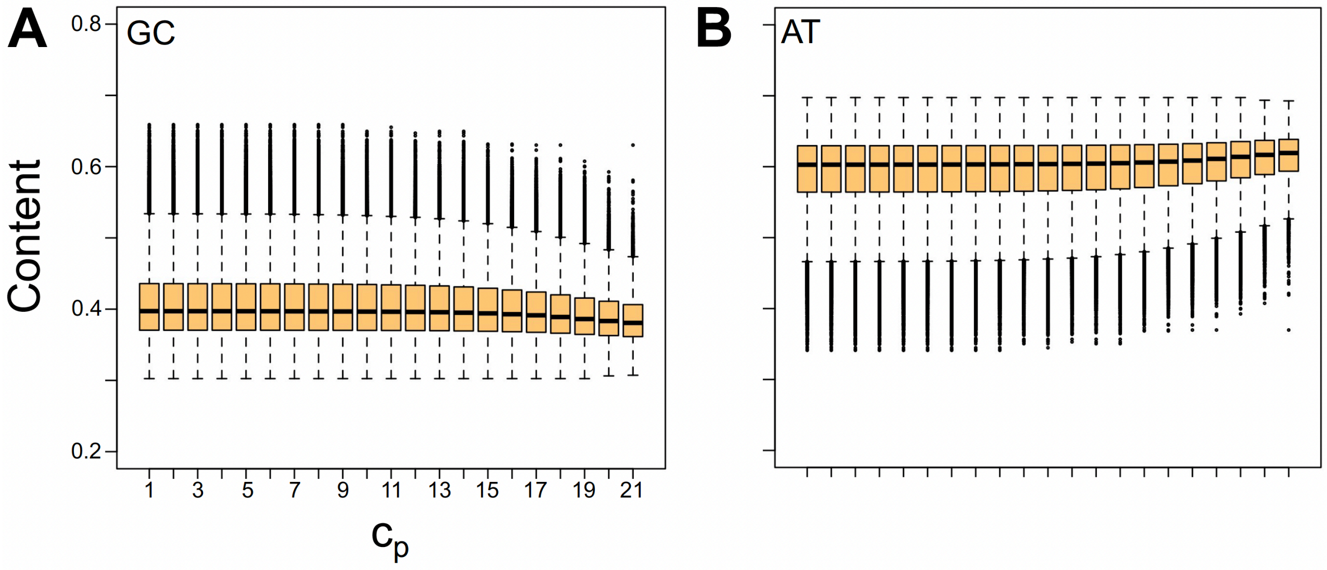

***Figure 14. GC and AT contents of contact regions across*** $c_p$ *as fractions of the 40-kb sequence.*

#### 4.3 Associations between Contact Persistence and Gene-Related Features

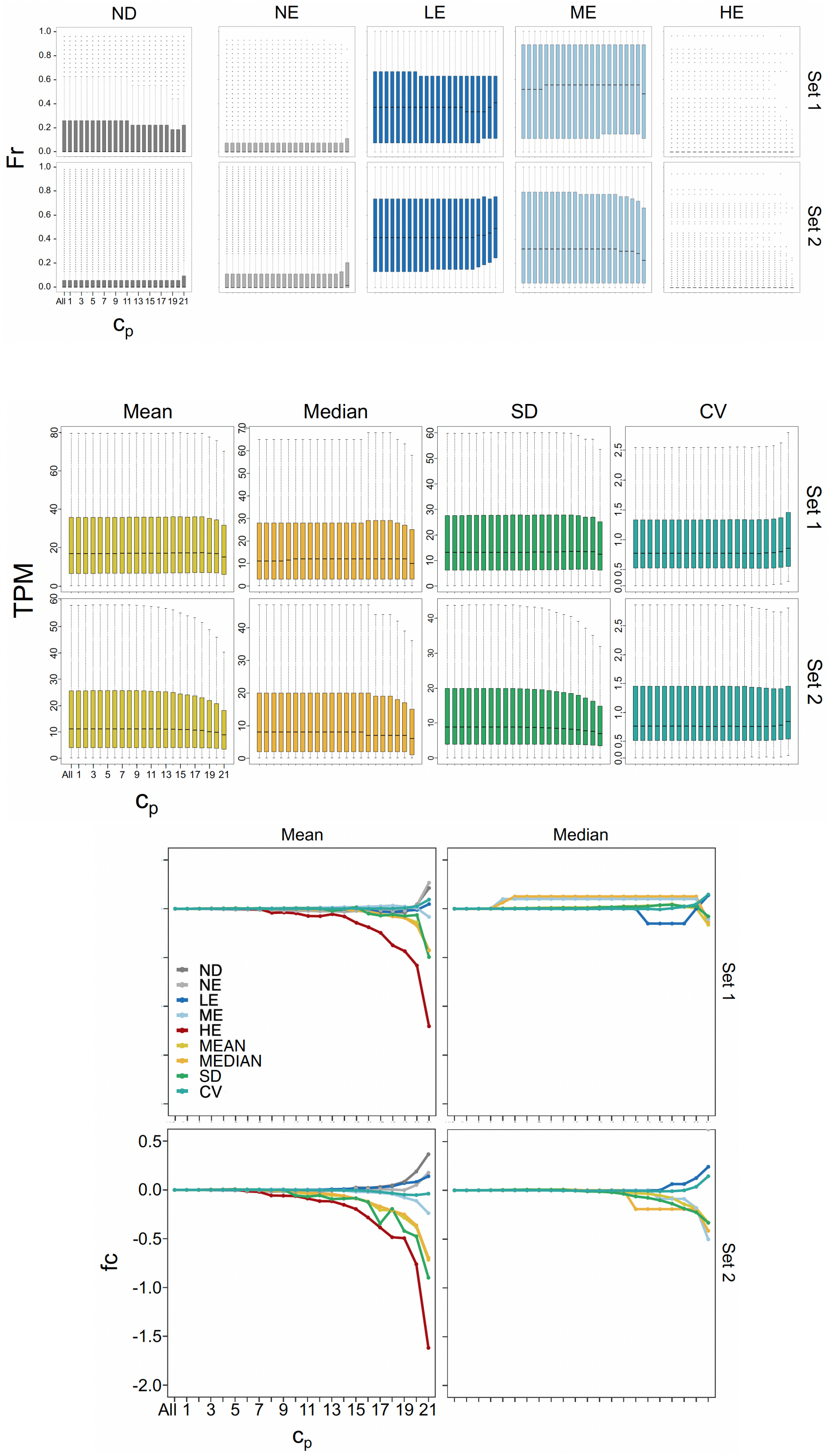

***Figure 15. Cross-tissue average gene expression and variation across $c_p$.*** *Mean, median, standard deviation (SD) and sd/mean coefficient of variation (CV) of expression values are shown, in TPM, of genes per $c_p$. Only genes with data in at least 70% of the tissues were considered. Fold changes (fc) of the mean and median values of distributions relative to value at $c_p = 1$ are also shown. No line for HE in the Median plots because the median across $c_p$s is 0. The $c_p \leq 3$ and $c_p \geq 19$ are significantly different (p-values $<$ 0.002) except for the CV (p-value of 0.105).*

**Processing of gene expression datasets.** Two sets of human average baseline expression data (in TPM), generated *via* RNA-seq, were obtained from EMBL-EBI Expression Atlas [15]. Set 1 is the expression data of coding RNAs from 27 normal tissues from 95 adult individuals [16]. Set 2 is the expression data from 53 normal tissues from the Genotype-Tissue Expression (GTEx) project [17]. Sets 1 and 2, filtered to contain only genes with expression data in at least 1 tissue, have 80.31% (20,006) and 81.43% (20,284) of 24,910 unique genes in the UCSC hg19 gene annotation table, respectively. For genes with multiple entries in the data (only 17 and 7 genes for Sets 1 and 2, respectively), values were added per tissue. The definition of the expression levels was taken from EMBL-EBI i.e. not expressed (NE): TPM/FPKM $< 0.5$, low-expressed (LE): TPM/FPKM = [0.5,10], medium-expressed (ME): TPM/FPKM = (10,1000], and high-expressed (HE): TPM/FPKM $> 1000$ [15].

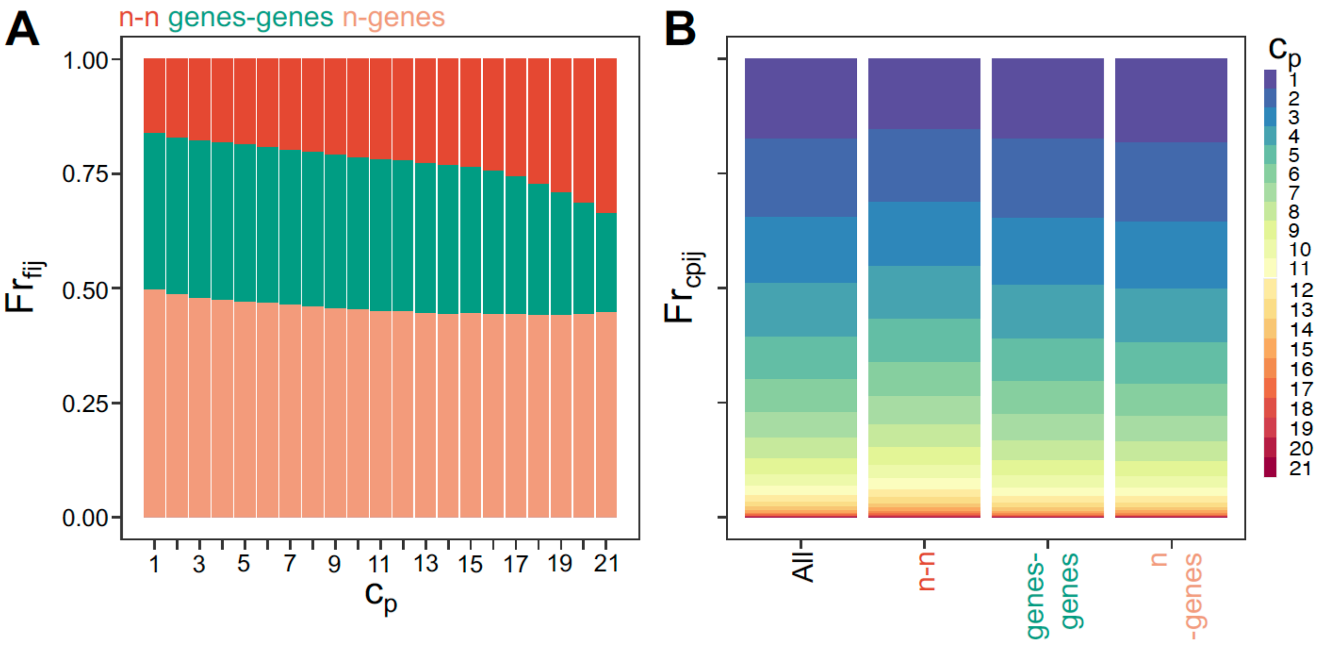

***Figure 16. Contact-wise enrichment of genes across*** $c_p$***.*** *(**A**) shows the fraction of contact types ($Fr_{fij}$) per $c_p$ defined by overlap of contact regions with genes. (**B**) shows the fraction of contacts of certain $c_p$ ($Fr_{cpij}$) per contact type. The classification of the contacts was determined by the overlaps of each of the two regions forming the contact with genes, hence the region1-region2 type of notation brought in the plots. "n" and "All" mean none (no overlap between region and genes) and all long-range contacts, respectively.*

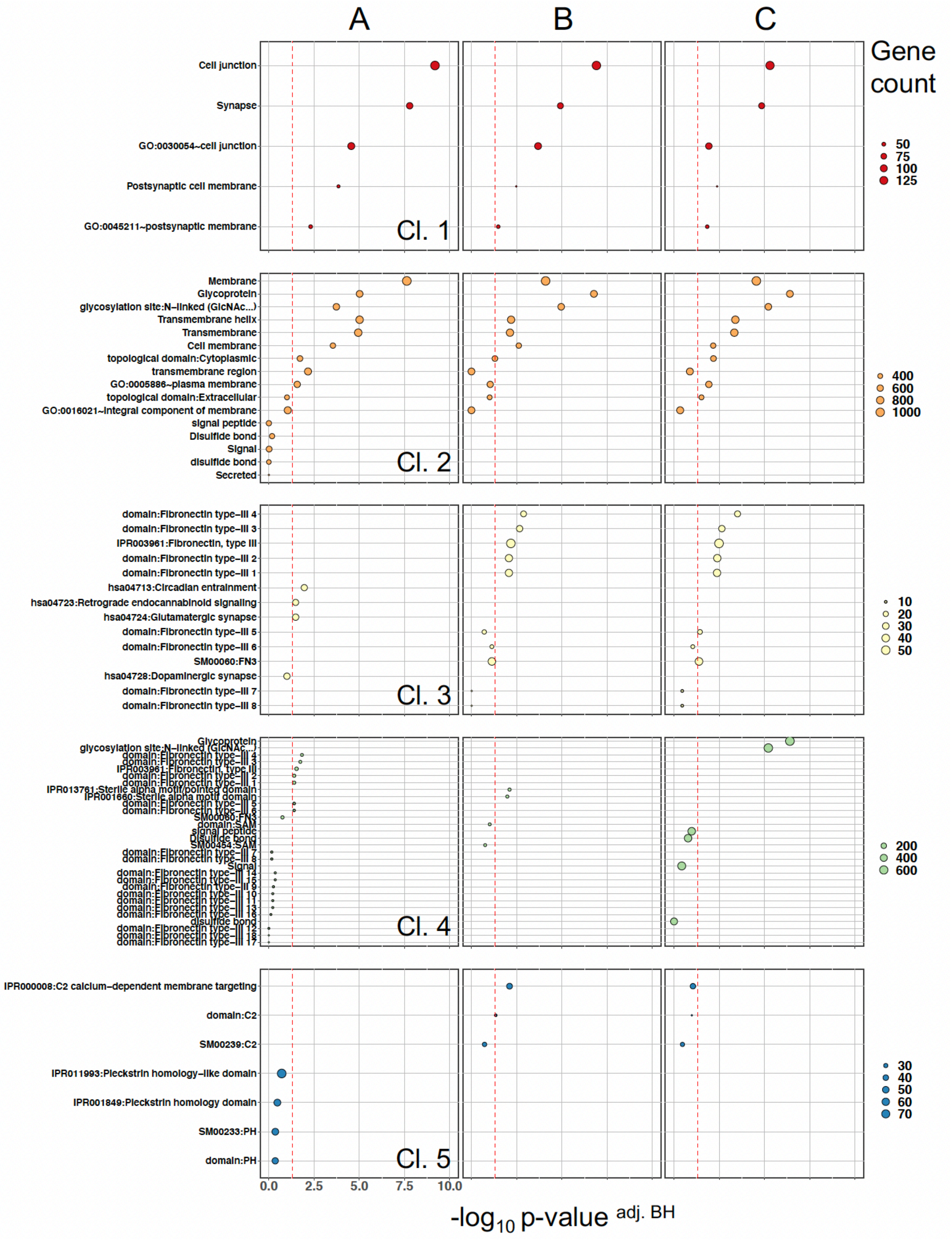

***Figure 17. Top 5* DAVID *clusters (Cl.) of functional terms related to genes at prime contact* ($c_p = 21$) *regions.*** *P-values are Benjamini-Hochberg (BH) adjusted (adj.). Red dashed lines mark -*$\log_{10}$ *0.05* $\approx$ *1.3010, and only terms with significance level greater than this value in at least 1 sample (from A, B and C) are displayed. Gene count is the number of prime contact genes associated with each term. The background used was the* DAVID *built-in set of Homo sapiens genes. See* **Methods** *for details.*

**Functional term enrichment analysis with DAVID.** Because DAVID can only be used to process up to 3000 inputs at a single instance, 3 sets (denoted as A, B and C) containing 2999 genes were randomly sampled without replacement from 4209 unique genes co-localising with prime contact ($c_p = 21$) regions. Only 2999 was sampled because in some instances, DAVID maps more than 1 DAVID gene identifier to a gene name. The following settings for DAVID functional annotation clustering were applied (built-in medium stringency): Kappa Similarity (Similarity Term Overlap=3, Similarity Threshold=0.5), Classification (Initial Group Membership=3, Final Group Membership=3, Multiple Linkage Threshold=0.5) and Enrichment Threshold (EASE=1.0). The enrichment or EASE score of a cluster is the -$\log_{10}$-transformed geometric mean of the enrichment p-values (a modified Fisher exact p-value) of member terms [18, 19]. The EASE scores of clusters in **Figure 17** range from 2.186 to 8.177 with Cluster (Cl.) 1 having the highest score.

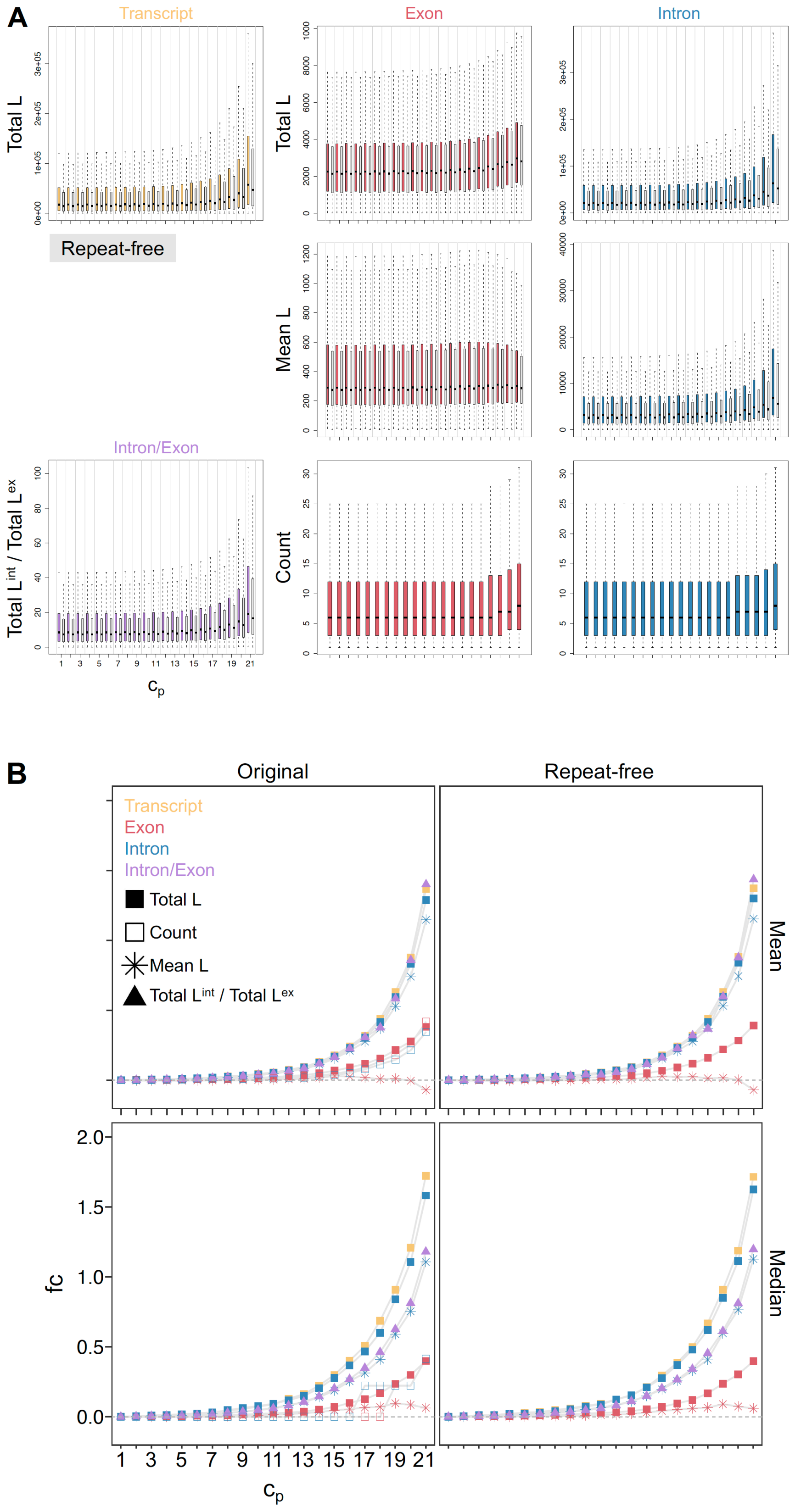

***Figure 18. Lengths and counts of genes and their components across*** $c_p$***.*** *Total transcript (tr), exon (ex) and intron (int) lengths (L), mean exon and intron lengths, exon and intron counts, and total intron and total exon length ratio of genes at contact regions per* $c_p$ *are shown. The distributions (****A****) as boxplots and (****B****) their central values (mean and median). Repeat-free lengths were obtained by subtracting repeat elements on genes (corresponding grey boxplots). The repeat-free mean lengths were calculated using the original exon and intron counts. Repeats include all sites in the UCSC hg19 RepeatMasker annotation table. Permutation test showed that the mean length of genes overlapping with* $c_p = 21$ *and* $c_p \geq 19$ *contact regions is significantly higher than* $c_p \geq 1$ *genes (p-value < 0.0001).*

#### 4.4 Replication timing and $c_p$

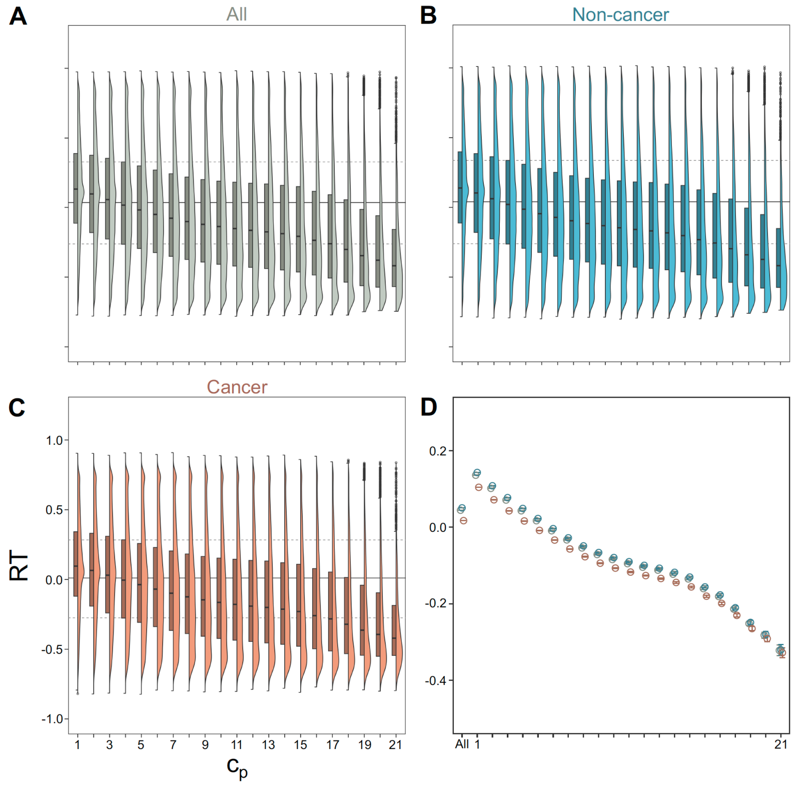

***Figure 19. Replication timing across $c_p$.*** *Values are the consensus replication timings (RT), given by the $log_2$ ratio of early and late signals, for each contact. A region was represented by the mean or median of the RT measurements overlapping with it. The consensus RT for a contact was then calculated as the mean of the two means or two medians from the two contacting regions. Results were comparable when taking the mean of the medians instead. Distributions of RT values across $c_p$s from (**A**) all ($N = 61$), (**B**) non-cancer only ($N = 50$) and (**C**) cancer-related only ($N = 11$) cell lines (colours indicate sample group). Solid and dashed horizontal lines mark the median and the 25$^{th}$ and 75$^{th}$ percentiles of the RT distribution considering all contacts. All pairwise comparisons of the distributions from each $c_p$ and from all long-range contacts are significantly different (BH $P$ value $<$ 0.002). (**D**) Shown are the means of the distributions in (**A**-**C**) with error bars marking the 95% confidence intervals. The $c_p$ =“All” refers to all contacts.*

**Replication timing data.** The hg19 replication timing (RT) data were downloaded from Replication-Domain (`http://www.replicationdomain.org`) [20] and processed using custom R scripts written by James Ashford for the hg38 versions of the data. There were 192 samples obtained from the repository, counting technical and biological replicates. Only 155 were used for subsequent analyses after removing those not from humans and those with data from chr. 21 only. RT measurements were binned at 40 kb to match the contact data. For each 40-kb bin or region, measurements from technical replicates were averaged reducing the number of unique samples to 117. Regions with less than 3 data points coming from a sample even after combining technical replicates were excluded from the subsequent biological replicate averaging, which further reduced the number of unique samples to 61 (referred to as 61 unique cell lines). This yielded a dataset, wherein a 40-kb region has one average RT value from each of the 61 unique cell lines (50 non-cancer- and 11 cancer-related cell lines). For the association with $c_p$, the mean and median of the average values from each group of cell lines were calculated for each 40-kb region. Only regions with data from $\geq$ 59 cell lines were considered, which represented 95% of all possible 40-kb regions from all human autosomes and chr. X.

##### 4.5 Somatic mutations and $c_p$

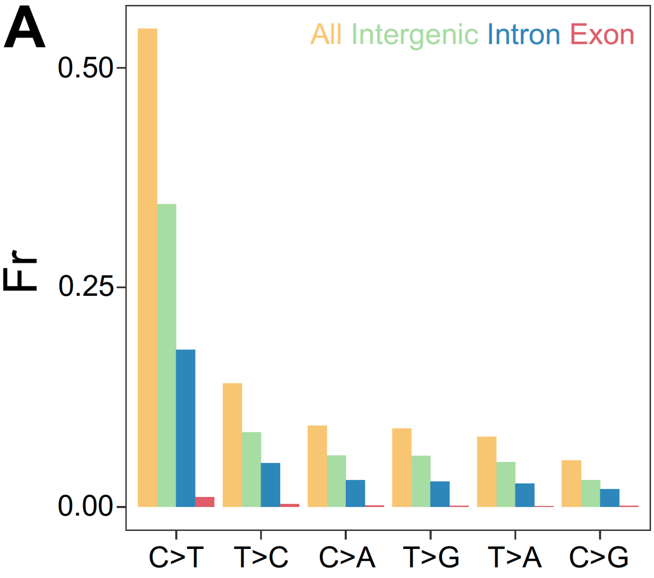

***Figure 20. Proportion of SNVs per mutation type and location.*** *The fractional value (Fr) is relative to the total number of SNVs in the final dataset, 38,428,969. Symmetric mutations were combined adopting the COSMIC convention to denote SNV types, C $>$ T (G $>$ A), C $>$ G (G $>$ C), C $>$ A (G $>$ T), T $>$ G (A $>$ C), T $>$ C (A $>$ G), T $>$ A (A $>$ T). Colours denote the location. Types are arranged in the plot based on decreasing fractions when combining locations. Intergenic was defined as 2000 bp upstream and downstream of transcript boundaries. See* ***Methods*** *for details on the SNV dataset.*

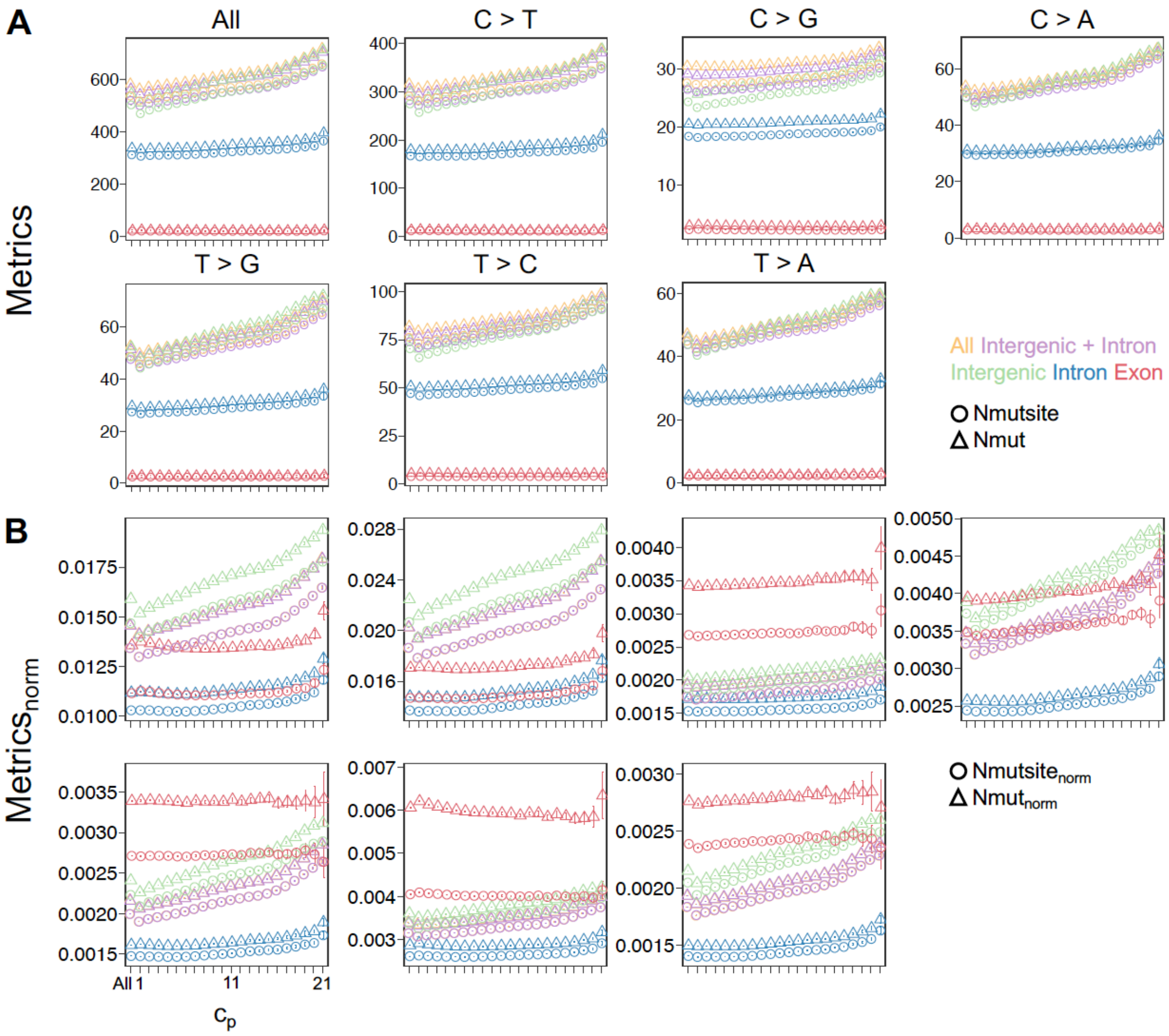

***Figure 21. Mean values of mutation metrics across $c_p$ per mutation type and location.*** *"All" refers to all SNV types. Colours denote location. Shown are the means of the distributions across $c_p$ with error bars marking the 95% confidence intervals for (**A**) $Nmutsite$ and $Nmut$, and (**B**) $Nmutsite_{norm}$ and $Nmut_{norm}$. Point shapes differentiate $Nmutsite$/$Nmutsite_{norm}$ (circle) from $Nmut$/$Nmut_{norm}$ (triangle).*

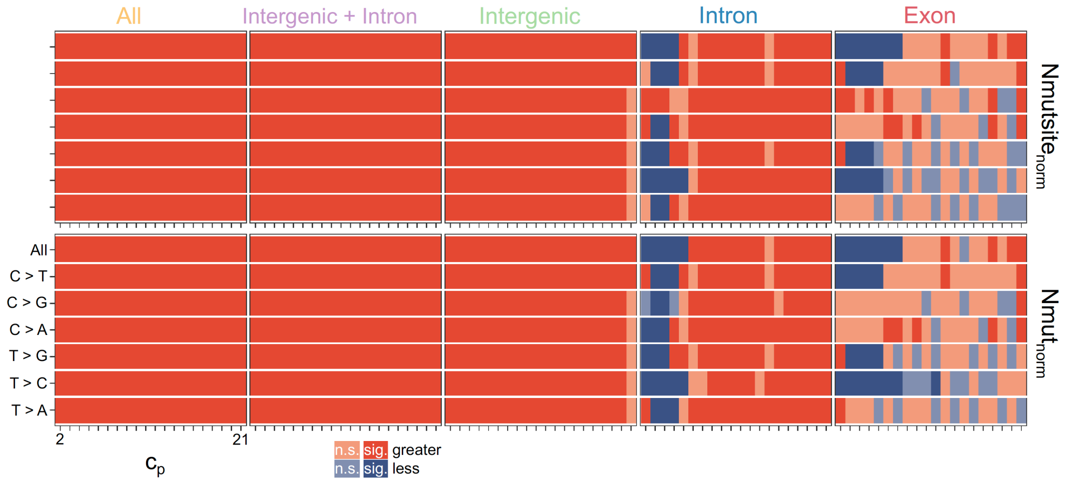

***Figure 22. Significance of trends of mutation metrics across $c_p$.*** *Dark orange (or violet) means that the mean of the metric distribution at a $c_p$ (e.g. $c_p = 2$) was greater (or less) than the corresponding mean at the previous $c_p$ (e.g. $c_p = 1$) and that the difference between the distributions was significant (sig.) (BH $P$ value $<$ 0.05, MWW test). If not significant (n.s.), corresponding lighter shades of the colours were used.*

**Somatic cancer SNV data.** Two datasets were used both coming from the International Cancer Genome Consortium (ICGC). The main dataset was downloaded from the ICGC Data Portal (https://dcc.icgc.org/, Release 28, 2019 March 27) [21] with the following Donor filters: Donor Analysis Type=WGS (Whole-Genome Sequencing), Available Data Type=SSM (Simple Somatic Mutation); and Mutation filters: Type=Substitution, Analysis Type=WGS (Whole-Genome Sequencing), Verification Status=Not tested (majority of the mutations), Tested and verified, Tested and inconclusive. The other dataset includes mutation calls from samples or specimens that are part of the Pan-Cancer Analysis of Whole Genomes (PCAWG) study [22], and was retrieved from UCSC Xena (https://xena.ucsc.edu/) [23]. Datasets only contain samples from projects not from the United States (US) (since US samples only have coding mutations) and one representative confirmed tumor from multi-tumor donors. Both datasets are based on hg19, 1-based coordinates.

### 5 Higher sequence complementarity between persistent contacts; Repeat in the observed sequence complementarity of contacts

#### 5.1 Studies on the inherent property of double-stranded DNA to self-interact

##### DNA duplex self-assembly

*Inoue, S., Sugiyama, S., Travers, A. A., & Ohyama, T. (2007). Self-assembly of double-stranded DNA molecules at nanomolar concentrations. Biochemistry, 46(1), 164–171.* [24] *In vitro* work showing that double-stranded DNA molecules self-assemble in aqueous solutions containing physiological concentrations of $Mg^{2+}$, observed using atomic force microscopy. "DNA molecules preferentially interact with molecules with an identical sequence and length even in a solution composed of heterogeneous DNA species".

*Baldwin, G. S., Brooks, N. J., Robson, R. E., Wynveen, A., Goldar, A., Leikin, S., Seddon, J. M., & Kornyshev, A. A. (2008). DNA double helices recognize mutual sequence homology in a protein free environment. The journal of physical chemistry. B, 112(4), 1060–1064.* [25] Another *in vitro* work showing recognition and segregation of intact DNA duplexes (without any single-stranded elements) in a heterogeneous mixture of DNA sequences. Data is from imaging of fluorescently tagged DNA duplexes.

*Danilowicz, C., Lee, C. H., Kim, K., Hatch, K., Coljee, V. W., Kleckner, N., & Prentiss, M. (2009). Single molecule detection of direct, homologous, DNA/DNA pairing. Proceedings of the National Academy of Sciences of the United States of America, 106(47), 19824–19829.* [26] Another *in vitro* work using magnetic tweezers showing pairing of "double-stranded DNA molecules in the absence of proteins, divalent metal ions, crowding agents, or free DNA ends" (shown for regions of homology of 5 kb or more)

*Ohyama T. (2019). New Aspects of Magnesium Function: A Key Regulator in Nucleosome Self-Assembly, Chromatin Folding and Phase Separation. International journal of molecular sciences, 20(17), 4232.* [27] Review on this topic citing studies above.

##### DNA duplex self-assembly even in the presence of nucleosomes

*Nishikawa, J., & Ohyama, T. (2013). Selective association between nucleosomes with identical DNA sequences. Nucleic acids research, 41(3), 1544–1554.* [28] Authors used atomic force microscopy to show that the DNA duplex self-assembly, shown by studies above, can occur in the presence of nucleosomes.

##### *In vivo* support of direct recognition of sequence identity

*Gladyshev, E., & Kleckner, N. (2014). Direct recognition of homology between double helices of DNA in Neurospora crassa. Nature communications, 5, 3509.* [29] "*In vivo* study using *Neurospora crassa* suggesting the presence of the direct sequence recognition mechanism between identical DNA regions", “with the DNA segments aligning even in the absence of known pairing complexes involved in homology recognition”.

#### 5.2 Sequence complementarity and $c_p$

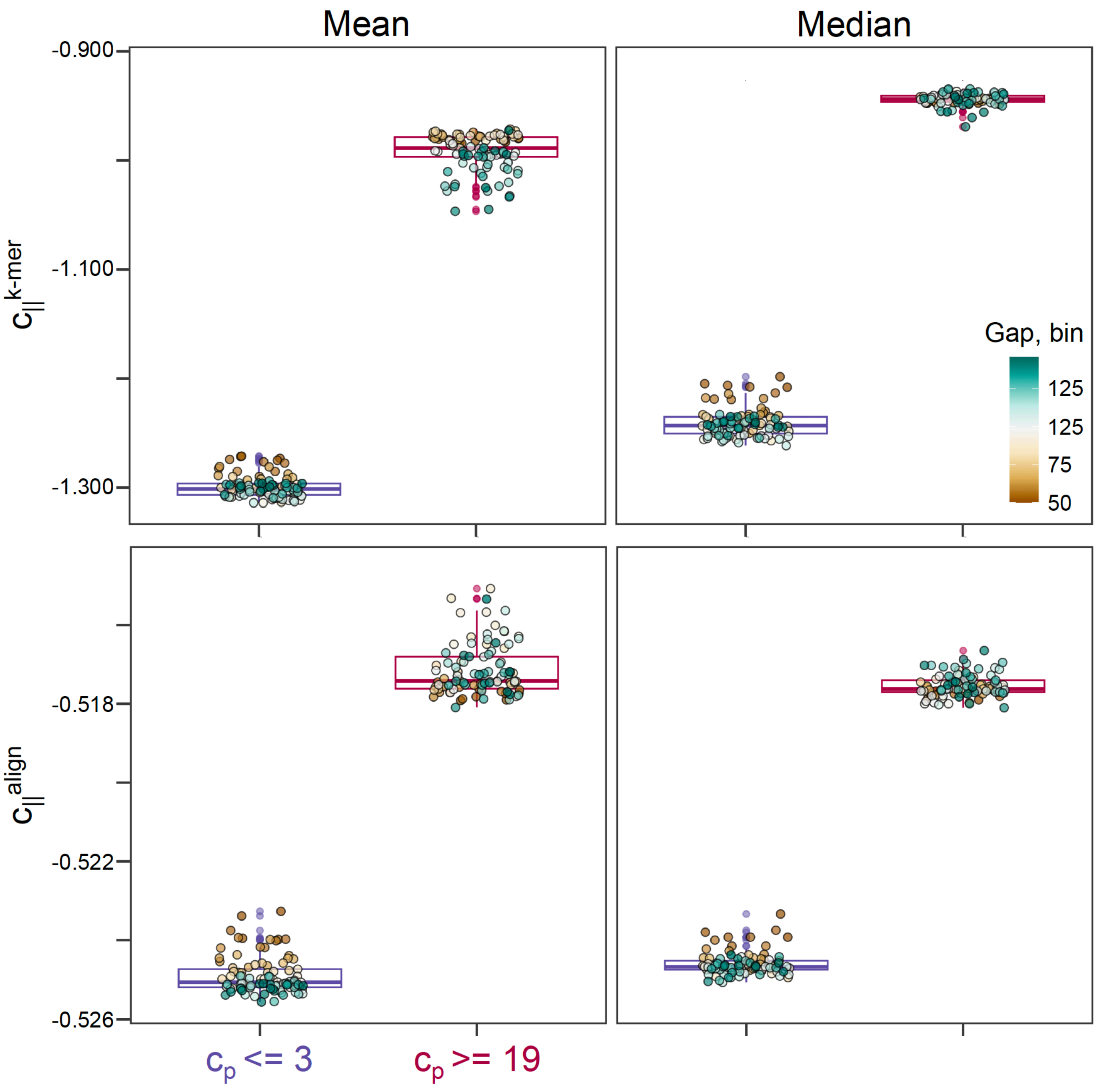

***Figure 23. Mean sequence complementarity of persistent vs. variable contacts across contact gaps.*** *Distributions of (**A**) $c_{||}^{k-mer}$ and (**B**) $c_{||}^{align}$ were compared at a given contact gap each with $\geq$ 100 persistent and variable contacts. All comparisons were significant (Mann-Whitney-Wilcoxon / MWW test, BH $P$ value $<$ 0.0001).*

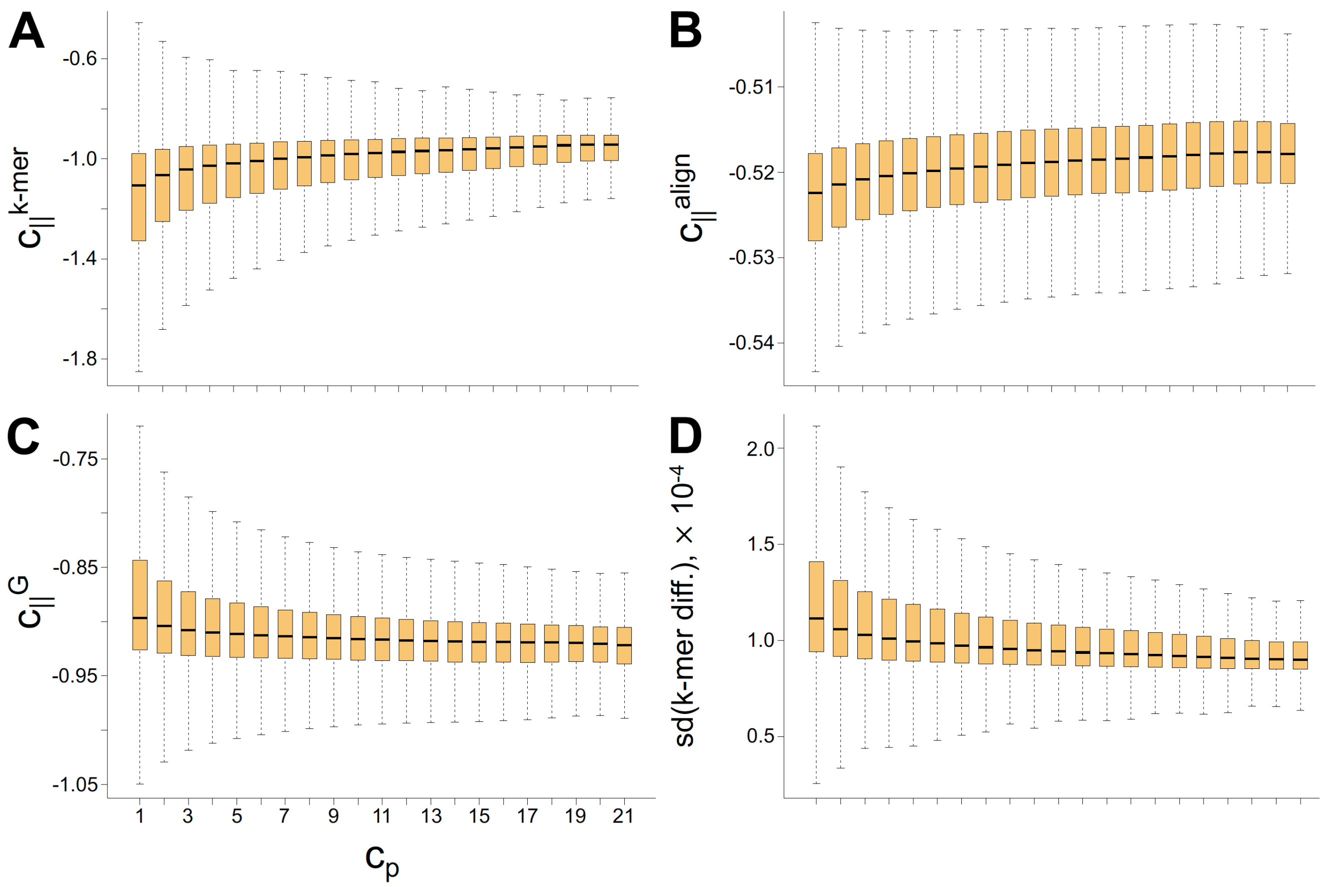

***Figure 24. Sequence complementarity ($c_{||}$) between contacting regions across $c_p$.*** *Sequence complementarity is based on (**A**) k-mer matching, $c_{||}^{k-mer}$, (**B**) global alignment using edit distance, $c_{||}^{align}$, and (**C**) hybridisation free energy, $c_{||}^{G}$ (see **Methods** for details). The latter, $c_{||}^{G}$, is the only metric that is expected to decrease in value for more similar sequences as observed from the plot. (**D**) shows the standard deviation (sd) of the differences in counts of the 7-mers for each contact per $c_p$ (k-mer diff.). The outliers are not shown though fully accounted for in statistical tests and inference of the boxplot metrics. Contacts formed by regions involving at least 1 missing bp were excluded. Directions of $c_{||}$ metrics and $c_p$ relationships based on Pearson and Spearman correlation analyses were consistent with the observed trends with absolute coefficients between 0.16 to 0.23 (p-values $< 2.2\times 10^{-16}$). One-way Analysis of Variance (ANOVA) and Kruskal-Wallis H test showed significant results for all metrics (p-values $< 2.2\times 10^{-16}$). Pairwise comparison of distributions between any two neighbouring $c_p$ values show significant differences (BH adj. p-value $< 0.01$) except for some comparisons between the highly persistent contacts ($c_p \geq 17$).*

**Global sequence alignment.** We used the open-source C/C$^{++}$ library edlib for the global alignment of contacting regions using edit distance, a metric quantifying the difference between two sequences in terms of the cost of transforming one sequence to another based on their alignment [30]. We specifically calculated the classical Levenshtein edit distance. With global alignment, gaps, in addition to substitutions, within or at the ends of the sequence are penalised by +1 regardless of base identity. For each contact, the calculation of edit distance considered possible combinations of all strands (in their correct 5'-3' direction), two from each $i$ and $j$ contact regions. In particular, edit distance was calculated between **1)** the plus strands of the contact regions, and **2)** the plus strand of one region and the reversed minus strand of its pair. The score for the complementarity was the smaller of the two resulting edit distances i.e. coming from the strand combination showing the better complementarity. The value was then scaled by the length of the contact regions. The few cases, in which the contact regions had different lengths (contacts formed by the ends of chromosomes), were excluded. The scaled value was negated to make the final alignment-based complementarity score, $c_{||}^{align}$, positively correlating with the degree of complementarity i.e. more similar sequences resulting in higher $c_{||}^{align}$. The R library `Rcpp` was used to interface with edlib in R. Note that all methods to measure $c_{||}$ excluded contacts formed by regions with at least one missing base pair.

**Matching of k-mer counts.** Sequence similarity quantification based on k-mer counts has long been used in sequence homology modelling and phylogeny reconstruction [31–35]. Here, for a given 40-kb region, the count of each $4^7$ possible 7-mers was determined considering both strands, sliding one base at a time. Heptamers were used taking into account the feasibility of the calculations for $4^7$ k-mers and the previously observed non-additive nature of DNA sequence properties while trying to represent them with k-mers shorter than 7-mers [36]. The k-mer-based complementarity score, $c_{||}^{k-mer}$, of a contact, is then given by the negative of the sum of the absolute differences between the 7-mer scaled counts of pairs of regions in contact. The value was similarly negated so that the final score was higher for more similar sequences. For each contact, the standard deviation of the scaled count differences was also obtained. Unlike $c_{||}^{align}$, the $c_{||}^{k-mer}$ of a contact between regions of different lengths can be calculated because the scaled count of a k-mer is a feature of a region not that of the whole contact.

**Crude estimation of hybridisation free energy.** Widely used programs that can be repurposed to calculate hybridisation free pseudo-energies of DNA duplexes are MultiRNAFold, UNAFold and ViennaRNA. In Tulpan et al. [37] (2010), it was shown that these methods perform comparably well, but given the high number and low resolution of our contacts requiring longer sequence spans, the use of any of those would be memory- and time-extensive. Instead, estimation was done using calculated free energy parameters for unique, perfectly matched DNA triplets reported in the same paper. They calculated these parameters using published, experimentally-derived free energies (measured at 37°C and 1 M sodium concentration) of fully complementary DNA duplexes. In their model, the experimental free energy for a fully complementary duplex was the sum of the parametric values for each triplet multiplied by the count of the triplet. This method is still an inadequate estimator of the real hybridisation free energies because it ignores multiple conformations possible, along with potential mismatches, internal loops, and dangling ends. Therefore, the use of this metric in this work was mainly for observing the general trend across $c_p$. Its major advantage over the other metrics is that it could somehow differentiate the hybridisation abilities of the short spans depending on the base composition.

The $c_{||}^{k-mer}$ calculation provided the 7-mer scaled counts of contacting regions. In order to maintain the 7-meric granularity of sequence matching but to apply the triplet-based free energies from [37] to our set of contacts, 7-mer free energy parameters were derived from the triplet energy parameters by sliding a 3-bp window along the 7-mers and averaging the parametric values of triplets present. The hybridisation free energy complementarity score, $c_{||}^{G}$, of a contact is then given by the sum of the products of the 7-mer parametric values with the matching 7-mer counts of regions in contact. This score was not negated i.e. more similar sequences would have a more negative $c_{||}^{G}$ value so as to reflect the directionality of the change in hybridisation free energy.

#### 5.3 Pairwise shuffling of contacting regions

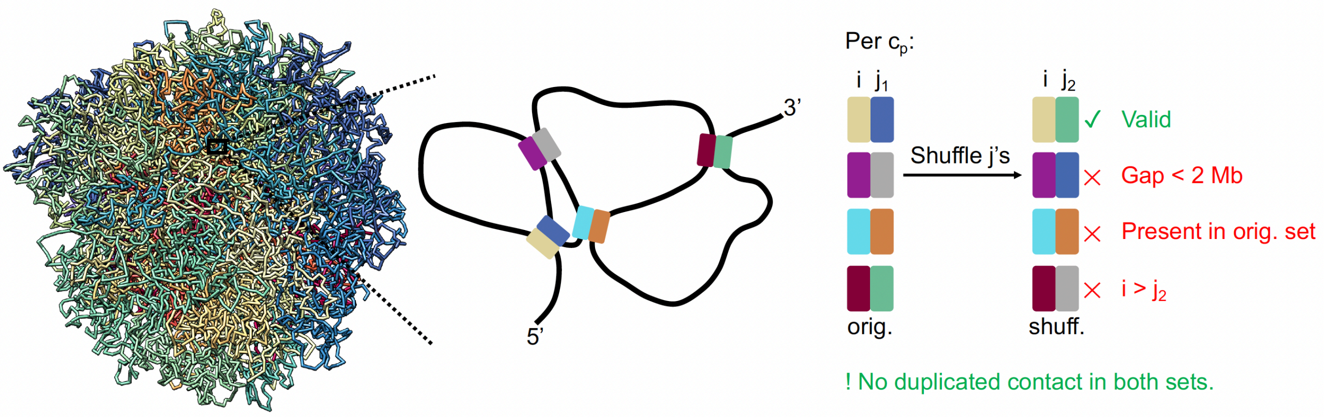

***Figure 25. Shuffling procedure to form fake contacts per $c_p$.*** *The real/original (orig.) set of 2-Mb contacts satisfies the following criteria:* ***1)*** *The set only contains unique, long-range contacts, $i$-$j_1$, with $i$ and $j_1$ regions always separated by a gap $\geq$ 2 Mb or 50 40-kb regions,* ***2)*** *$i$ is set to be always upstream of $j_1$ (i.e. $i < j_1$) to avoid duplicated contacts. Note that a region can take part in more than one contact. Shuffling, performed per chromosome and per $c_p$ category, was done using our general-purpose optimisation engine `ROptimus` [38, 39] in order to maximise the number of fake/shuffled (shuff.) contacts that would not be present in the real/orig. set but would comply with the aforementioned criteria (see* ***Methods*** *for details). The figure illustrates the conditions that should be satisfied by the shuffled contacts. Due to the required constraints on the shuffled set of contacts, the number of fake contacts per $c_p$ was less than that of the original set. However, for all chromosomes and $c_p$ values, the decrease in total number of contacts did not exceed 13% (****Figure 27*** *and* ***Table 4****).*

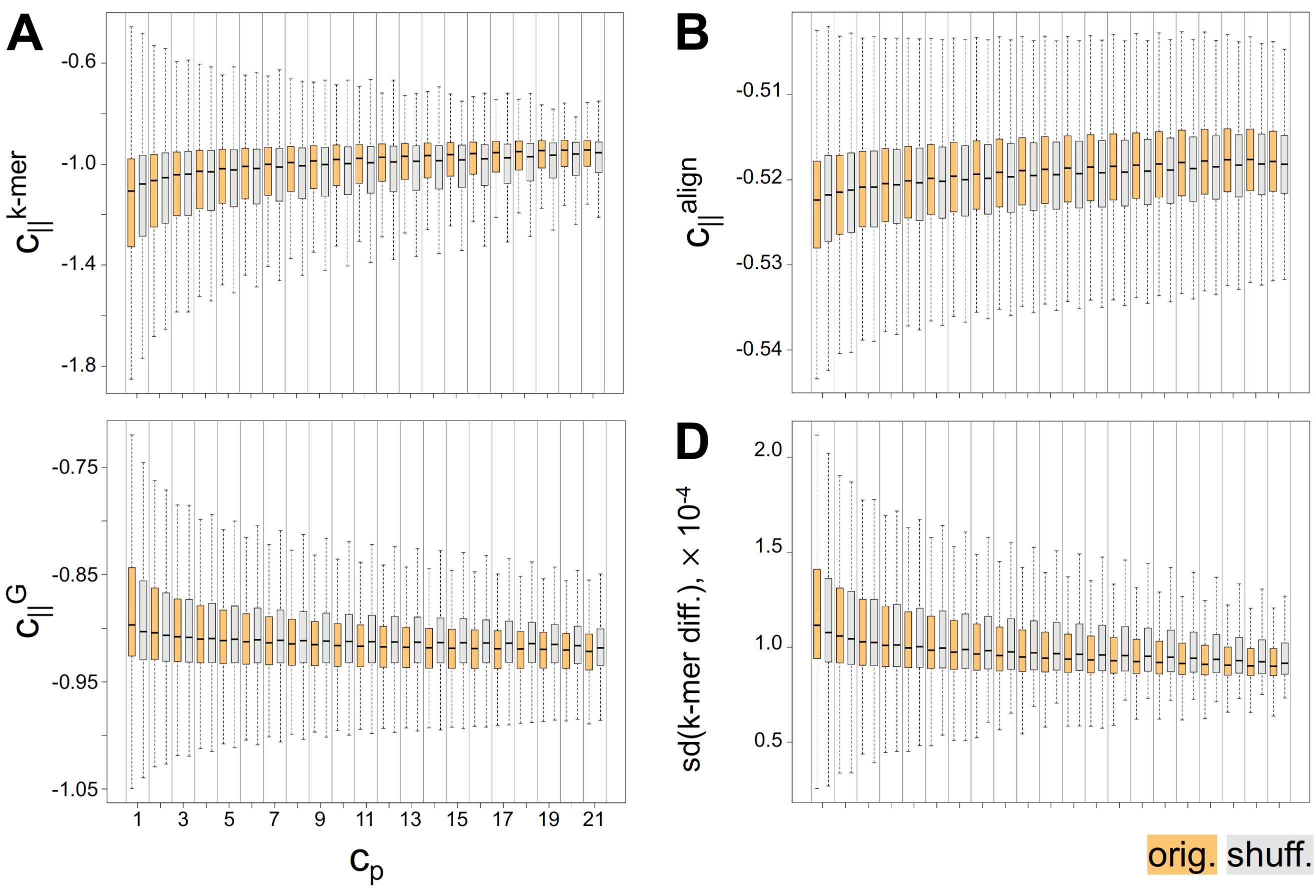

***Figure 26. Sequence complementarity ($c_{||}$) of original/real and shuffled/fake contacts across $c_p$.*** *Sequence complementarity is based on (**A**) k-mer matching, $c_{||}^{k-mer}$, (**B**) global alignment using edit distance, $c_{||}^{align}$, and (**C**) hybridisation free energy, $c_{||}^{G}$ (see **Methods** for details). The latter, $c_{||}^{G}$, is the only metric that is expected to decrease in value for more similar sequences as observed from the plot. (**D**) shows the standard deviation (sd) of the differences in counts of the 7-mers for each contact per $c_p$ (k-mer diff.). The outliers are not shown though fully accounted for in statistical tests and inference of the boxplot metrics. Contacts formed by regions involving at least 1 missing bp were excluded. Comparisons of complementarity distributions of original (orig., in yellow) and shuffled (shuff., in light grey) contacts show significant differences (BH adj. p-value $<$ 0.001).*

**Shuffling procedure using our R library `ROptimus`.** Given original (orig.) contacts between $i$ and $j_1$ regions, where $i > j_1$, the $j_1$ regions were shuffled once and then inputted to `Optimus()` from our R library `ROptimus` [39]. For each iteration, `Optimus()` altered the data by randomly taking two $j_1$ regions and exchanging them, followed by the calculation of the pseudo-energy and decision on the acceptance of the move based on the optimisation regimen. `Optimus()` was set to perform 12 M iterations while annealing the default acceptance ratios in 3 cycles and to produce one set of fake or shuffled (shuff.) contacts, $i$-$j_2$, per chromosome per $c_p$.

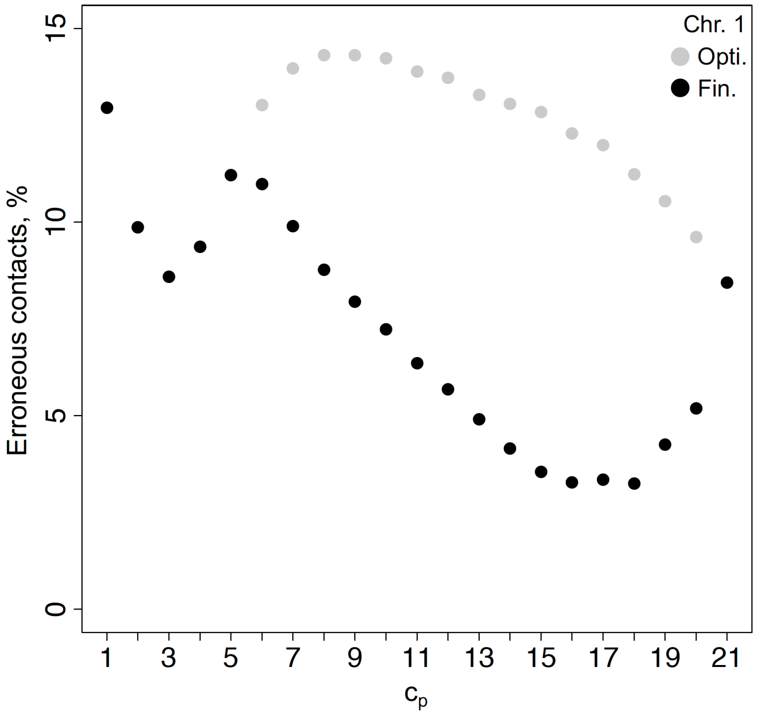

***Figure 27. Proportion of erroneous contacts in the chromosome 1 shuffled set of contacts** as percentage relative to the number of original contacts per $c_p$. The % Opti. (ROptimus) is the proportion of invalid contacts in the `Optimus()` output as described in **Figure** 25. To decrease this error rate, the regions in shuffled contacts with negative contact gaps (i.e. $i > j_1$) were flipped and the proportion of valid shuffled contacts was recalculated. The flipping was only accepted when it resulted in a valid contact, hence decreasing the error rate. The % Fin. (Final) is the error rate of the final shuffled set used for the subsequent analyses.*

***Table 4. Maximum error rates of shuffled sets of contacts per chromosome.***

| Max. erroneous contacts, % | | | | | |
|---|---|---|---|---|---|
| **Chr.** | **Opti.** | **Fin.** | **Chr.** | **Opti.** | **Fin.** |
| 1 | 14.32 | 12.96 | 13 | 11.23 | 10.23 |
| 2 | 15.17 | 12.38 | 14 | 9.89 | 9.89 |
| 3 | 13.24 | 11.70 | 15 | 9.87 | 9.71 |
| 4 | 13.87 | 11.90 | 16 | 10.73 | 10.73 |
| 5 | 13.19 | 11.43 | 17 | 10.34 | 10.34 |
| 6 | 12.56 | 11.29 | 18 | 10.68 | 10.68 |
| 7 | 11.58 | 11.26 | 19 | 11.24 | 11.24 |
| 8 | 11.83 | 10.68 | 20 | 10.49 | 10.49 |
| 9 | 10.77 | 10.77 | 21 | 12.20 | 12.20 |
| 10 | 11.04 | 10.19 | 22 | 11.76 | 9.54 |
| 11 | 11.15 | 10.43 | X | 12.34 | 12.34 |
| 12 | 11.20 | 10.63 | | | |

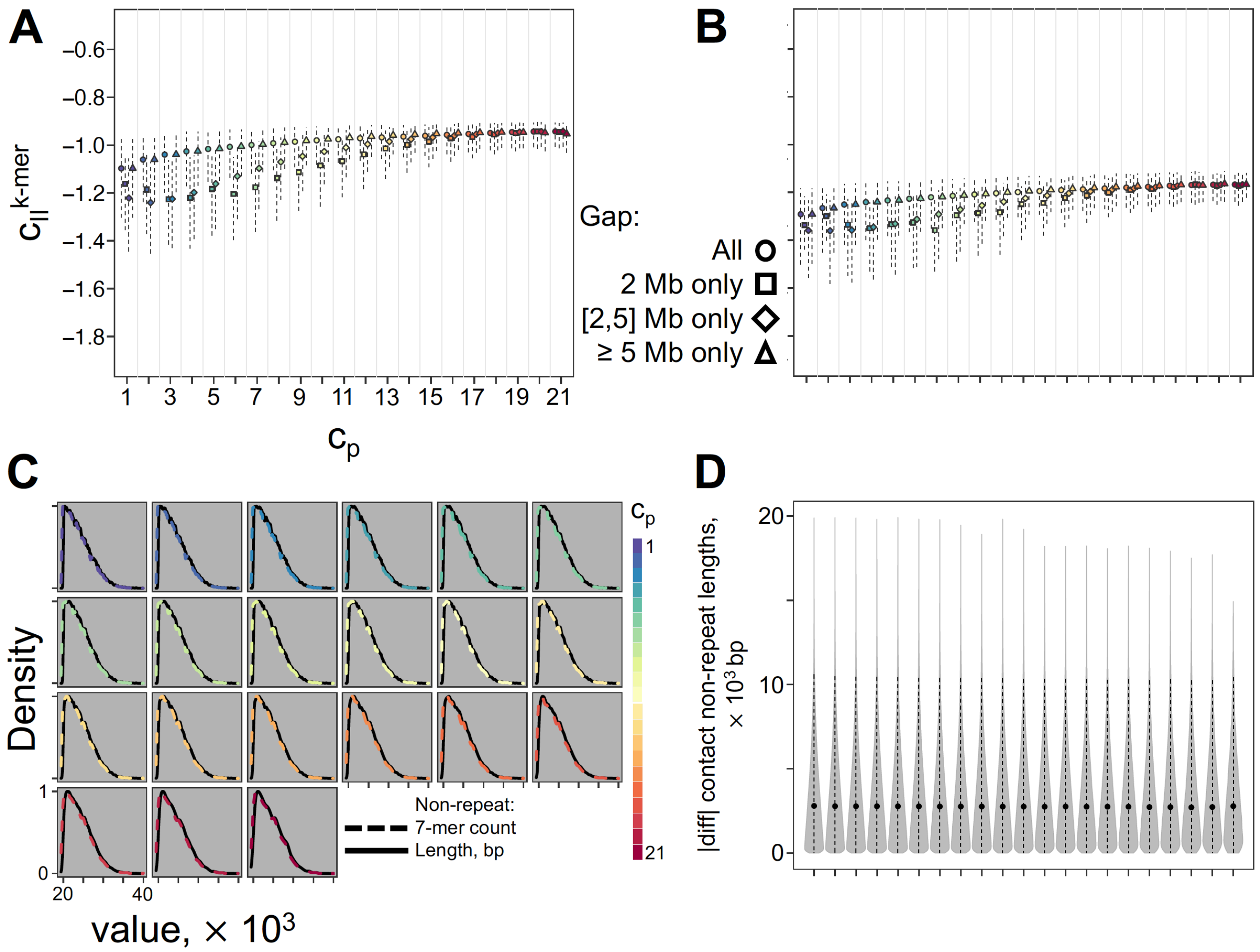

***Figure 28. Sequence complementarity ($c_{||}$) between contacting regions across $c_p$, minimising contact gap variation across $c_p$ and using repeat-masked human genome.*** *(**A-B**) Sequence complementarity shown is based on k-mer matching, $c_{||}^{k-mer}$. Points are located at the median and the dashed segments extend to the $25^{th}$ and $75^{th}$ percentiles. Shown are the $c_{||}^{k-mer}$ distribution for all long-range contacts (square), and for those with contact gap within the closed range i.e. 2-4 Mb (circle) and exactly 2 Mb (triangle) derived using (**A**) unmasked and (**B**) repeat-masked human genome. By using the repeat-masked genome, low complexity and interspersed repeats were ignored in the calculation of 7-mer counts such that a 7-mer with at least one repeat base pair was not counted. The counts were then divided by the total non-repeat sequence length. Contacts formed by regions with $>$ 50% of their sequence masked were excluded. In addition, those involving regions with at least one missing base pair in the unmasked genome were excluded also to be consistent with the analysis using the unmasked genome. Upon filtering based on repeat content of regions, about 21 to 35% of original contacts per $c_p$ (with the proportion increasing with $c_p$) remained valid for analysis. Pearson and Spearman correlation analyses show positive coefficients between 0.20 to 0.40 (p-values $< 2.2\times 10^{-16}$). One-way ANOVA and Kruskal-Wallis H test showed significant results for all metrics (p-values $< 2.2\times 10^{-16}$). Pairwise comparisons of the 5 most persistent and 5 most variable distributions show significant differences (BH adj. p-value $< 0.05$). (**C-D**) The following are plots assessing the effect of repeat masking on the contact regions. (**C**) compares the distribution of total non-repeat length (black solid line) and total 7-mer count of contact regions (non-black dashed line) across $c_p$. Duplicates of contact regions per $c_p$ were not removed to reflect the number of contacts formed by each region. (**D**) shows the absolute difference between the non-repeat lengths of two regions in contact across $c_p$ values. Points are located at the median and the dashed segments extend to the lower and upper whiskers of a default boxplot if drawn. The $c_{||}^{align}$ values were not calculated using the repeat-masked genome because, unlike the other measures, the applied method of scaling $c_{||}^{align}$ by the region length needs regions in contact to be of the same length.*

### 6 Additional analyses

#### 6.1 Molecular dynamics simulation to model the 3D genome using $c_{||}$ restraints

Although we have not obtained finalised contact maps usable for further investigations, we described below the simulations based on sequence complementarity performed by Dr. Zahra Fahmi with the supervision of Prof. Rosana Collepardo-Guevara. We used two kinds of bead-spring polymer models to simulate the folding of human chr. 1, where each bead represented a 40-kb region. The first model was a simple polymer model, where consecutive beads were connected by harmonic bonds and the Lennard-Jones potential governed interactions of non-bonded beads. The second model was a self-repulsive polymer model based on the MiChroM model [40], where a soft repulsive potential derived the interactions of non-bonded beads. Both models were informed by discretised $c_{||}$ scores through additional Lennard-Jones terms between non-bonded beads. Since the $c_{||}$ scores do not measure the actual complementarity degree or hybridisation energy, we initially opted to use the binned version of $c_{||}^{k-mer}$ (**Figure 29**) used in the Hi-$C_{||}$ maps. Contacts assigned to the top (top 5% contacts with highest $c_{||}$), rest (not belonging to top or bottom groups) and bottom (bottom 5% contacts with lowest $c_{||}$) groups were assigned interactions of strong, moderate, and weak strengths, respectively. **Figure 30** shows the contact maps obtained from the simulations.

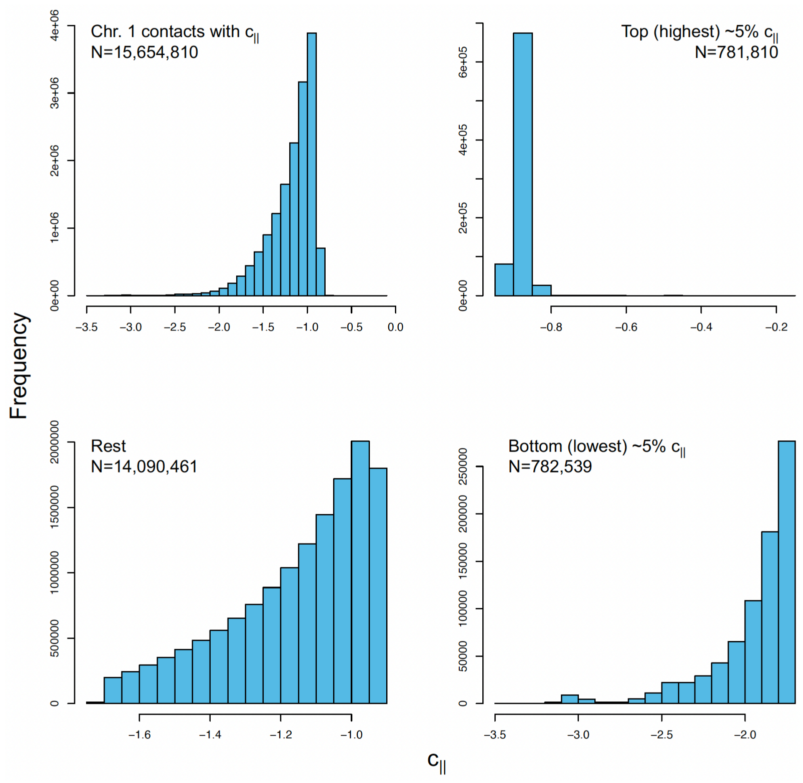

***Figure 29. Chromosome 1 contacts grouped based on** $c_{||}$. The grouping considered all possible contacts of chr. 1 (at 40-kb resolution), both short-range and long-range, with $c_{||}^{k-mer}$ value (15,654,810 contacts out of 19,415,796 possible chr. 1 contacts). The top and bottom groups include the top and bottom 5% of contacts with the highest and lowest $c_{||}^{k-mer}$, respectively. The remaining ones comprise the rest group. N refers to the count of contacts per plot or group.*

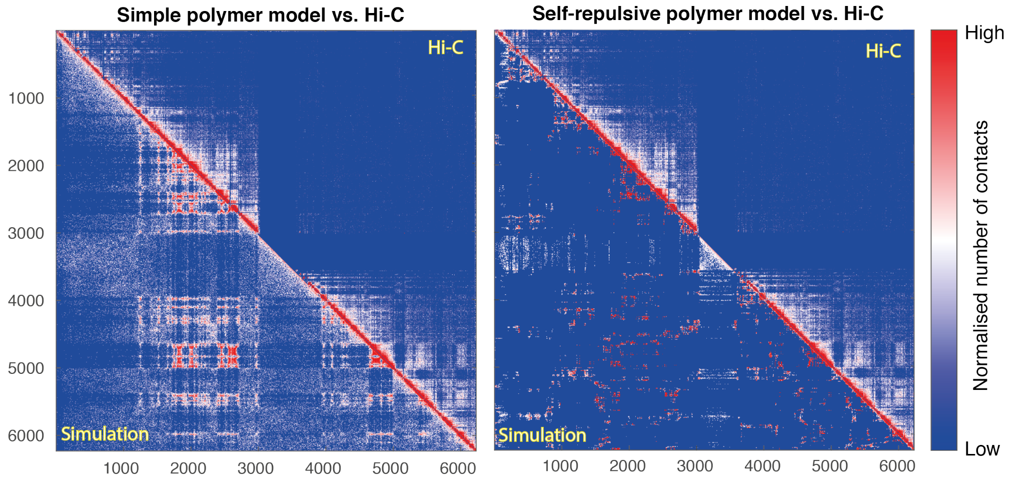

***Figure 30. Comparison of simulated and experimental chromosome 1 contact maps.*** *The upper triangles show the experimental Hi-C$_f$ map, which is the summation of the 21 Hi-C data sets used in this study [1] (upper triangles). The lower triangles show the contact maps from the simulations (Hi-C$_{sim}$) based on the simple polymer model (left) and the self-avoiding polymer model (right). For both models, each simulation was run to collect more than 500 data points that were sampled every 500,000 time steps throughout the simulation. To make Hi-C$_{sim}$ and Hi-C$_f$ comparable, the main diagonal line was set to zero. To normalise all the contact maps, every element was divided by the total number of contacts. Simulation trajectories were analysed in Python 3.5 using MDAnalysis library. All figures were plotted by MATLAB R2021b.*

#### 6.2 Associations with known motifs and *de novo* sequences

To check whether contact persistence is associated with any sequence motif, we determined the enrichment of **1)** known vertebrate motifs (**Figure 31**) and **2)** *de novo* motifs derived from sequences of $c_p = 21$ contact regions, where any effect should be the most pronounced.

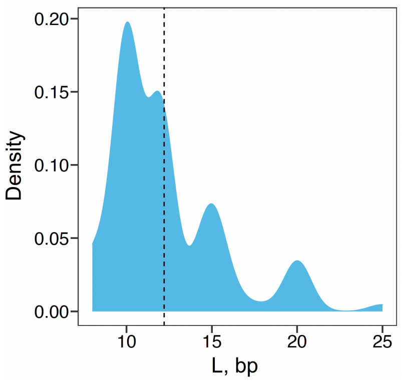

***Figure 31. Lengths (L) of 426* HOMER *vertebrate motifs used in the known motif analysis.*** *Values range from 8 to 25 bp, with the mean value at around 12 bp (dashed line).*

The set of unique contact regions with $c_p \geq 1$ was used as background for the analyses. Multiple runs were done each using 50, 60, and 70% of the target and background regions (randomly sampled). Although it is possible to limit the searching area for motifs in HOMER, the whole 40-kb span of each region was scanned in this case. For *de novo* discovery, the search length for the new motifs was limited to within 8 to 12 bp. The top five most enriched new motifs per length were returned for further consideration.

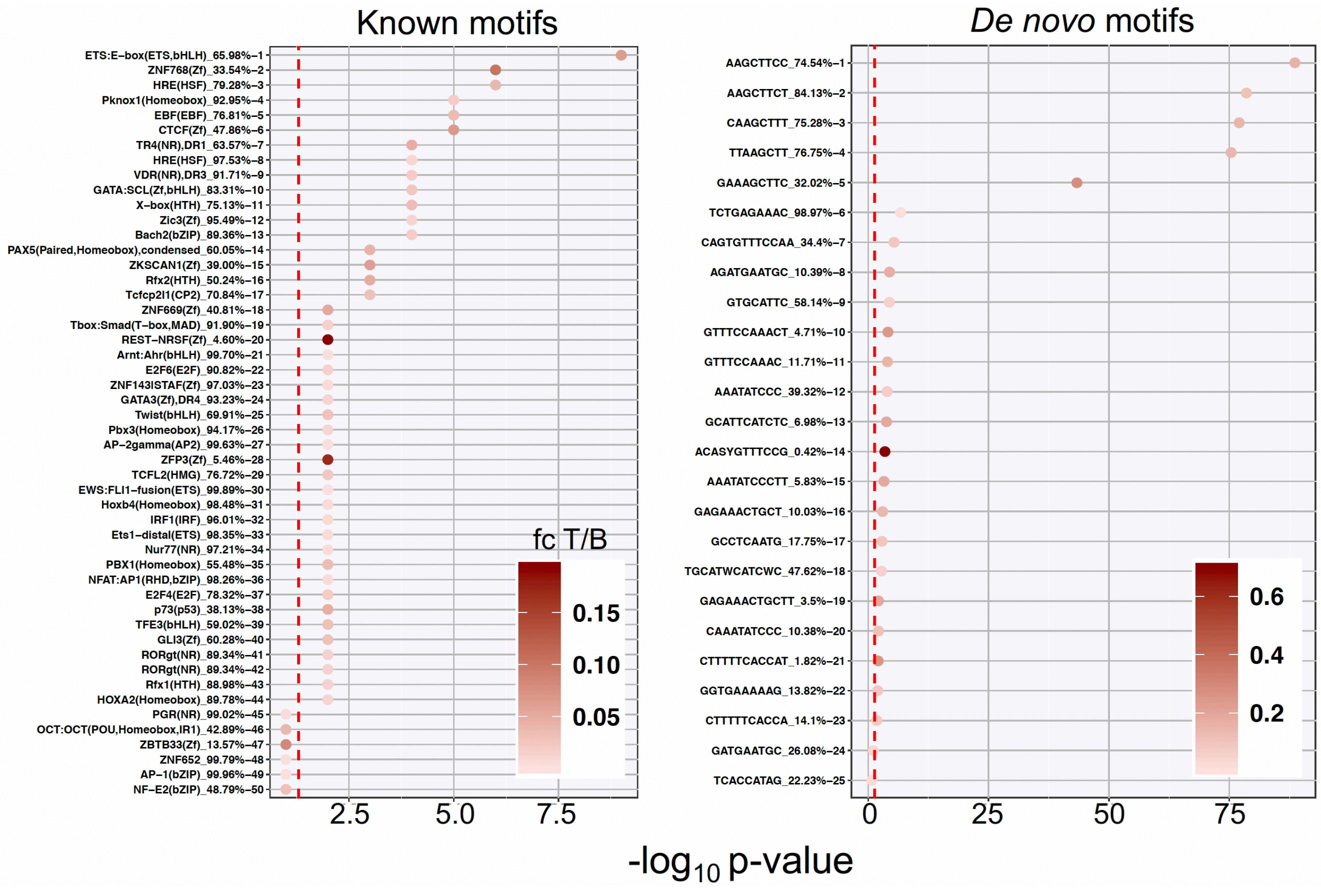

***Figure 32. Enrichment of known and de novo motifs at prime contact ($c_p = 21$) regions*** *against the set of all unique contact ($c_p \geq 1$) regions. Of the total number of target, T, and background, B, regions, 70% were used (randomly sampled) due to memory limitations. Before sampling, regions with at least one missing bp were removed, and out of the remaining regions, there were a few excluded by the* HOMER *algorithm based from the returned number of regions per set (9121 target and 42,979 background regions). Percentage values on the y-axis label is the proportion of background regions containing the motif. The colour of the points indicates the fold change (fc) of the proportion of target regions with the motif relative to that of the background. The red dashed line marks the significance threshold at -log*$_{10}$ *0.05* $\approx$ *1.3010. The analysis was done via the interfacing function* `find_motifs_genome()` *from the R library* `marge`*.*

We focused on the most significantly enriched motifs, with p-values $\leq 10^{-50}$, as recommended on the HOMER manual (`http://homer.ucsd.edu/homer/ngs/peakMotifs.html`). These include four 8-bp *de novo* motifs, which were used in the enrichment analyses in (**Subsection 4.1**). Overall, given the short lengths of the motifs studied here and the increasing amount of Hi-C data at higher than 40-kb resolution, this matter can be revisited in the future.

#### 6.3 Enrichment of genes across contacts of varying $c_p$

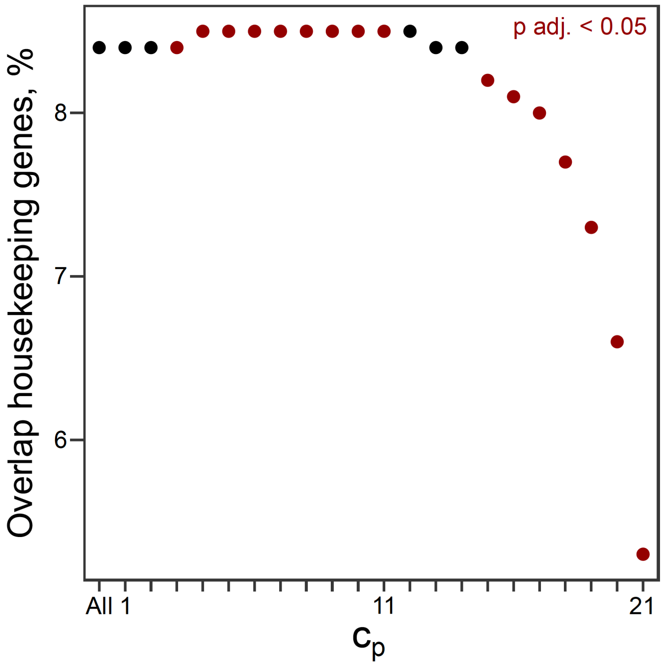

***Figure 33. Fraction of housekeeping genes overlapping with contacts across $c_p$.*** *List of 2176 housekeeping human genes stably expressed across 52 tissues and cells types was downloaded from HRT Atlas v1.0 database [41]. The significance of overlap, Benjamini-Hochberg (BH) adjusted (adj.), was derived via permutation test, using the set of unique genes that overlap with all long-range contacts ("All") as control set. For each set of genes per $c_p$, 10,000 gene sets of the same size were randomly sampled without replacement from the control set to generate the null distribution.*

### 7 Supplementary Methods

#### 7.1 Computational platforms and resources

The computational analyses were mainly done on the high-performance computing facilities of the Medical Research Council (MRC) Weatherall Institute of Molecular Medicine, University of Oxford, utilising cluster nodes with 256 GB random access memory, and Intel Xeon E5-2680v3 12-core (24-thread) and Intel Xeon E7-8891v3 10-core (20-thread) processors. Code was mostly written in the R programming language [42] through RStudio, an Integrated Development Environment for R [43]. The molecular dynamics (MD) simulations for the polymer models of chromatin were performed on the computing facilities at the University of Cambridge (CSD3). Each simulation was run on a single node with 32 CPUs and 6 GB of memory per CPU. Results of simulations were analysed using Python [44] and MATLAB [45] programming languages.

#### 7.2 R packages

The following are the R packages used for the computational analyses. The specific versions are indicated, if applicable. `BiocManager` (1.30.16) [46], `Biostrings` (2.58.0) [47], `BSgenome` (1.58.0) [48], `cluster` (2.1.2) [49], `clusterProfiler` (3.18.1) [50], `compiler` (4.0.5) [42], `ComplexHeatmap` (2.6.2) [51], `cowplot` (1.1.1) [52], `data.table` (1.14.0) [53], `devtools` (2.4.2) [54], `doParallel` (1.0.16) [55], `dplyr` (1.0.7) [56], `DT` (0.18) [57], `expm` (0.999-6) [58], `factoextra` (1.0.7) [59], `foreach` (1.5.1) [60], `GenomicRanges` (1.42.0) [61], `ggnewscale` (0.4.5) [62], `ggplot2` (3.3.5) [63], `ggpubr` (0.4.0) [64], `ggrepel` (0.9.1) [65], `ggsci` (2.9) [66], `ggseqlogo` (0.1) [67], `gplots` (3.1.1) [68], `grDevices` (4.0.5) [42], `grid` (4.0.5) [42], `gridExtra` (2.3) [69], `gridGraphics` (0.5-1) [70], `hexbin` (1.28.2) [71], `Hmisc` (4.5-0) [72], `IRanges` (2.24.1) [61], `itertools` (0.1-3) [73], `marge` (0.0.4.9999) [74], `methods` (4.0.5) [42], `np` (0.60-11) [75], `Optimus` [38, 39], `org.Hs.eg.db` (3.12.0) [76], `R4RNA` (1.18.0) [77, 78], `RColorBrewer` (1.1-2) [79], `Rcpp` (1.0.7) [6, 80, 81], `readxl` (1.3.1) [82], `regioneR` (1.22.0) [83], `reshape` (0.8.8) [84], `reshape2` (1.4.4) [84], `rtracklayer` (1.50.0) [85], `scales` (1.1.1) [86], `scuttle` (1.12.0) [7], `shiny` (1.6.0) [87], `shinydashboard` (0.7.1) [88], `shinyWidgets` (0.6.0) [89], `sna` (2.6) [90], `splines` (4.0.5) [42], `stats` (4.0.5) [42], `stringr` (1.4.0) [91], `utils` (4.0.5) [42], `viridis` (0.6.1) [92], `visNetwork` (2.0.9) [93], and `yarrr` (0.1.5) [94].

#### 7.3 Additional software

- DAVID (The Database for Annotation, Visualization and Integrated Discovery, v6.8) [18, 19] https://david.ncifcrf.gov
- edlib (v1.2.6) [30]

https://github.com/Martinsos/edlib

- GO (Gene Ontology, July 2020 release) [95, 96]

  https://geneontology.org

- HOMER (v4.10) [97]

  https://homer.ucsd.edu/homer

- Juicer (v1.9.9, CUDA v8.0) [98]

  https://github.com/aidenlab/juicer

- KEGG (Kyoto Encyclopedia of Genes and Genomes, v94.2 release) [99–101]

  https://www.genome.jp/kegg

- LAMMPS [102]

  https://github.com/lammps/lammps

- MATLAB (R2021b) [45]

  https://matlab.mathworks.com/

- `MDAnalysis` (v0.20.1, Python package) [103, 104]

  https://github.com/MDAnalysis/mdanalysis

- Python (v3.5) [44]

  https://www.python.org/

#### 7.4 Outsourced data

This thesis is a purely computational work that relied on these publicly available datasets:

- Bed files of $\approx 300$ *H. sapiens* features downloaded mainly from the Encyclopedia of DNA Elements (ENCODE) [105, 106] and National Institutes of Health (NIH) Roadmap Epigenomics Project [107] (refer to **Table 7** for the detailed list)

- *D. melanogaster* dm6 genome [108, 109] (GCA_000001215.4, NCBI Assembly database [110])

  https://www.ncbi.nlm.nih.gov/assembly/GCF_000001215.4

- *H. sapiens* GRCh37 (hg19) genome unmasked and repeat-masked versions (Ensembl Release 73 September 2013) [111, 112]

  https://ftp.ensembl.org/pub/release-73/fasta/homo_sapiens/dna

- *H. sapiens* GRCh38 (hg38) genome unmasked version (Ensembl Release 108 October 2022) [**cunningham_ensembl_2022**]

  http://ftp.ensembl.org/pub/release-108/fasta/homo_sapiens/dna

- *H. sapiens* genes/gene predictions and repeat annotations (Feb. 2009 GRCh37/hg19, UCSC Table Browser data retrieval tool [113])

  https://genome.ucsc.edu/cgi-bin/hgTables

- *H. sapiens* baseline expression datasets (E-MTAB-1733 and E-MTAB-5214) [16, 17] from the EMBL-EBI Expression Atlas [15])

  https://www.ebi.ac.uk/gxa/home

- Hi-C contact data and TAD boundaries from 21 *H. sapiens* cell lines and tissues (GEO:GSE87112) [1]

  https://www.ncbi.nlm.nih.gov/geo/query/acc.cgi?acc=GSE87112

- Hi-C contact data of *H. sapiens* IMR-90 fetal lung fibroblast cell line (GEO:GSE63525) [12]

  https://www.ncbi.nlm.nih.gov/geo/query/acc.cgi?acc=GSE63525

- Hi-C contact data of *H. sapiens* H1 human embryonic stem cell line (4DNFIQYQWPF5) [114]

  https://data.4dnucleome.org/files-processed/4DNFIQYQWPF5

- Hi-C contact data (.hic) from 2 *D. melanogaster* cell lines (GEO:GSE122603) [115]

  *Email correspondence with authors*

- Phylo-HMRF evolutionary states data (GEO:GSE128800) [10]

  https://www.ncbi.nlm.nih.gov/geo/query/acc.cgi?acc=GSE128800 and *email correspondence with authors for undeposited data*

- Replication timing data from ReplicationDomain [20]

  http://www.replicationdomain.org

- Somatic cancer SNV data from ICGC Data Portal (Release 28, 2019 March 27) [21]

  https://dcc.icgc.org

- TAD data from 2 *D. melanogaster* cell lines (GEO:GSE122603) [115]

  https://www.ncbi.nlm.nih.gov/geo/query/acc.cgi?acc=GSE122603

- TAD boundaries from 2 *D. melanogaster* cell lines (GEO:GSE122603) [115]

  https://www.ncbi.nlm.nih.gov/geo/query/acc.cgi?acc=GSE122603

- Triplet-based free energy parameters [37]

  https://doi.org/10.1186/1471-2105-11-105https://doi.org/10.1186/1471-2105-11-105

#### 7.5 Default calculations, processing and settings

**Boxplots.** Lower and upper whiskers of boxplots were set to extend to $Q_1 - 1.5\times$IQR and $Q_3 + 1.5\times$IQR, where $Q_1$, $Q_3$ and IQR are the $25^{th}$ and $75^{th}$ percentiles, and the interquartile range, respectively.

**Contact gap.** This is equal to the total number of base pairs between regions in contact, excluding any bp from within the regions. For example, given a contact resolution of 40 kb, the gap between region 1 (1 to 40,000 bp) and region 3 (80,001 to 120,000 bp) is the length of sequence counting from the $40{,}0001^{st}$ bp to $80{,}000^{th}$ bp, which is equal to 40,000 bp. In the text, the contact gap is either expressed in bp or the number of regions/bins of length equal to the contact resolution.

**Correlation tests.** Correlation tests were done using the R function `cor.test()`, providing correlation coefficients and p-values.

**Fold change.** This was calculated by taking the $log_2$ of the ratio of a value to the reference value.

**Gene transcript representative.** In analyses using gene coordinates, genes with multiple transcripts were represented either by the single longest transcript or, in the case of ties, by any of the longest transcripts preferring coding over non-coding ones. The latter was the case for 962 genes out of 24,910 ($\approx$ 3.86%) unique genes from the UCSC hg19 gene annotation table.

**Genomic coordinate conversion between genomes.** Conversion between different versions of reference genomes was done using `liftOver()` from the R library `rtracklayer`.

**Overlap of genomic regions.** Having an overlap means having at least 1 shared or common bp.

**Statistical significance tests.** Pairwise comparisons of $> 2$ distributions were done using pairwise implementations of the Student's t-test (no assumption of equal variances) and Mann-Whitney-Wilcoxon (MWW) test *via* R functions `paired.t.test()` and `paired.wilcox.test()`, respectively. For both methods, the alternative hypothesis was two-sided and the p-values reported were Benjamini-Hochberg (BH) adjusted (adj.) [116]. Student's t-test (no assumption of equal variances) and Mann-Whitney-Wilcoxon test for comparing two distributions were implemented using the R functions `t.test()` and `wilcox.text()`, respectively. Two-sided Pearson, Spearman and Mann-Kendall correlation tests were done *via* `cor.test(exact=FALSE)`. The alpha ($\alpha$) or significance level is by default 0.05.

### 8 Appendix

***Table 5. Counts of long-range contacts per cell line or tissue*** *(CT) for minimum contact gaps, (**A**) 2 and (**B**) 0.5 Mb. "All$c_p$" and "AllCT" considered all $c_p$ values, and cell lines and tissues, respectively.*

**(A)**

| | Cell line / Tissue (CT) | | | | | | | | | | | | | | | | | | | | |
|---|---|---|---|---|---|---|---|---|---|---|---|---|---|---|---|---|---|---|---|---|---|
| $c_p$ | **Co** | **Hi** | **Lu** | **LV** | **RV** | **Ao** | **PM** | **Pa** | **Sp** | **Li** | **SB** | **AG** | **Ov** | **Bl** | **MesC** | **MSC** | **NPC** | **TLC** | **ESC** | **FC** | **LC** | **AllCT** |
| **1** | 499162 | 615617 | 207850 | 1014763 | 879294 | 1446850 | 277425 | 484743 | 553037 | 2086326 | 501904 | 434599 | 497241 | 629713 | 984988 | 865600 | 651224 | 795325 | 532906 | 3172810 | 2561138 | 19692515 |
| **2** | 1018967 | 1233154 | 419892 | 1994768 | 1752084 | 2926063 | 587809 | 986699 | 1061490 | 4053866 | 1048887 | 921180 | 1005012 | 1280177 | 1993362 | 1879584 | 1216792 | 1606805 | 1112595 | 5739661 | 4643105 | 19240976 |
| **3** | 1349945 | 1607948 | 547883 | 2570532 | 2277498 | 3841861 | 806366 | 1284658 | 1330895 | 5096238 | 1385558 | 1237182 | 1305939 | 1685758 | 2641388 | 2694503 | 1481650 | 2114880 | 1524257 | 6882505 | 5546374 | 16404606 |
| **4** | 1485504 | 1743529 | 594003 | 2759269 | 2465361 | 4176006 | 912252 | 1379546 | 1387720 | 5318960 | 1500833 | 1366150 | 1408961 | 1844418 | 2948443 | 3224886 | 1512331 | 2328657 | 1765146 | 6948405 | 5573180 | 13160890 |
| **5** | 1498842 | 1733739 | 591722 | 2724034 | 2443850 | 4133784 | 943850 | 1353810 | 1338009 | 5070105 | 1478887 | 1373200 | 1398353 | 1841996 | 3039262 | 3481032 | 1435856 | 2360781 | 1904019 | 6468592 | 5186822 | 10360109 |
| **6** | 1449372 | 1656581 | 566806 | 2579657 | 2320709 | 3890454 | 927075 | 1276685 | 1239886 | 4624568 | 1395707 | 1323548 | 1334877 | 1757030 | 2995111 | 3509232 | 1328404 | 2291694 | 1981135 | 5757506 | 4645645 | 8141947 |
| **7** | 1365693 | 1543780 | 530359 | 2383928 | 2149936 | 3533949 | 891437 | 1180406 | 1130331 | 4097772 | 1287739 | 1243965 | 1248090 | 1635642 | 2850188 | 3351717 | 1219093 | 2167226 | 1989809 | 4971201 | 4061794 | 6404865 |
| **8** | 1261473 | 1413631 | 489964 | 2151244 | 1950165 | 3109803 | 832886 | 1076885 | 1015909 | 3530500 | 1162852 | 1148427 | 1144226 | 1490137 | 2617296 | 3052059 | 1108997 | 1996718 | 1931645 | 4171062 | 3467097 | 5015372 |
| **9** | 1139925 | 1263485 | 443635 | 1890114 | 1722928 | 2647421 | 759837 | 961077 | 897966 | 2955345 | 1028133 | 1034727 | 1027270 | 1320223 | 2314132 | 2657402 | 991507 | 1787176 | 1794476 | 3395218 | 2883773 | 3879530 |
| **10** | 1005698 | 1103909 | 393508 | 1616678 | 1481646 | 2182974 | 675251 | 841553 | 778110 | 2402544 | 888089 | 909747 | 894734 | 1144156 | 1969088 | 2221305 | 869658 | 1554200 | 1596398 | 2683893 | 2333111 | 2954625 |
| **11** | 867794 | 945518 | 343580 | 1345074 | 1242582 | 1749748 | 587874 | 723058 | 662113 | 1901927 | 751785 | 783955 | 764250 | 967726 | 1619728 | 1795230 | 748194 | 1317614 | 1366145 | 2073019 | 1843880 | 2218254 |
| **12** | 733619 | 791516 | 295486 | 1092737 | 1016644 | 1368412 | 497690 | 608081 | 551377 | 1469315 | 622434 | 662108 | 635098 | 800341 | 1295033 | 1407348 | 636803 | 1085867 | 1130092 | 1565368 | 1424351 | 1640810 |
| **13** | 610397 | 650975 | 250600 | 870114 | 816489 | 1047761 | 414517 | 504431 | 452716 | 1112737 | 505991 | 548539 | 520302 | 651625 | 1011554 | 1078297 | 534187 | 875118 | 908862 | 1163246 | 1080993 | 1200727 |
| **14** | 498001 | 525773 | 209517 | 675318 | 640140 | 784932 | 339707 | 411872 | 366347 | 824099 | 405691 | 446747 | 416858 | 519706 | 768465 | 805815 | 441343 | 688166 | 707700 | 848643 | 803906 | 866339 |
| **15** | 400065 | 418139 | 176037 | 517796 | 495463 | 579586 | 274639 | 332789 | 293224 | 602328 | 322058 | 360419 | 331722 | 411173 | 574508 | 593783 | 361225 | 528811 | 541179 | 613340 | 590781 | 621271 |
| **16** | 308509 | 318969 | 143884 | 379951 | 366965 | 412408 | 217105 | 259189 | 227701 | 423660 | 249233 | 279511 | 255854 | 313970 | 411180 | 419858 | 284173 | 388191 | 393620 | 428091 | 417578 | 431225 |
| **17** | 226708 | 232616 | 114560 | 266758 | 259890 | 281399 | 166364 | 195226 | 172168 | 286528 | 185819 | 209508 | 191469 | 229695 | 281356 | 285031 | 213162 | 271548 | 273594 | 288001 | 283504 | 289112 |
| **18** | 153426 | 155898 | 86103 | 171715 | 169079 | 177202 | 119243 | 135694 | 120984 | 179129 | 128802 | 144184 | 132657 | 154500 | 177406 | 178537 | 146580 | 173741 | 174452 | 179463 | 178001 | 179822 |
| **19** | 90999 | 91894 | 58989 | 97717 | 96823 | 99204 | 76364 | 83964 | 76678 | 99693 | 79956 | 87571 | 82104 | 91552 | 99286 | 99587 | 88777 | 98321 | 98478 | 99777 | 99378 | 99848 |
| **20** | 43489 | 43651 | 34167 | 44924 | 44748 | 45140 | 39642 | 41966 | 39801 | 45205 | 40343 | 42771 | 41189 | 43678 | 45160 | 45198 | 43024 | 44998 | 45026 | 45220 | 45160 | 45225 |
| **21** | 13281 | 13281 | 13281 | 13281 | 13281 | 13281 | 13281 | 13281 | 13281 | 13281 | 13281 | 13281 | 13281 | 13281 | 13281 | 13281 | 13281 | 13281 | 13281 | 13281 | 13281 | 13281 |
| **All$c_p$** | 16020869 | 18103603 | 6511826 | 27160372 | 24605575 | 38448238 | 10360614 | 14135613 | 13709743 | 46194126 | 14983982 | 14571319 | 14649487 | 18826497 | 30650215 | 33659285 | 15326261 | 24489118 | 21784815 | 57508302 | 47682852 | 112861349 |

**(B)**

| | Cell line / Tissue (CT) | | | | | | | | | | | | | | | | | | | | |
|---|---|---|---|---|---|---|---|---|---|---|---|---|---|---|---|---|---|---|---|---|---|
| $c_p$ | **Co** | **Hi** | **Lu** | **LV** | **RV** | **Ao** | **PM** | **Pa** | **Sp** | **Li** | **SB** | **AG** | **Ov** | **Bl** | **MesC** | **MSC** | **NPC** | **TLC** | **ESC** | **FC** | **LC** | **AllCT** |
| **1** | 499366 | 615829 | 207908 | 1015325 | 879729 | 1447502 | 277527 | 484911 | 553256 | 2087608 | 502002 | 434740 | 497337 | 629957 | 985836 | 866527 | 651466 | 795839 | 533456 | 3174397 | 2563048 | 19703566 |
| **2** | 1019383 | 1233637 | 420027 | 1995786 | 1752908 | 2927263 | 587973 | 987100 | 1062015 | 4056245 | 1049068 | 921479 | 1005196 | 1280616 | 1995064 | 1881294 | 1217224 | 1607873 | 1113719 | 5742261 | 4646359 | 19251245 |
| **3** | 1350740 | 1608811 | 548142 | 2572119 | 2278839 | 3843754 | 806654 | 1285368 | 1331733 | 5100052 | 1385920 | 1237673 | 1306248 | 1686480 | 2644181 | 2697097 | 1482396 | 2116798 | 1526107 | 6886373 | 5551432 | 16415639 |
| **4** | 1486864 | 1745009 | 594430 | 2761825 | 2467548 | 4179040 | 912752 | 1380747 | 1389119 | 5324853 | 1501496 | 1367027 | 1409511 | 1845568 | 2952851 | 3228795 | 1513574 | 2331795 | 1768114 | 6954504 | 5580810 | 13174058 |
| **5** | 1501171 | 1736273 | 592446 | 2728034 | 2447244 | 4138501 | 944707 | 1355782 | 1340192 | 5078931 | 1479937 | 1374659 | 1399289 | 1843808 | 3046001 | 3487125 | 1437893 | 2365836 | 1908710 | 6477574 | 5197567 | 10376336 |
| **6** | 1452878 | 1660354 | 567895 | 2585432 | 2325736 | 3897493 | 928368 | 1279697 | 1243202 | 4636651 | 1397376 | 1325770 | 1336296 | 1759860 | 3004880 | 3517990 | 1331370 | 2299438 | 1988114 | 5770059 | 4660093 | 8161492 |
| **7** | 1371280 | 1549973 | 532115 | 2392776 | 2157804 | 3545049 | 893505 | 1185123 | 1135386 | 4115635 | 1290430 | 1247688 | 1250386 | 1640032 | 2865043 | 3365178 | 1223826 | 2179117 | 2000709 | 4989558 | 4082096 | 6430387 |
| **8** | 1269888 | 1422811 | 492613 | 2164309 | 1961785 | 3126153 | 836096 | 1083973 | 1023243 | 3555145 | 1166969 | 1154176 | 1147941 | 1496791 | 2638746 | 3071862 | 1116191 | 2014062 | 1947817 | 4196533 | 3494448 | 5047694 |
| **9** | 1152740 | 1277450 | 447847 | 1909737 | 1740195 | 2671728 | 765153 | 971734 | 908726 | 2990082 | 1034497 | 1043881 | 1033225 | 1330430 | 2345268 | 2686654 | 1002931 | 1812989 | 1818955 | 3430997 | 2921017 | 3921804 |
| **10** | 1024641 | 1124270 | 399669 | 1645105 | 1507183 | 2218409 | 683222 | 857080 | 793354 | 2449738 | 898116 | 923688 | 904156 | 1159722 | 2012701 | 2262944 | 886586 | 1591415 | 1632097 | 2732398 | 2382706 | 3008920 |
| **11** | 896400 | 976472 | 353119 | 1388023 | 1281143 | 1802051 | 600503 | 746589 | 684764 | 1967738 | 767496 | 805827 | 779130 | 992028 | 1681929 | 1855405 | 774488 | 1372400 | 1419668 | 2140293 | 1911739 | 2290655 |
| **12** | 776452 | 837553 | 309957 | 1155619 | 1073826 | 1442889 | 517346 | 642533 | 583503 | 1558222 | 647150 | 695652 | 658578 | 837413 | 1380861 | 1491219 | 676878 | 1162987 | 1206551 | 1655856 | 1514991 | 1735503 |
| **13** | 674528 | 719082 | 272826 | 961547 | 900729 | 1153170 | 444906 | 555089 | 499287 | 1232409 | 543863 | 600196 | 557318 | 708340 | 1128573 | 1193643 | 595318 | 983189 | 1016649 | 1284314 | 1201766 | 1325134 |
| **14** | 591611 | 624415 | 243169 | 804264 | 760629 | 928442 | 386577 | 485758 | 433556 | 981109 | 463462 | 524630 | 473064 | 604915 | 923518 | 959393 | 532318 | 834414 | 854281 | 1006832 | 961461 | 1026987 |
| **15** | 531882 | 555935 | 226516 | 691484 | 660264 | 767442 | 345412 | 437166 | 387259 | 801569 | 406568 | 473639 | 415398 | 534314 | 772330 | 790396 | 491310 | 718921 | 732122 | 813289 | 790179 | 822893 |
| **16** | 484794 | 502428 | 216795 | 601257 | 579791 | 645448 | 318852 | 400899 | 355454 | 665170 | 368721 | 435951 | 375178 | 482371 | 651764 | 659727 | 460358 | 622936 | 629309 | 670055 | 658998 | 674141 |
| **17** | 452352 | 464419 | 217526 | 534905 | 521144 | 557829 | 308788 | 381712 | 340590 | 568350 | 348229 | 415554 | 353956 | 449253 | 562713 | 565962 | 439161 | 549046 | 552045 | 570074 | 565224 | 571696 |
| **18** | 423241 | 430627 | 225290 | 475214 | 467885 | 485349 | 307146 | 367272 | 332532 | 490112 | 336275 | 398062 | 340765 | 420106 | 488174 | 489127 | 417102 | 482554 | 483737 | 490522 | 488816 | 491106 |
| **19** | 384826 | 388691 | 237470 | 412079 | 408970 | 415568 | 306217 | 349479 | 323388 | 417164 | 323616 | 371560 | 326559 | 383523 | 416675 | 416916 | 383541 | 414942 | 415353 | 417267 | 416777 | 417399 |
| **20** | 324928 | 326219 | 244869 | 334194 | 333318 | 334849 | 289410 | 310837 | 297762 | 335160 | 295240 | 320754 | 295742 | 325007 | 335096 | 335135 | 325406 | 334782 | 334856 | 335177 | 335099 | 335192 |
| **21** | 220783 | 220783 | 220783 | 220783 | 220783 | 220783 | 220783 | 220783 | 220783 | 220783 | 220783 | 220783 | 220783 | 220783 | 220783 | 220783 | 220783 | 220783 | 220783 | 220783 | 220783 | 220783 |
| **All$c_p$** | 17890748 | 20021041 | 7571412 | 29349817 | 26727453 | 40748712 | 11681897 | 15769632 | 15239104 | 48632726 | 16427214 | 16293389 | 16086056 | 20631317 | 33052987 | 36043172 | 17180120 | 26812116 | 24103152 | 59959116 | 50145409 | 115402630 |

***Table 6. Counts of unique regions forming the long-range contacts per cell line or tissue*** *(CT) for minimum contact gaps, (**A**) 2 and (**B**) 0.5 Mb. "All$c_p$" and "AllCT" considered all $c_p$ values, and cell lines and tissues, respectively.*

**(A)**

| | Cell line / Tissue (CT) | | | | | | | | | | | | | | | | | | | | |
|---|---|---|---|---|---|---|---|---|---|---|---|---|---|---|---|---|---|---|---|---|---|
| $c_p$ | **Co** | **Hi** | **Lu** | **LV** | **RV** | **Ao** | **PM** | **Pa** | **Sp** | **Li** | **SB** | **AG** | **Ov** | **Bl** | **MesC** | **MSC** | **NPC** | **TLC** | **ESC** | **FC** | **LC** | **AllCT** |
| **1** | 68697 | 69160 | 63650 | 69875 | 69714 | 70526 | 66040 | 68205 | 68508 | 70684 | 67806 | 67832 | 67590 | 69073 | 69997 | 70250 | 68188 | 69660 | 68455 | 70767 | 70713 | 70917 |
| **2** | 70264 | 70348 | 68637 | 70551 | 70506 | 70634 | 69575 | 70078 | 70172 | 70676 | 69990 | 70048 | 69991 | 70343 | 70577 | 70637 | 70107 | 70482 | 70304 | 70676 | 70680 | 70781 |
| **3** | 70292 | 70318 | 69650 | 70461 | 70436 | 70510 | 70029 | 70220 | 70265 | 70532 | 70200 | 70220 | 70202 | 70376 | 70461 | 70513 | 70285 | 70442 | 70404 | 70539 | 70529 | 70615 |
| **4** | 70200 | 70197 | 69711 | 70306 | 70306 | 70365 | 70005 | 70135 | 70195 | 70404 | 70047 | 70135 | 70065 | 70239 | 70329 | 70377 | 70170 | 70293 | 70282 | 70425 | 70392 | 70490 |
| **5** | 70012 | 70042 | 69604 | 70173 | 70174 | 70233 | 69844 | 69999 | 70029 | 70253 | 69914 | 69946 | 69921 | 70058 | 70201 | 70240 | 69968 | 70143 | 70118 | 70266 | 70262 | 70352 |
| **6** | 69835 | 69867 | 69413 | 70005 | 69994 | 70066 | 69627 | 69809 | 69890 | 70125 | 69713 | 69787 | 69707 | 69887 | 70072 | 70093 | 69827 | 70008 | 69956 | 70139 | 70124 | 70212 |
| **7** | 69673 | 69702 | 69124 | 69823 | 69808 | 69886 | 69379 | 69632 | 69677 | 69944 | 69465 | 69531 | 69452 | 69682 | 69875 | 69889 | 69564 | 69823 | 69778 | 69962 | 69949 | 70034 |
| **8** | 69418 | 69451 | 68671 | 69617 | 69570 | 69693 | 69057 | 69399 | 69434 | 69753 | 69173 | 69270 | 69148 | 69427 | 69683 | 69727 | 69326 | 69620 | 69578 | 69786 | 69768 | 69849 |
| **9** | 69163 | 69195 | 68195 | 69409 | 69330 | 69472 | 68650 | 69099 | 69175 | 69543 | 68821 | 68917 | 68764 | 69208 | 69453 | 69506 | 69049 | 69375 | 69324 | 69572 | 69576 | 69639 |
| **10** | 68821 | 68825 | 67512 | 69049 | 69022 | 69172 | 68188 | 68749 | 68780 | 69281 | 68375 | 68495 | 68238 | 68830 | 69198 | 69219 | 68584 | 69082 | 69042 | 69290 | 69304 | 69365 |
| **11** | 68346 | 68376 | 66774 | 68672 | 68601 | 68812 | 67547 | 68265 | 68250 | 68906 | 67781 | 67972 | 67583 | 68298 | 68833 | 68830 | 67982 | 68733 | 68657 | 68938 | 68926 | 69011 |
| **12** | 67669 | 67778 | 65591 | 68106 | 67990 | 68280 | 66721 | 67614 | 67595 | 68414 | 67034 | 67304 | 66784 | 67718 | 68264 | 68313 | 67300 | 68173 | 68072 | 68432 | 68435 | 68507 |
| **13** | 66827 | 66959 | 64338 | 67334 | 67244 | 67586 | 65572 | 66692 | 66654 | 67706 | 65986 | 66411 | 65868 | 66811 | 67561 | 67596 | 66336 | 67446 | 67305 | 67747 | 67737 | 67816 |
| **14** | 65616 | 65743 | 62365 | 66145 | 66038 | 66458 | 64053 | 65369 | 65175 | 66596 | 64618 | 65133 | 64347 | 65534 | 66427 | 66484 | 64906 | 66272 | 66165 | 66617 | 66615 | 66694 |
| **15** | 63918 | 64019 | 60054 | 64527 | 64397 | 64841 | 62015 | 63523 | 63216 | 65006 | 62695 | 63300 | 62419 | 63767 | 64864 | 64910 | 63075 | 64684 | 64548 | 65068 | 65048 | 65121 |
| **16** | 61144 | 61254 | 56770 | 61951 | 61794 | 62285 | 58976 | 60624 | 60256 | 62462 | 59829 | 60523 | 59576 | 61070 | 62299 | 62377 | 60287 | 62125 | 62001 | 62519 | 62466 | 62571 |
| **17** | 57082 | 57322 | 52103 | 57993 | 57812 | 58370 | 54763 | 56478 | 55912 | 58521 | 55509 | 56479 | 55335 | 57067 | 58367 | 58473 | 56263 | 58237 | 58135 | 58559 | 58526 | 58609 |
| **18** | 50850 | 51004 | 45760 | 51696 | 51557 | 52021 | 48563 | 50182 | 49451 | 52170 | 49165 | 50260 | 49224 | 50797 | 52043 | 52106 | 50020 | 51893 | 51833 | 52187 | 52142 | 52223 |
| **19** | 41414 | 41502 | 37079 | 42087 | 41986 | 42296 | 39560 | 40727 | 40042 | 42401 | 40199 | 40936 | 40097 | 41379 | 42345 | 42357 | 40877 | 42229 | 42245 | 42410 | 42373 | 42423 |
| **20** | 28511 | 28551 | 26120 | 28866 | 28793 | 28943 | 27532 | 28131 | 27743 | 28973 | 27787 | 28326 | 27914 | 28506 | 28958 | 28972 | 28246 | 28914 | 28918 | 28978 | 28960 | 28980 |
| **21** | 13049 | 13049 | 13049 | 13049 | 13049 | 13049 | 13049 | 13049 | 13049 | 13049 | 13049 | 13049 | 13049 | 13049 | 13049 | 13049 | 13049 | 13049 | 13049 | 13049 | 13049 | 13049 |
| **All$c_p$** | 70751 | 70761 | 70655 | 70805 | 70794 | 70832 | 70708 | 70729 | 70770 | 70854 | 70725 | 70736 | 70733 | 70776 | 70787 | 70807 | 70756 | 70774 | 70756 | 70832 | 70824 | 70917 |

**(B)**

| | Cell line / Tissue (CT) | | | | | | | | | | | | | | | | | | | | |
|---|---|---|---|---|---|---|---|---|---|---|---|---|---|---|---|---|---|---|---|---|---|
| $c_p$ | **Co** | **Hi** | **Lu** | **LV** | **RV** | **Ao** | **PM** | **Pa** | **Sp** | **Li** | **SB** | **AG** | **Ov** | **Bl** | **MesC** | **MSC** | **NPC** | **TLC** | **ESC** | **FC** | **LC** | **AllCT** |
| **1** | 68704 | 69170 | 63662 | 69885 | 69726 | 70531 | 66050 | 68213 | 68516 | 70687 | 67813 | 67851 | 67599 | 69086 | 70004 | 70256 | 68207 | 69680 | 68476 | 70772 | 70718 | 70920 |
| **2** | 70290 | 70370 | 68669 | 70580 | 70524 | 70657 | 69605 | 70100 | 70201 | 70707 | 70012 | 70078 | 70007 | 70364 | 70623 | 70678 | 70135 | 70524 | 70369 | 70707 | 70705 | 70805 |
| **3** | 70354 | 70381 | 69693 | 70522 | 70500 | 70566 | 70077 | 70262 | 70329 | 70616 | 70237 | 70284 | 70238 | 70435 | 70566 | 70592 | 70352 | 70515 | 70510 | 70605 | 70604 | 70693 |
| **4** | 70276 | 70288 | 69767 | 70434 | 70417 | 70465 | 70075 | 70211 | 70285 | 70518 | 70146 | 70234 | 70122 | 70332 | 70482 | 70499 | 70279 | 70416 | 70429 | 70519 | 70509 | 70594 |
| **5** | 70166 | 70175 | 69731 | 70355 | 70337 | 70373 | 69982 | 70136 | 70184 | 70427 | 70033 | 70100 | 70036 | 70201 | 70401 | 70420 | 70162 | 70340 | 70335 | 70435 | 70430 | 70504 |
| **6** | 70044 | 70061 | 69607 | 70258 | 70237 | 70283 | 69817 | 70003 | 70108 | 70361 | 69893 | 69997 | 69868 | 70086 | 70317 | 70343 | 70078 | 70285 | 70252 | 70360 | 70361 | 70432 |
| **7** | 69952 | 69975 | 69415 | 70179 | 70124 | 70192 | 69642 | 69907 | 70026 | 70276 | 69742 | 69859 | 69686 | 69987 | 70243 | 70239 | 69959 | 70178 | 70163 | 70274 | 70264 | 70320 |
| **8** | 69851 | 69878 | 69128 | 70120 | 70047 | 70117 | 69467 | 69795 | 69916 | 70195 | 69598 | 69734 | 69509 | 69859 | 70177 | 70171 | 69854 | 70105 | 70106 | 70202 | 70214 | 70260 |
| **9** | 69749 | 69768 | 68908 | 70041 | 69963 | 70025 | 69272 | 69707 | 69788 | 70131 | 69404 | 69560 | 69322 | 69785 | 70104 | 70118 | 69764 | 70028 | 70002 | 70129 | 70134 | 70182 |
| **10** | 69629 | 69647 | 68585 | 69903 | 69864 | 69932 | 69077 | 69581 | 69681 | 70014 | 69195 | 69443 | 69096 | 69651 | 70008 | 70003 | 69600 | 69932 | 69921 | 70023 | 70030 | 70056 |
| **11** | 69505 | 69530 | 68327 | 69810 | 69768 | 69832 | 68846 | 69461 | 69563 | 69928 | 69057 | 69337 | 68847 | 69506 | 69916 | 69901 | 69478 | 69849 | 69835 | 69941 | 69929 | 69962 |
| **12** | 69349 | 69437 | 67918 | 69670 | 69623 | 69699 | 68590 | 69321 | 69414 | 69784 | 68872 | 69161 | 68513 | 69347 | 69778 | 69779 | 69364 | 69721 | 69700 | 69798 | 69800 | 69820 |
| **13** | 69241 | 69244 | 67659 | 69571 | 69483 | 69575 | 68428 | 69198 | 69252 | 69664 | 68646 | 69039 | 68366 | 69242 | 69642 | 69632 | 69198 | 69607 | 69586 | 69665 | 69668 | 69682 |
| **14** | 69040 | 69103 | 67334 | 69353 | 69306 | 69367 | 68149 | 68982 | 69032 | 69453 | 68430 | 68825 | 68154 | 69024 | 69434 | 69430 | 69004 | 69400 | 69371 | 69465 | 69457 | 69471 |
| **15** | 68877 | 68884 | 67168 | 69135 | 69099 | 69163 | 67910 | 68767 | 68851 | 69243 | 68154 | 68613 | 67897 | 68804 | 69235 | 69223 | 68817 | 69205 | 69159 | 69248 | 69239 | 69254 |
| **16** | 68535 | 68568 | 66931 | 68830 | 68773 | 68858 | 67562 | 68416 | 68513 | 68914 | 67828 | 68336 | 67545 | 68505 | 68911 | 68907 | 68496 | 68882 | 68856 | 68917 | 68912 | 68926 |
| **17** | 68079 | 68147 | 66688 | 68385 | 68341 | 68396 | 67233 | 68015 | 68069 | 68455 | 67488 | 67943 | 67111 | 68060 | 68447 | 68451 | 68054 | 68427 | 68409 | 68450 | 68451 | 68458 |
| **18** | 67365 | 67410 | 66015 | 67635 | 67587 | 67632 | 66494 | 67247 | 67325 | 67681 | 66710 | 67202 | 66458 | 67326 | 67677 | 67675 | 67351 | 67666 | 67655 | 67682 | 67686 | 67687 |
| **19** | 66185 | 66237 | 64941 | 66414 | 66368 | 66414 | 65449 | 66087 | 66102 | 66445 | 65675 | 66043 | 65382 | 66145 | 66445 | 66443 | 66173 | 66427 | 66428 | 66447 | 66445 | 66448 |
| **20** | 63740 | 63787 | 62778 | 63888 | 63877 | 63902 | 63222 | 63645 | 63655 | 63915 | 63349 | 63712 | 63191 | 63726 | 63914 | 63914 | 63741 | 63911 | 63904 | 63915 | 63913 | 63916 |
| **21** | 57348 | 57348 | 57348 | 57348 | 57348 | 57348 | 57348 | 57348 | 57348 | 57348 | 57348 | 57348 | 57348 | 57348 | 57348 | 57348 | 57348 | 57348 | 57348 | 57348 | 57348 | 57348 |
| **All$c_p$** | 70758 | 70768 | 70667 | 70810 | 70801 | 70835 | 70720 | 70735 | 70781 | 70856 | 70733 | 70750 | 70737 | 70782 | 70795 | 70809 | 70762 | 70784 | 70772 | 70836 | 70829 | 70920 |

***Table 7. List of features used in this study*** *obtained from (**A**) published studies and databases, and (**B**-**D**) ENCODE [105, 106]. Features with cell type "hg19" are cell-type-invariant ones. Features with the affix "shared" on their names were generated by extracting regions shared by all cell types, tissues or species with available data. Sources in (**A**) are **0)** this study, **1)** Sadaie et al. [117] and van Steensel and Belmont [118], **2)** Costantini et al. [13] and Jabbari and Bernardi [14], **3)** Liu et al. [119], **4)** Manville et al. [120], **5)** Nash et al. [121], **6)** NIH Roadmap Epigenomics Mapping Consortium [107], **7)** non-B DB [122], **8)** Rao et al. [12], **9)** Sahakyan et al. [123], **10)** Schmitt et al. [1], **11)** UCSC Table Browser [113], and **12)** Rao et al. [12] and Xiong et al. [11]. Refer to **Table 1** for the abbreviations of the cell lines and tissues.*

**(A)**

| | Cell type | Feature | Source | Cell type | Feature | Source |
|---|---|---|---|---|---|---|
| 1 | ESC;FC | LMNB1 | 1 | ESC;FC | H4K8ac_br | 6 |
| 2 | hg19 | LMNB1_shared | 0;1 | ESC;FC | H4K91ac_br | 6 |
| 3 | hg19 | H | 0;2 | ESC;FC | DNase.hotspot.all_na | 6 |
| 4 | hg19 | H1 | 0;2 | ESC;FC | DNase.hotspot.fdr0.01_na | 6 |
| 5 | hg19 | H12 | 0;2 | ESC;FC;LC | DNase.macs2_na | 6 |
| 6 | hg19 | H2 | 0;2 | ESC;FC;LC | H2A.Z_na | 6 |
| 7 | hg19 | H23 | 0;2 | ESC;FC | H2AK5ac_na | 6 |
| 8 | hg19 | H3 | 0;2 | FC | H2AK9ac_na | 6 |
| 9 | hg19 | L | 0;2 | ESC;FC | H2BK120ac_na | 6 |
| 10 | hg19 | L1 | 0;2 | ESC;FC | H2BK12ac_na | 6 |
| 11 | hg19 | L12 | 0;2 | ESC;FC | H2BK15ac_na | 6 |
| 12 | hg19 | L2 | 0;2 | ESC;FC | H2BK20ac_na | 6 |
| 13 | hg19 | L2H1 | 0;2 | ESC;FC | H2BK5ac_na | 6 |
| 14 | hg19 | LH | 0;2 | ESC;FC | H3K14ac_na | 6 |
| 15 | hg19 | forest | 3 | ESC;FC | H3K18ac_na | 6 |
| 16 | hg19 | prairie | 3 | ESC;FC | H3K23ac_na | 6 |
| 17 | hg19 | TOP2B | 4 | ESC | H3K23me2_na | 6 |
| 18 | hg19 | GRB_canFam3 | 5 | ESC;FC;LC | H3K27ac_na | 6 |
| 19 | hg19 | GRB_galGal4 | 5 | ESC;FC;LC | H3K27me3_na | 6 |
| 20 | hg19 | GRB_gorGor3 | 5 | ESC;FC;LC | H3K36me3_na | 6 |
| 21 | hg19 | GRB_lepOcu1 | 5 | ESC;FC | H3K4ac_na | 6 |
| 22 | hg19 | GRB_monDom5 | 5 | ESC;FC;LC | H3K4me1_na | 6 |
| 23 | hg19 | GRB_rheMac3 | 5 | ESC;FC;LC | H3K4me2_na | 6 |
| 24 | hg19 | huCNE_canFam3_100pc_50col | 5 | ESC;FC;LC | H3K4me3_na | 6 |
| 25 | hg19 | huCNE_canFam3_96pc_50col | 5 | ESC;FC | H3K56ac_na | 6 |
| 26 | hg19 | huCNE_galGal4_80pc_50col | 5 | ESC;FC | H3K79me1_na | 6 |
| 27 | hg19 | huCNE_galGal4_90pc_50col | 5 | ESC;FC;LC | H3K79me2_na | 6 |
| 28 | hg19 | huCNE_galGal4_98pc_50col | 5 | ESC;FC;LC | H3K9ac_na | 6 |
| 29 | hg19 | huCNE_gorGor3_100pc_400col | 5 | FC | H3K9me1_na | 6 |
| 30 | hg19 | huCNE_gorGor3_100pc_500col | 5 | ESC;FC;LC | H3K9me3_na | 6 |
| 31 | hg19 | huCNE_LepOcu1_70pc_30col | 5 | ESC;FC;LC | H4K20me1_na | 6 |
| 32 | hg19 | huCNE_LepOcu1_80pc_30col | 5 | ESC;FC | H4K5ac_na | 6 |
| 33 | hg19 | huCNE_LepOcu1_96.6pc_30col | 5 | ESC;FC | H4K8ac_na | 6 |
| 34 | hg19 | huCNE_monDom5_100pc_50col | 5 | ESC;FC | H4K91ac_na | 6 |
| 35 | hg19 | huCNE_monDom5_80pc_50col | 5 | hg19 | H3K9me3brna_shared | 0;6 |
| 36 | hg19 | huCNE_monDom5_96pc_50col | 5 | hg19 | H3K9me3br_shared | 0;6 |
| 37 | hg19 | huCNE_rheMac3_100pc_150col | 5 | hg19 | H3K9me3na_shared | 0;6 |
| 38 | hg19 | huCNE_rheMac3_99.3pc_150col | 5 | hg19 | A_Phased_Repeat | 7 |
| 39 | hg19 | GRB_shared | 0;5 | hg19 | Cruciform_Motif | 7 |
| 40 | hg19 | huCNE_shared | 0;5 | hg19 | Direct_Repeat | 7 |
| 41 | ESC;FC | DNase.hotspot_br | 6 | hg19 | G_Quadruplex_Motif | 7 |
| 42 | ESC;FC | DNase.hotspot.fdr0.01_br | 6 | hg19 | Inverted_Repeat | 7 |
| 43 | ESC;FC;LC | H2A.Z_br | 6 | hg19 | Mirror_Repeat | 7 |
| 44 | ESC;FC | H2AK5ac_br | 6 | hg19 | Short_Tandem_Repeat | 7 |
| 45 | FC | H2AK9ac_br | 6 | hg19 | Slipped_Motif | 7 |
| 46 | ESC;FC | H2BK120ac_br | 6 | hg19 | Triplex_Motif | 7 |
| 47 | ESC;FC | H2BK12ac_br | 6 | hg19 | Z_DNA_Motif | 7 |
| 48 | ESC;FC | H2BK15ac_br | 6 | LC | B4 | 8 |
| 49 | ESC;FC | H2BK20ac_br | 6 | hg19 | G-quadruplex_greq19 | 9 |
| 50 | ESC;FC | H2BK5ac_br | 6 | All 21 | A | 10 |
| 51 | ESC;FC | H3K14ac_br | 6 | All 21 | B | 10 |
| 52 | ESC;FC | H3K18ac_br | 6 | hg19 | A_shared | 0;10 |
| 53 | ESC;FC | H3K23ac_br | 6 | hg19 | B_shared | 0;10 |
| 54 | ESC | H3K23me2_br | 6 | hg19 | AAGCTTCC | 0 |
| 55 | ESC;FC;LC | H3K27ac_br | 6 | hg19 | AAGCTTCT | 0 |
| 56 | ESC;FC;LC | H3K27me3_br | 6 | hg19 | CAAGCTTT | 0 |
| 57 | ESC;FC;LC | H3K36me3_br | 6 | hg19 | TTAAGCTT | 0 |
| 58 | ESC;FC | H3K4ac_br | 6 | hg19 | centromere | 11 |
| 59 | ESC;FC;LC | H3K4me1_br | 6 | hg19 | CGI_m | 11 |
| 60 | ESC;FC;LC | H3K4me2_br | 6 | hg19 | CGI_um | 11 |
| 61 | ESC;FC;LC | H3K4me3_br | 6 | hg19 | genes | 11 |
| 62 | ESC;FC | H3K56ac_br | 6 | hg19 | genes_coding | 11 |
| 63 | ESC;FC | H3K79me1_br | 6 | hg19 | repeat | 11 |
| 64 | ESC;FC;LC | H3K79me2_br | 6 | hg19 | genes_LTr | 0;11 |
| 65 | ESC;FC;LC | H3K9ac_br | 6 | FC;hg19;LC | A1 | 12 |
| 66 | FC | H3K9me1_br | 6 | FC;hg19;LC | A2 | 12 |
| 67 | ESC;FC;LC | H3K9me3_br | 6 | FC;hg19;LC | B1 | 12 |
| 68 | ESC;FC;LC | H4K20me1_br | 6 | FC;hg19;LC | B2 | 12 |
| 69 | ESC;FC | H4K5ac_br | 6 | FC;hg19;LC | B3 | 12 |

(B)

| target | Experiment target | File accession | index | File format | Output type | Experiment accession | Assay | Biosample term name | Biosample type | Experiment date released | Project | Assembly | File Status |
|---|---|---|---|---|---|---|---|---|---|---|---|---|---|
| ELK1-human | ELK1-human | ENCFF757YVY | 5 | bed narrowPeak | optimal idr thresholded peaks | ENCSR664OKA | ChIP-seq | IMR-90 | cell line | 01/09/2016 | ENCODE | hg19 | released |
| FOS-human | FOS-human | ENCFF359IPV | 13 | bed narrowPeak | optimal idr thresholded peaks | ENCSR124AIG | ChIP-seq | IMR-90 | cell line | 19/12/2016 | ENCODE | hg19 | released |
| CEBPB-human | CEBPB-human | ENCFF002CVG | 23 | bed narrowPeak | optimal idr thresholded peaks | ENCSR000EFM | ChIP-seq | IMR-90 | cell line | 14/05/2012 | ENCODE | hg19 | archived |
| RAD21-human | RAD21-human | ENCFF002CVK | 36 | bed narrowPeak | optimal idr thresholded peaks | ENCSR000EFJ | ChIP-seq | IMR-90 | cell line | 14/05/2012 | ENCODE | hg19 | archived |
| MAFK-human | MAFK-human | ENCFF002CVI | 48 | bed narrowPeak | optimal idr thresholded peaks | ENCSR000EFH | ChIP-seq | IMR-90 | cell line | 14/05/2012 | ENCODE | hg19 | archived |
| CHD1-human | CHD1-human | ENCFF441IFB | 72 | bed narrowPeak | optimal idr thresholded peaks | ENCSR000EFC | ChIP-seq | IMR-90 | cell line | 20/08/2012 | ENCODE | hg19 | released |
| MAZ-human | MAZ-human | ENCFF851XSC | 85 | bed narrowPeak | optimal idr thresholded peaks | ENCSR000EFF | ChIP-seq | IMR-90 | cell line | 20/08/2012 | ENCODE | hg19 | released |
| USF2-human | USF2-human | ENCFF759MVB | 89 | bed narrowPeak | optimal idr thresholded peaks | ENCSR513UQG | ChIP-seq | IMR-90 | cell line | 15/06/2017 | ENCODE | hg19 | released |
| SMC3-human | SMC3-human | ENCFF116RLU | 101 | bed narrowPeak | optimal idr thresholded peaks | ENCSR000HPG | ChIP-seq | IMR-90 | cell line | 26/10/2016 | ENCODE | hg19 | released |
| CTCF-human | CTCF-human | ENCFF002CVH | 108 | bed narrowPeak | optimal idr thresholded peaks | ENCSR000EFI | ChIP-seq | IMR-90 | cell line | 14/05/2012 | ENCODE | hg19 | archived |
| BHLHE40-human | BHLHE40-human | ENCFF942FBW | 120 | bed narrowPeak | optimal idr thresholded peaks | ENCSR957KYB | ChIP-seq | IMR-90 | cell line | 01/09/2016 | ENCODE | hg19 | released |
| POLR2A-human | POLR2A-human | ENCFF002CVJ | 128 | bed narrowPeak | optimal idr thresholded peaks | ENCSR000EFK | ChIP-seq | IMR-90 | cell line | 14/05/2012 | ENCODE | hg19 | archived |
| NFE2L2-human | NFE2L2-human | ENCFF641RNZ | 133 | bed narrowPeak | optimal idr thresholded peaks | ENCSR197WGI | ChIP-seq | IMR-90 | cell line | 01/09/2016 | ENCODE | hg19 | released |

(C)

| target | Experiment target | File accession | index | File format | Output type | Experiment accession | Assay | Biosample term name | Biosample type | Experiment date released | Project | Assembly | File Status |
|---|---|---|---|---|---|---|---|---|---|---|---|---|---|
| BCL11A-human | BCL11A-human | ENCFF002CIT | 190 | bed narrowPeak | optimal idr thresholded peaks | ENCSR000BIP | ChIP-seq | H1 | cell line | 18/07/2011 | ENCODE | hg19 | archived |
| USF1-human | USF1-human | ENCFF358BEF | 14 | bed narrowPeak | optimal idr thresholded peaks | ENCSR000BIU | ChIP-seq | H1 | cell line | 18/07/2011 | ENCODE | hg19 | released |
| MYC-human | MYC-human | ENCFF002CQV | 20 | bed narrowPeak | optimal idr thresholded peaks | ENCSR000EBY | ChIP-seq | H1 | cell line | 14/05/2012 | ENCODE | hg19 | archived |
| ASH2L-human | ASH2L-human | ENCFF254HZM | 32 | bed narrowPeak | pseudoreplicated idr thresholded peaks | ENCSR850KIP | ChIP-seq | H1 | cell line | 28/11/2017 | ENCODE | hg19 | released |
| KDM1A-human | KDM1A-human | ENCFF768BPC | 36 | bed narrowPeak | optimal idr thresholded peaks | ENCSR418RRF | ChIP-seq | H1 | cell line | 06/12/2016 | ENCODE | hg19 | released |
| CBX8-human | CBX8-human | ENCFF485WCF | 49 | bed narrowPeak | optimal idr thresholded peaks | ENCSR771SNW | ChIP-seq | H1 | cell line | 06/12/2016 | ENCODE | hg19 | released |
| SIN3A-human | SIN3A-human | ENCFF184HWT | 606 | bed narrowPeak | optimal idr thresholded peaks | ENCSR000EBO | ChIP-seq | H1 | cell line | 14/05/2012 | ENCODE | hg19 | released |
| SIX5-human | SIX5-human | ENCFF002CJJ | 65 | bed narrowPeak | optimal idr thresholded peaks | ENCSR000BIQ | ChIP-seq | H1 | cell line | 18/07/2011 | ENCODE | hg19 | archived |
| GABPA-human | GABPA-human | ENCFF894MNP | 86 | bed narrowPeak | optimal idr thresholded peaks | ENCSR000BIW | ChIP-seq | H1 | cell line | 18/07/2011 | ENCODE | hg19 | released |
| TCF12-human | TCF12-human | ENCFF274AQE | 99 | bed narrowPeak | optimal idr thresholded peaks | ENCSR000BIT | ChIP-seq | H1 | cell line | 18/07/2011 | ENCODE | hg19 | released |
| ZNF143-human | ZNF143-human | ENCFF002CRJ | 103 | bed narrowPeak | optimal idr thresholded peaks | ENCSR000EBW | ChIP-seq | H1 | cell line | 14/05/2012 | ENCODE | hg19 | archived |
| CHD1-human | CHD1-human | ENCFF002CDR | 115 | bed narrowPeak | optimal idr thresholded peaks | ENCSR000AQK | ChIP-seq | H1 | cell line | 06/03/2012 | ENCODE | hg19 | archived |
| SAP30-human | SAP30-human | ENCFF361QZZ | 136 | bed narrowPeak | optimal idr thresholded peaks | ENCSR000ATR | ChIP-seq | H1 | cell line | 06/08/2012 | ENCODE | hg19 | released |
| RBBP5-human | RBBP5-human | ENCFF135DOZ | 144 | bed narrowPeak | optimal idr thresholded peaks | ENCSR000AQC | ChIP-seq | H1 | cell line | 06/03/2012 | ENCODE | hg19 | released |
| CTCF-human | CTCF-human | ENCFF002CDS | 150 | bed narrowPeak | optimal idr thresholded peaks | ENCSR000AMF | ChIP-seq | H1 | cell line | 10/02/2011 | ENCODE | hg19 | archived |
| MXI1-human | MXI1-human | ENCFF002CRB | 156 | bed narrowPeak | optimal idr thresholded peaks | ENCSR000EBR | ChIP-seq | H1 | cell line | 14/05/2012 | ENCODE | hg19 | archived |
| HDAC6-human | HDAC6-human | ENCFF012LET | 158 | bed narrowPeak | pseudoreplicated idr thresholded peaks | ENCSR000ATQ | ChIP-seq | H1 | cell line | 06/08/2012 | ENCODE | hg19 | released |
| REST-human | REST-human | ENCFF453HPD | 171 | bed narrowPeak | optimal idr thresholded peaks | ENCSR000BHM | ChIP-seq | H1 | cell line | 18/07/2011 | ENCODE | hg19 | released |
| ATF2-human | ATF2-human | ENCFF002CIR | 179 | bed narrowPeak | optimal idr thresholded peaks | ENCSR000BQU | ChIP-seq | H1 | cell line | 29/02/2012 | ENCODE | hg19 | archived |
| PHF8-human | PHF8-human | ENCFF687NII | 184 | bed narrowPeak | optimal idr thresholded peaks | ENCSR000ATK | ChIP-seq | H1 | cell line | 06/08/2012 | ENCODE | hg19 | released |
| YY1-human | YY1-human | ENCFF686VJY | 206 | bed narrowPeak | optimal idr thresholded peaks | ENCSR000BKD | ChIP-seq | H1 | cell line | 18/07/2011 | ENCODE | hg19 | released |
| CHD7-human | CHD7-human | ENCFF628RLE | 213 | bed narrowPeak | optimal idr thresholded peaks | ENCSR000AVA | ChIP-seq | H1 | cell line | 06/08/2012 | ENCODE | hg19 | released |
| EP300-human | EP300-human | ENCFF843SYD | 385 | bed narrowPeak | optimal idr thresholded peaks | ENCSR000BKK | ChIP-seq | H1 | cell line | 18/07/2011 | ENCODE | hg19 | released |
| BRCA1-human | BRCA1-human | ENCFF002CQQ | 228 | bed narrowPeak | optimal idr thresholded peaks | ENCSR000EBX | ChIP-seq | H1 | cell line | 14/05/2012 | ENCODE | hg19 | archived |
| TAF7-human | TAF7-human | ENCFF963TZI | 249 | bed narrowPeak | optimal idr thresholded peaks | ENCSR000BLU | ChIP-seq | H1 | cell line | 18/07/2011 | ENCODE | hg19 | released |
| SP4-human | SP4-human | ENCFF002CJM | 254 | bed narrowPeak | optimal idr thresholded peaks | ENCSR000BQV | ChIP-seq | H1 | cell line | 29/02/2012 | ENCODE | hg19 | archived |
| TEAD4-human | TEAD4-human | ENCFF002CJR | 257 | bed narrowPeak | optimal idr thresholded peaks | ENCSR000BRY | ChIP-seq | H1 | cell line | 10/09/2012 | ENCODE | hg19 | released |
| ATF3-human | ATF3-human | ENCFF002CIS | 272 | bed narrowPeak | optimal idr thresholded peaks | ENCSR000BKC | ChIP-seq | H1 | cell line | 18/07/2011 | ENCODE | hg19 | archived |
| FOSL1-human | FOSL1-human | ENCFF002CIW | 285 | bed narrowPeak | optimal idr thresholded peaks | ENCSR000BNS | ChIP-seq | H1 | cell line | 18/07/2011 | ENCODE | hg19 | archived |
| HDAC2-human | HDAC2-human | ENCFF468CFJ | 312 | bed narrowPeak | optimal idr thresholded peaks | ENCSR000AVB | ChIP-seq | H1 | cell line | 06/08/2012 | ENCODE | hg19 | released |
| CTBP2-human | CTBP2-human | ENCFF222TBM | 325 | bed narrowPeak | optimal idr thresholded peaks | ENCSR000EUO | ChIP-seq | H1 | cell line | 29/10/2011 | ENCODE | hg19 | released |
| JUND-human | JUND-human | ENCFF073PCP | 343 | bed narrowPeak | optimal idr thresholded peaks | ENCSR000BKP | ChIP-seq | H1 | cell line | 18/07/2011 | ENCODE | hg19 | released |
| RAD21-human | RAD21-human | ENCFF002CJG | 514 | bed narrowPeak | optimal idr thresholded peaks | ENCSR000BLD | ChIP-seq | H1 | cell line | 18/07/2011 | ENCODE | hg19 | archived |
| KDM4A-human | KDM4A-human | ENCFF838KYH | 359 | bed narrowPeak | pseudoreplicated idr thresholded peaks | ENCSR000AVC | ChIP-seq | H1 | cell line | 06/08/2012 | ENCODE | hg19 | released |
| BACH1-human | BACH1-human | ENCFF002CQP | 365 | bed narrowPeak | optimal idr thresholded peaks | ENCSR000EBQ | ChIP-seq | H1 | cell line | 14/05/2012 | ENCODE | hg19 | archived |
| GTF2F1-human | GTF2F1-human | ENCFF002CQX | 390 | bed narrowPeak | optimal idr thresholded peaks | ENCSR000EBP | ChIP-seq | H1 | cell line | 14/05/2012 | ENCODE | hg19 | archived |
| POU5F1-human | POU5F1-human | ENCFF002CJF | 403 | bed narrowPeak | optimal idr thresholded peaks | ENCSR000BMU | ChIP-seq | H1 | cell line | 18/07/2011 | ENCODE | hg19 | archived |
| CBX5-human | CBX5-human | ENCFF936JMD | 415 | bed narrowPeak | optimal idr thresholded peaks | ENCSR814KIO | ChIP-seq | H1 | cell line | 06/12/2016 | ENCODE | hg19 | released |
| CHD2-human | CHD2-human | ENCFF002CQT | 419 | bed narrowPeak | optimal idr thresholded peaks | ENCSR000EBT | ChIP-seq | H1 | cell line | 14/05/2012 | ENCODE | hg19 | archived |
| RXRA-human | RXRA-human | ENCFF369JAI | 442 | bed narrowPeak | optimal idr thresholded peaks | ENCSR000BJW | ChIP-seq | H1 | cell line | 18/07/2011 | ENCODE | hg19 | released |
| JUN-human | JUN-human | ENCFF002CQU | 444 | bed narrowPeak | optimal idr thresholded peaks | ENCSR000ECA | ChIP-seq | H1 | cell line | 29/10/2011 | ENCODE | hg19 | archived |
| EZH2-human | EZH2-human | ENCFF002CDT | 456 | bed narrowPeak | optimal idr thresholded peaks | ENCSR000ASY | ChIP-seq | H1 | cell line | 06/08/2012 | ENCODE | hg19 | archived |
| TAF1-human | TAF1-human | ENCFF202SNU | 468 | bed narrowPeak | optimal idr thresholded peaks | ENCSR000BHO | ChIP-seq | H1 | cell line | 18/07/2011 | ENCODE | hg19 | released |
| SUZ12-human | SUZ12-human | ENCFF336HCY | 477 | bed narrowPeak | optimal idr thresholded peaks | ENCSR000ATS | ChIP-seq | H1 | cell line | 06/08/2012 | ENCODE | hg19 | released |
| SIRT6-human | SIRT6-human | ENCFF719RAM | 487 | bed narrowPeak | pseudoreplicated idr thresholded peaks | ENCSR000AUS | ChIP-seq | H1 | cell line | 06/08/2012 | ENCODE | hg19 | released |
| RFX5-human | RFX5-human | ENCFF002CRE | 489 | bed narrowPeak | optimal idr thresholded peaks | ENCSR000ECF | ChIP-seq | H1 | cell line | 29/10/2011 | ENCODE | hg19 | archived |
| TBP-human | TBP-human | ENCFF002CRH | 524 | bed narrowPeak | optimal idr thresholded peaks | ENCSR000ECB | ChIP-seq | H1 | cell line | 29/10/2011 | ENCODE | hg19 | archived |
| NRF1-human | NRF1-human | ENCFF002CRC | 536 | bed narrowPeak | optimal idr thresholded peaks | ENCSR000ECC | ChIP-seq | H1 | cell line | 29/10/2011 | ENCODE | hg19 | archived |
| NANOG-human | NANOG-human | ENCFF379EPK | 559 | bed narrowPeak | optimal idr thresholded peaks | ENCSR000BMT | ChIP-seq | H1 | cell line | 18/07/2011 | ENCODE | hg19 | released |
| MAFK-human | MAFK-human | ENCFF002CQZ | 561 | bed narrowPeak | optimal idr thresholded peaks | ENCSR000EBS | ChIP-seq | H1 | cell line | 14/05/2012 | ENCODE | hg19 | archived |
| SRF-human | SRF-human | ENCFF002CJN | 574 | bed narrowPeak | optimal idr thresholded peaks | ENCSR000BIV | ChIP-seq | H1 | cell line | 18/07/2011 | ENCODE | hg19 | archived |
| SP1-human | SP1-human | ENCFF317TBZ | 596 | bed narrowPeak | optimal idr thresholded peaks | ENCSR000BIR | ChIP-seq | H1 | cell line | 18/07/2011 | ENCODE | hg19 | released |
| USF2-human | USF2-human | ENCFF002CRI | 611 | bed narrowPeak | optimal idr thresholded peaks | ENCSR000ECD | ChIP-seq | H1 | cell line | 29/10/2011 | ENCODE | hg19 | archived |
| EGR1-human | EGR1-human | ENCFF220IXP | 647 | bed narrowPeak | optimal idr thresholded peaks | ENCSR000BJA | ChIP-seq | H1 | cell line | 18/07/2011 | ENCODE | hg19 | released |
| KDM5A-human | KDM5A-human | ENCFF583YFS | 662 | bed narrowPeak | optimal idr thresholded peaks | ENCSR160ZLP | ChIP-seq | H1 | cell line | 06/03/2012 | ENCODE | hg19 | released |
| CEBPB-human | CEBPB-human | ENCFF002CQR | 671 | bed narrowPeak | optimal idr thresholded peaks | ENCSR000EBV | ChIP-seq | H1 | cell line | 14/05/2012 | ENCODE | hg19 | archived |
| SP2-human | SP2-human | ENCFF002CJL | 674 | bed narrowPeak | optimal idr thresholded peaks | ENCSR000BQG | ChIP-seq | H1 | cell line | 29/02/2012 | ENCODE | hg19 | archived |
| POLR2AphosphoS5-human | POLR2AphosphoS5-human | ENCFF450NDU | 682 | bed narrowPeak | optimal idr thresholded peaks | ENCSR000BIC | ChIP-seq | H1 | cell line | 18/07/2011 | ENCODE | hg19 | released |
| POLR2A-human | POLR2A-human | ENCFF468LYO | 698 | bed narrowPeak | optimal idr thresholded peaks | ENCSR000BHN | ChIP-seq | H1 | cell line | 18/07/2011 | ENCODE | hg19 | released |

(D)

| target | Experiment target | File accession | index | File format | Output type | Experiment accession | Assay | Biosample term name | Biosample type | Experiment date released | Project | Assembly | File Status |
|---|---|---|---|---|---|---|---|---|---|---|---|---|---|
| CHD4-human | CHD4-human | ENCFF499ZPP | 3 | bed narrowPeak | optimal idr thresholded peaks | ENCSR751CJG | ChIP-seq | GM12878 | cell line | 23/11/2016 | ENCODE | hg19 | released |
| RBBP5-human | RBBP5-human | ENCFF814CZI | 17 | bed narrowPeak | optimal idr thresholded peaks | ENCSR330EXS | ChIP-seq | GM12878 | cell line | 23/11/2016 | ENCODE | hg19 | released |
| CHD1-human | CHD1-human | ENCFF002CON | 22 | bed narrowPeak | optimal idr thresholded peaks | ENCSR000DZE | ChIP-seq | GM12878 | cell line | 14/05/2012 | ENCODE | hg19 | archived |
| ASH2L-human | ASH2L-human | ENCFF917CXD | 41 | bed narrowPeak | optimal idr thresholded peaks | ENCSR849WCQ | ChIP-seq | GM12878 | cell line | 08/01/2017 | ENCODE | hg19 | released |
| PKNOX1-human | PKNOX1-human | ENCFF020GMZ | 55 | bed narrowPeak | optimal idr thresholded peaks | ENCSR711XNY | ChIP-seq | GM12878 | cell line | 17/11/2016 | ENCODE | hg19 | released |
| PML-human | PML-human | ENCFF002CHM | 61 | bed narrowPeak | optimal idr thresholded peaks | ENCSR000BQM | ChIP-seq | GM12878 | cell line | 29/02/2012 | ENCODE | hg19 | archived |
| SIN3A-human | SIN3A-human | ENCFF778DJD | 68 | bed narrowPeak | optimal idr thresholded peaks | ENCSR000DYX | ChIP-seq | GM12878 | cell line | 14/05/2012 | ENCODE | hg19 | released |
| POU2F2-human | POU2F2-human | ENCFF002CHP | 77 | bed narrowPeak | optimal idr thresholded peaks | ENCSR000BGP | ChIP-seq | GM12878 | cell line | 18/07/2011 | ENCODE | hg19 | archived |
| BCL11A-human | BCL11A-human | ENCFF002CGR | 80 | bed narrowPeak | optimal idr thresholded peaks | ENCSR000BHA | ChIP-seq | GM12878 | cell line | 18/07/2011 | ENCODE | hg19 | archived |
| RXRA-human | RXRA-human | ENCFF461FGJ | 101 | bed narrowPeak | optimal idr thresholded peaks | ENCSR000BJD | ChIP-seq | GM12878 | cell line | 18/07/2011 | ENCODE | hg19 | released |
| PAX5-human | PAX5-human | ENCFF002CHJ | 106 | bed narrowPeak | optimal idr thresholded peaks | ENCSR000BHD | ChIP-seq | GM12878 | cell line | 18/07/2011 | ENCODE | hg19 | archived |
| BRCA1-human | BRCA1-human | ENCFF082JDH | 123 | bed narrowPeak | optimal idr thresholded peaks | ENCSR000DZS | ChIP-seq | GM12878 | cell line | 29/10/2011 | ENCODE | hg19 | released |
| MAZ-human | MAZ-human | ENCFF002COX | 130 | bed narrowPeak | optimal idr thresholded peaks | ENCSR000DZA | ChIP-seq | GM12878 | cell line | 14/05/2012 | ENCODE | hg19 | archived |
| YY1-human | YY1-human | ENCFF002CIC | 143 | bed narrowPeak | optimal idr thresholded peaks | ENCSR000BNP | ChIP-seq | GM12878 | cell line | 18/07/2011 | ENCODE | hg19 | archived |
| KDM1A-human | KDM1A-human | ENCFF996RRT | 159 | bed narrowPeak | optimal idr thresholded peaks | ENCSR391IWM | ChIP-seq | GM12878 | cell line | 24/11/2016 | ENCODE | hg19 | released |
| JUNB-human | JUNB-human | ENCFF939TZS | 167 | bed narrowPeak | optimal idr thresholded peaks | ENCSR897MMC | ChIP-seq | GM12878 | cell line | 13/06/2016 | ENCODE | hg19 | released |
| IKZF1-human | IKZF1-human | ENCFF229YSW | 950 | bed narrowPeak | optimal idr thresholded peaks | ENCSR874AFU | ChIP-seq | GM12878 | cell line | 20/09/2016 | ENCODE | hg19 | released |
| TCF3-human | TCF3-human | ENCFF002CIA | 188 | bed narrowPeak | optimal idr thresholded peaks | ENCSR000BQT | ChIP-seq | GM12878 | cell line | 29/02/2012 | ENCODE | hg19 | archived |
| BHLHE40-human | BHLHE40-human | ENCFF095GMM | 1016 | bed narrowPeak | optimal idr thresholded peaks | ENCSR987MTA | ChIP-seq | GM12878 | cell line | 31/01/2017 | ENCODE | hg19 | released |
| NR2C2-human | NR2C2-human | ENCFF002CPS | 200 | bed narrowPeak | optimal idr thresholded peaks | ENCSR000EUL | ChIP-seq | GM12878 | cell line | 29/10/2011 | ENCODE | hg19 | archived |
| JUND-human | JUND-human | ENCFF556PIY | 446 | bed narrowPeak | optimal idr thresholded peaks | ENCSR000DYS | ChIP-seq | GM12878 | cell line | 20/08/2012 | ENCODE | hg19 | released |
| NFYA-human | NFYA-human | ENCFF002CPB | 214 | bed narrowPeak | optimal idr thresholded peaks | ENCSR000DNN | ChIP-seq | GM12878 | cell line | 29/10/2011 | ENCODE | hg19 | archived |
| SPI1-human | SPI1-human | ENCFF002CHQ | 225 | bed narrowPeak | optimal idr thresholded peaks | ENCSR000BGQ | ChIP-seq | GM12878 | cell line | 18/07/2011 | ENCODE | hg19 | archived |
| ZBTB33-human | ZBTB33-human | ENCFF539CKD | 245 | bed narrowPeak | optimal idr thresholded peaks | ENCSR542FLV | ChIP-seq | GM12878 | cell line | 13/11/2017 | ENCODE | hg19 | released |
| ZBED1-human | ZBED1-human | ENCFF123WEZ | 255 | bed narrowPeak | optimal idr thresholded peaks | ENCSR207PFI | ChIP-seq | GM12878 | cell line | 05/02/2016 | ENCODE | hg19 | released |
| TBP-human | TBP-human | ENCFF002CPR | 257 | bed narrowPeak | optimal idr thresholded peaks | ENCSR000DZZ | ChIP-seq | GM12878 | cell line | 29/10/2011 | ENCODE | hg19 | archived |
| EGR1-human | EGR1-human | ENCFF002CGW | 269 | bed narrowPeak | optimal idr thresholded peaks | ENCSR000BRG | ChIP-seq | GM12878 | cell line | 29/02/2012 | ENCODE | hg19 | archived |
| ARNT-human | ARNT-human | ENCFF166OZW | 283 | bed narrowPeak | optimal idr thresholded peaks | ENCSR590KEQ | ChIP-seq | GM12878 | cell line | 14/11/2017 | ENCODE | hg19 | released |
| STAT1-human | STAT1-human | ENCFF817QHW | 1539 | bed narrowPeak | optimal idr thresholded peaks | ENCSR332EYT | ChIP-seq | GM12878 | cell line | 07/07/2014 | ENCODE | hg19 | released |
| MAFK-human | MAFK-human | ENCFF689AXJ | 289 | bed narrowPeak | optimal idr thresholded peaks | ENCSR000DYV | ChIP-seq | GM12878 | cell line | 20/08/2012 | ENCODE | hg19 | released |
| CTCF-human | CTCF-human | ENCFF002COQ | 1139 | bed narrowPeak | optimal idr thresholded peaks | ENCSR000DZN | ChIP-seq | GM12878 | cell line | 29/10/2011 | ENCODE | hg19 | archived |
| STAT5A-human | STAT5A-human | ENCFF492YMU | 316 | bed narrowPeak | optimal idr thresholded peaks | ENCSR000BQZ | ChIP-seq | GM12878 | cell line | 29/02/2012 | ENCODE | hg19 | released |
| RCOR1-human | RCOR1-human | ENCFF898MUV | 325 | bed narrowPeak | optimal idr thresholded peaks | ENCSR000DZC | ChIP-seq | GM12878 | cell line | 14/05/2012 | ENCODE | hg19 | released |
| MEF2C-human | MEF2C-human | ENCFF261ZWR | 346 | bed narrowPeak | optimal idr thresholded peaks | ENCSR000BNG | ChIP-seq | GM12878 | cell line | 18/07/2011 | ENCODE | hg19 | released |
| RAD21-human | RAD21-human | ENCFF002CHR | 1593 | bed narrowPeak | optimal idr thresholded peaks | ENCSR000BMY | ChIP-seq | GM12878 | cell line | 18/07/2011 | ENCODE | hg19 | archived |
| RAD51-human | RAD51-human | ENCFF445UJF | 355 | bed narrowPeak | optimal idr thresholded peaks | ENCSR482TWQ | ChIP-seq | GM12878 | cell line | 17/11/2016 | ENCODE | hg19 | released |
| HDGF-human | HDGF-human | ENCFF935HIG | 362 | bed narrowPeak | optimal idr thresholded peaks | ENCSR145XQO | ChIP-seq | GM12878 | cell line | 24/08/2016 | ENCODE | hg19 | released |
| NFATC3-human | NFATC3-human | ENCFF704ZYN | 375 | bed narrowPeak | optimal idr thresholded peaks | ENCSR437GBJ | ChIP-seq | GM12878 | cell line | 26/01/2018 | ENCODE | hg19 | released |
| TARDBP-human | TARDBP-human | ENCFF733UTZ | 388 | bed narrowPeak | optimal idr thresholded peaks | ENCSR412QBS | ChIP-seq | GM12878 | cell line | 22/04/2016 | ENCODE | hg19 | released |
| CEBPB-human | CEBPB-human | ENCFF002CGU | 391 | bed narrowPeak | optimal idr thresholded peaks | ENCSR000BRX | ChIP-seq | GM12878 | cell line | 10/09/2012 | ENCODE | hg19 | archived |
| HDAC2-human | HDAC2-human | ENCFF107UHX | 392 | bed narrowPeak | optimal idr thresholded peaks | ENCSR330OEO | ChIP-seq | GM12878 | cell line | 12/08/2017 | ENCODE | hg19 | released |
| SP1-human | SP1-human | ENCFF002CHV | 404 | bed narrowPeak | optimal idr thresholded peaks | ENCSR000BHK | ChIP-seq | GM12878 | cell line | 18/07/2011 | ENCODE | hg19 | released |
| BCLAF1-human | BCLAF1-human | ENCFF563QBC | 430 | bed narrowPeak | optimal idr thresholded peaks | ENCSR000BJZ | ChIP-seq | GM12878 | cell line | 18/07/2011 | ENCODE | hg19 | released |
| CHD2-human | CHD2-human | ENCFF002COO | 432 | bed narrowPeak | optimal idr thresholded peaks | ENCSR000DZR | ChIP-seq | GM12878 | cell line | 29/10/2011 | ENCODE | hg19 | archived |
| ELF1-human | ELF1-human | ENCFF807AKG | 1283 | bed narrowPeak | optimal idr thresholded peaks | ENCSR841NDX | ChIP-seq | GM12878 | cell line | 26/06/2017 | ENCODE | hg19 | released |
| POLR3G-human | POLR3G-human | ENCFF002CPJ | 456 | bed narrowPeak | optimal idr thresholded peaks | ENCSR000EYU | ChIP-seq | GM12878 | cell line | 29/10/2011 | ENCODE | hg19 | released |
| MTA3-human | MTA3-human | ENCFF413VUC | 469 | bed narrowPeak | optimal idr thresholded peaks | ENCSR000BRH | ChIP-seq | GM12878 | cell line | 29/02/2012 | ENCODE | hg19 | released |
| BATF-human | BATF-human | ENCFF002CGQ | 472 | bed narrowPeak | optimal idr thresholded peaks | ENCSR000BGT | ChIP-seq | GM12878 | cell line | 18/07/2011 | ENCODE | hg19 | archived |
| NFYB-human | NFYB-human | ENCFF002CPC | 484 | bed narrowPeak | optimal idr thresholded peaks | ENCSR000DNM | ChIP-seq | GM12878 | cell line | 29/10/2011 | ENCODE | hg19 | archived |
| EED-human | EED-human | ENCFF242VUO | 504 | bed narrowPeak | optimal idr thresholded peaks | ENCSR199WXF | ChIP-seq | GM12878 | cell line | 30/03/2016 | ENCODE | hg19 | released |
| SUZ12-human | SUZ12-human | ENCFF746JPU | 510 | bed narrowPeak | pseudoreplicated idr thresholded peaks | ENCSR091BOQ | ChIP-seq | GM12878 | cell line | 17/01/2018 | ENCODE | hg19 | released |
| TRIM22-human | TRIM22-human | ENCFF899JBI | 996 | bed narrowPeak | optimal idr thresholded peaks | ENCSR835XKS | ChIP-seq | GM12878 | cell line | 28/07/2016 | ENCODE | hg19 | released |
| CBX3-human | CBX3-human | ENCFF141LEI | 534 | bed narrowPeak | optimal idr thresholded peaks | ENCSR549NPZ | ChIP-seq | GM12878 | cell line | 23/11/2016 | ENCODE | hg19 | released |
| ATF7-human | ATF7-human | ENCFF726VEK | 547 | bed narrowPeak | optimal idr thresholded peaks | ENCSR014YCR | ChIP-seq | GM12878 | cell line | 08/07/2016 | ENCODE | hg19 | released |
| TBX21-human | TBX21-human | ENCFF405NFV | 557 | bed narrowPeak | optimal idr thresholded peaks | ENCSR739IHN | ChIP-seq | GM12878 | cell line | 14/06/2016 | ENCODE | hg19 | released |
| SMC3-human | SMC3-human | ENCFF002CPN | 559 | bed narrowPeak | optimal idr thresholded peaks | ENCSR000DZP | ChIP-seq | GM12878 | cell line | 29/10/2011 | ENCODE | hg19 | archived |
| ZBTB40-human | ZBTB40-human | ENCFF357WYN | 574 | bed narrowPeak | optimal idr thresholded peaks | ENCSR189YYK | ChIP-seq | GM12878 | cell line | 20/09/2016 | ENCODE | hg19 | released |
| EP300-human | EP300-human | ENCFF002CPE | 1393 | bed narrowPeak | optimal idr thresholded peaks | ENCSR000DZD | ChIP-seq | GM12878 | cell line | 14/05/2012 | ENCODE | hg19 | archived |
| REST-human | REST-human | ENCFF002CHH | 593 | bed narrowPeak | optimal idr thresholded peaks | ENCSR000BQS | ChIP-seq | GM12878 | cell line | 29/02/2012 | ENCODE | hg19 | archived |
| ZNF207-human | ZNF207-human | ENCFF273BVT | 612 | bed narrowPeak | optimal idr thresholded peaks | ENCSR117KWH | ChIP-seq | GM12878 | cell line | 07/07/2014 | ENCODE | hg19 | released |
| EBF1-human | EBF1-human | ENCFF002COS | 615 | bed narrowPeak | optimal idr thresholded peaks | ENCSR000DZQ | ChIP-seq | GM12878 | cell line | 29/10/2011 | ENCODE | hg19 | archived |
| EZH2-human | EZH2-human | ENCFF002CDQ | 627 | bed narrowPeak | optimal idr thresholded peaks | ENCSR000ARD | ChIP-seq | GM12878 | cell line | 06/03/2012 | ENCODE | hg19 | archived |
| BMI1-human | BMI1-human | ENCFF809PXE | 643 | bed narrowPeak | optimal idr thresholded peaks | ENCSR469WII | ChIP-seq | GM12878 | cell line | 18/11/2016 | ENCODE | hg19 | released |
| RFX5-human | RFX5-human | ENCFF002CPL | 649 | bed narrowPeak | optimal idr thresholded peaks | ENCSR000DZW | ChIP-seq | GM12878 | cell line | 29/10/2011 | ENCODE | hg19 | archived |
| RUNX3-human | RUNX3-human | ENCFF002CHS | 662 | bed narrowPeak | optimal idr thresholded peaks | ENCSR000BRI | ChIP-seq | GM12878 | cell line | 29/02/2012 | ENCODE | hg19 | archived |
| ATF2-human | ATF2-human | ENCFF609KOY | 1427 | bed narrowPeak | optimal idr thresholded peaks | ENCSR000BQK | ChIP-seq | GM12878 | cell line | 29/02/2012 | ENCODE | hg19 | released |
| SRF-human | SRF-human | ENCFF909FRA | 864 | bed narrowPeak | optimal idr thresholded peaks | ENCSR041XML | ChIP-seq | GM12878 | cell line | 26/01/2018 | ENCODE | hg19 | released |
| ZNF143-human | ZNF143-human | ENCFF141SAU | 702 | bed narrowPeak | optimal idr thresholded peaks | ENCSR000DZL | ChIP-seq | GM12878 | cell line | 29/10/2011 | ENCODE | hg19 | released |
| E4F1-human | E4F1-human | ENCFF711OSR | 712 | bed narrowPeak | optimal idr thresholded peaks | ENCSR439WAF | ChIP-seq | GM12878 | cell line | 26/01/2018 | ENCODE | hg19 | released |
| SMAD5-human | SMAD5-human | ENCFF134XDY | 722 | bed narrowPeak | optimal idr thresholded peaks | ENCSR251OVJ | ChIP-seq | GM12878 | cell line | 02/02/2016 | ENCODE | hg19 | released |
| PAX8-human | PAX8-human | ENCFF455YHJ | 737 | bed narrowPeak | optimal idr thresholded peaks | ENCSR192AFN | ChIP-seq | GM12878 | cell line | 30/11/2017 | ENCODE | hg19 | released |
| PAX8-human | PAX8-human | ENCFF455YHJ | 737 | bed narrowPeak | optimal idr thresholded peaks | ENCSR192AFN | ChIP-seq | GM12878 | cell line | 30/11/2017 | ENCODE | hg19 | released |
| MEF2B-human | MEF2B-human | ENCFF911BYP | 764 | bed narrowPeak | optimal idr thresholded peaks | ENCSR177VFS | ChIP-seq | GM12878 | cell line | 13/02/2018 | ENCODE | hg19 | released |
| STAT3-human | STAT3-human | ENCFF002CPP | 770 | bed narrowPeak | optimal idr thresholded peaks | ENCSR000DZV | ChIP-seq | GM12878 | cell line | 29/10/2011 | ENCODE | hg19 | archived |
| MTA2-human | MTA2-human | ENCFF966RPL | 784 | bed narrowPeak | optimal idr thresholded peaks | ENCSR293QAR | ChIP-seq | GM12878 | cell line | 20/09/2016 | ENCODE | hg19 | released |
| CBFB-human | CBFB-human | ENCFF788LSH | 800 | bed narrowPeak | optimal idr thresholded peaks | ENCSR860UHK | ChIP-seq | GM12878 | cell line | 13/06/2016 | ENCODE | hg19 | released |
| FOS-human | FOS-human | ENCFF002COM | 802 | bed narrowPeak | optimal idr thresholded peaks | ENCSR000EYZ | ChIP-seq | GM12878 | cell line | 29/10/2011 | ENCODE | hg19 | released |
| PBX3-human | PBX3-human | ENCFF002CHL | 805 | bed narrowPeak | optimal idr thresholded peaks | ENCSR000BGR | ChIP-seq | GM12878 | cell line | 18/07/2011 | ENCODE | hg19 | archived |
| CBX5-human | CBX5-human | ENCFF420YHY | 816 | bed narrowPeak | optimal idr thresholded peaks | ENCSR372GIN | ChIP-seq | GM12878 | cell line | 27/10/2016 | ENCODE | hg19 | released |
| IKZF2-human | IKZF2-human | ENCFF653MIQ | 845 | bed narrowPeak | optimal idr thresholded peaks | ENCSR822AHX | ChIP-seq | GM12878 | cell line | 23/06/2017 | ENCODE | hg19 | released |
| HDAC6-human | HDAC6-human | ENCFF109YBE | 847 | bed narrowPeak | pseudoreplicated idr thresholded peaks | ENCSR933EYC | ChIP-seq | GM12878 | cell line | 14/12/2017 | ENCODE | hg19 | released |
| BACH1-human | BACH1-human | ENCFF012JXJ | 859 | bed narrowPeak | optimal idr thresholded peaks | ENCSR636MKU | ChIP-seq | GM12878 | cell line | 31/01/2017 | ENCODE | hg19 | released |
| NFIC-human | NFIC-human | ENCFF628QJU | 881 | bed narrowPeak | optimal idr thresholded peaks | ENCSR000BRN | ChIP-seq | GM12878 | cell line | 29/02/2012 | ENCODE | hg19 | released |
| NBN-human | NBN-human | ENCFF055NJR | 883 | bed narrowPeak | optimal idr thresholded peaks | ENCSR278SQL | ChIP-seq | GM12878 | cell line | 24/08/2016 | ENCODE | hg19 | released |
| IRF5-human | IRF5-human | ENCFF100ERQ | 897 | bed narrowPeak | optimal idr thresholded peaks | ENCSR976TBC | ChIP-seq | GM12878 | cell line | 21/02/2017 | ENCODE | hg19 | released |
| E2F8-human | E2F8-human | ENCFF063WXY | 907 | bed narrowPeak | optimal idr thresholded peaks | ENCSR793HVL | ChIP-seq | GM12878 | cell line | 26/01/2018 | ENCODE | hg19 | released |
| SKIL-human | SKIL-human | ENCFF142EVF | 917 | bed narrowPeak | optimal idr thresholded peaks | ENCSR212YKD | ChIP-seq | GM12878 | cell line | 24/07/2017 | ENCODE | hg19 | released |
| TCF12-human | TCF12-human | ENCFF160WLU | 935 | bed narrowPeak | optimal idr thresholded peaks | ENCSR000BGZ | ChIP-seq | GM12878 | cell line | 18/07/2011 | ENCODE | hg19 | released |
| CEBPZ-human | CEBPZ-human | ENCFF920HDC | 945 | bed narrowPeak | optimal idr thresholded peaks | ENCSR347NOB | ChIP-seq | GM12878 | cell line | 02/03/2016 | ENCODE | hg19 | released |
| GATAD2B-human | GATAD2B-human | ENCFF456YII | 957 | bed narrowPeak | optimal idr thresholded peaks | ENCSR828NCB | ChIP-seq | GM12878 | cell line | 17/05/2017 | ENCODE | hg19 | released |
| DPF2-human | DPF2-human | ENCFF349MPF | 966 | bed narrowPeak | optimal idr thresholded peaks | ENCSR509FWH | ChIP-seq | GM12878 | cell line | 17/05/2017 | ENCODE | hg19 | released |
| IRF3-human | IRF3-human | ENCFF341HWZ | 1790 | bed narrowPeak | optimal idr thresholded peaks | ENCSR408JQO | ChIP-seq | GM12878 | cell line | 07/07/2014 | ENCODE | hg19 | released |
| ZSCAN29-human | ZSCAN29-human | ENCFF455HXN | 1004 | bed narrowPeak | optimal idr thresholded peaks | ENCSR412YGM | ChIP-seq | GM12878 | cell line | 17/05/2017 | ENCODE | hg19 | released |
| MLLT1-human | MLLT1-human | ENCFF173KPE | 1029 | bed narrowPeak | conservative idr thresholded peaks | ENCSR552XSN | ChIP-seq | GM12878 | cell line | 24/08/2016 | ENCODE | hg19 | released |
| ARID3A-human | ARID3A-human | ENCFF027VZK | 1040 | bed narrowPeak | optimal idr thresholded peaks | ENCSR778UBR | ChIP-seq | GM12878 | cell line | 31/01/2017 | ENCODE | hg19 | released |
| NFXL1-human | NFXL1-human | ENCFF166FRO | 1052 | bed narrowPeak | optimal idr thresholded peaks | ENCSR746XEG | ChIP-seq | GM12878 | cell line | 17/05/2017 | ENCODE | hg19 | released |
| ESRRA-human | ESRRA-human | ENCFF195PBG | 1063 | bed narrowPeak | optimal idr thresholded peaks | ENCSR000DYQ | ChIP-seq | GM12878 | cell line | 20/08/2012 | ENCODE | hg19 | released |
| ZNF622-human | ZNF622-human | ENCFF065QOM | 1070 | bed narrowPeak | optimal idr thresholded peaks | ENCSR075FNZ | ChIP-seq | GM12878 | cell line | 13/11/2017 | ENCODE | hg19 | released |
| YBX1-human | YBX1-human | ENCFF344OJI | 1079 | bed narrowPeak | optimal idr thresholded peaks | ENCSR205SKQ | ChIP-seq | GM12878 | cell line | 24/08/2016 | ENCODE | hg19 | released |
| NR2F1-human | NR2F1-human | ENCFF810CEL | 1089 | bed narrowPeak | optimal idr thresholded peaks | ENCSR514VYD | ChIP-seq | GM12878 | cell line | 14/04/2017 | ENCODE | hg19 | released |
| RELB-human | RELB-human | ENCFF456FQB | 1100 | bed narrowPeak | optimal idr thresholded peaks | ENCSR387QUV | ChIP-seq | GM12878 | cell line | 21/02/2017 | ENCODE | hg19 | released |
| NRF1-human | NRF1-human | ENCFF002CPD | 1107 | bed narrowPeak | optimal idr thresholded peaks | ENCSR000DZO | ChIP-seq | GM12878 | cell line | 29/10/2011 | ENCODE | hg19 | archived |
| NR2C1-human | NR2C1-human | ENCFF381SBC | 1124 | bed narrowPeak | optimal idr thresholded peaks | ENCSR784VIQ | ChIP-seq | GM12878 | cell line | 07/06/2017 | ENCODE | hg19 | released |
| MYC-human | MYC-human | ENCFF002DAI | 1127 | bed narrowPeak | optimal idr thresholded peaks | ENCSR000DKU | ChIP-seq | GM12878 | cell line | 17/03/2011 | ENCODE | hg19 | released |
| GABPA-human | GABPA-human | ENCFF116EXQ | 1129 | bed narrowPeak | optimal idr thresholded peaks | ENCSR331HPA | ChIP-seq | GM12878 | cell line | 30/06/2014 | ENCODE | hg19 | released |
| ZNF687-human | ZNF687-human | ENCFF643OWO | 1151 | bed narrowPeak | optimal idr thresholded peaks | ENCSR859FDL | ChIP-seq | GM12878 | cell line | 26/06/2017 | ENCODE | hg19 | released |
| E2F4-human | E2F4-human | ENCFF002COR | 1159 | bed narrowPeak | optimal idr thresholded peaks | ENCSR000DYY | ChIP-seq | GM12878 | cell line | 14/05/2012 | ENCODE | hg19 | archived |
| CUX1-human | CUX1-human | ENCFF721SDD | 1175 | bed narrowPeak | optimal idr thresholded peaks | ENCSR000DYR | ChIP-seq | GM12878 | cell line | 20/08/2012 | ENCODE | hg19 | released |
| MEF2A-human | MEF2A-human | ENCFF328QLX | 1201 | bed narrowPeak | optimal idr thresholded peaks | ENCSR000BKB | ChIP-seq | GM12878 | cell line | 18/07/2011 | ENCODE | hg19 | released |
| ZNF217-human | ZNF217-human | ENCFF513RIQ | 1218 | bed narrowPeak | optimal idr thresholded peaks | ENCSR764CZW | ChIP-seq | GM12878 | cell line | 07/06/2017 | ENCODE | hg19 | released |
| SIX5-human | SIX5-human | ENCFF002CHU | 1239 | bed narrowPeak | optimal idr thresholded peaks | ENCSR000BJE | ChIP-seq | GM12878 | cell line | 18/07/2011 | ENCODE | hg19 | archived |
| USF2-human | USF2-human | ENCFF002CPT | 1251 | bed narrowPeak | optimal idr thresholded peaks | ENCSR000DZU | ChIP-seq | GM12878 | cell line | 29/10/2011 | ENCODE | hg19 | archived |
| SMARCA5-human | SMARCA5-human | ENCFF478DRD | 1260 | bed narrowPeak | optimal idr thresholded peaks | ENCSR706YUH | ChIP-seq | GM12878 | cell line | 07/06/2017 | ENCODE | hg19 | released |
| ZNF592-human | ZNF592-human | ENCFF566YUW | 1276 | bed narrowPeak | optimal idr thresholded peaks | ENCSR173ZVL | ChIP-seq | GM12878 | cell line | 26/06/2017 | ENCODE | hg19 | released |
| ETS1-human | ETS1-human | ENCFF332PGQ | 1301 | bed narrowPeak | optimal idr thresholded peaks | ENCSR000BKA | ChIP-seq | GM12878 | cell line | 18/07/2011 | ENCODE | hg19 | released |
| ZNF24-human | ZNF24-human | ENCFF103OOV | 1311 | bed narrowPeak | optimal idr thresholded peaks | ENCSR072PWP | ChIP-seq | GM12878 | cell line | 26/06/2017 | ENCODE | hg19 | released |
| RELA-human | RELA-human | ENCFF002CPA | 1326 | bed narrowPeak | optimal idr thresholded peaks | ENCSR000EAG | ChIP-seq | GM12878 | cell line | 14/05/2012 | ENCODE | hg19 | released |
| IRF4-human | IRF4-human | ENCFF391DVL | 1339 | bed narrowPeak | optimal idr thresholded peaks | ENCSR000BGY | ChIP-seq | GM12878 | cell line | 18/07/2011 | ENCODE | hg19 | released |
| ELK1-human | ELK1-human | ENCFF002COT | 1354 | bed narrowPeak | optimal idr thresholded peaks | ENCSR000DZB | ChIP-seq | GM12878 | cell line | 14/05/2012 | ENCODE | hg19 | archived |
| MXI1-human | MXI1-human | ENCFF002COY | 1368 | bed narrowPeak | optimal idr thresholded peaks | ENCSR000DZI | ChIP-seq | GM12878 | cell line | 29/10/2011 | ENCODE | hg19 | archived |
| ETV6-human | ETV6-human | ENCFF760IFZ | 1405 | bed narrowPeak | optimal idr thresholded peaks | ENCSR626VUC | ChIP-seq | GM12878 | cell line | 07/07/2014 | ENCODE | hg19 | released |
| MYB-human | MYB-human | ENCFF215YWS | 1436 | bed narrowPeak | optimal idr thresholded peaks | ENCSR819ATC | ChIP-seq | GM12878 | cell line | 07/07/2014 | ENCODE | hg19 | released |
| MAX-human | MAX-human | ENCFF083KVY | 1453 | bed narrowPeak | optimal idr thresholded peaks | ENCSR000DZF | ChIP-seq | GM12878 | cell line | 14/05/2012 | ENCODE | hg19 | released |
| ZZZ3-human | ZZZ3-human | ENCFF002CPY | 1460 | bed narrowPeak | optimal idr thresholded peaks | ENCSR000DNQ | ChIP-seq | GM12878 | cell line | 29/10/2011 | ENCODE | hg19 | archived |
| SMAD1-human | SMAD1-human | ENCFF632MBO | 1477 | bed narrowPeak | optimal idr thresholded peaks | ENCSR813DCK | ChIP-seq | GM12878 | cell line | 26/01/2018 | ENCODE | hg19 | released |
| ZNF384-human | ZNF384-human | ENCFF371OGO | 1486 | bed narrowPeak | optimal idr thresholded peaks | ENCSR000DYP | ChIP-seq | GM12878 | cell line | 20/08/2012 | ENCODE | hg19 | released |
| FOXK2-human | FOXK2-human | ENCFF980KSV | 1501 | bed narrowPeak | optimal idr thresholded peaks | ENCSR861JUQ | ChIP-seq | GM12878 | cell line | 27/01/2017 | ENCODE | hg19 | released |
| POLR2A-human | POLR2A-human | ENCFF120VUT | 1509 | bed narrowPeak | optimal idr thresholded peaks | ENCSR000BGD | ChIP-seq | GM12878 | cell line | 18/07/2011 | ENCODE | hg19 | released |
| LARP7-human | LARP7-human | ENCFF449CJP | 1521 | bed narrowPeak | optimal idr thresholded peaks | ENCSR657PEW | ChIP-seq | GM12878 | cell line | 17/05/2017 | ENCODE | hg19 | released |
| NKRF-human | NKRF-human | ENCFF317VBQ | 1533 | bed narrowPeak | optimal idr thresholded peaks | ENCSR732PJX | ChIP-seq | GM12878 | cell line | 17/05/2017 | ENCODE | hg19 | released |
| FOXM1-human | FOXM1-human | ENCFF002CGZ | 1567 | bed narrowPeak | optimal idr thresholded peaks | ENCSR000BRU | ChIP-seq | GM12878 | cell line | 29/02/2012 | ENCODE | hg19 | archived |
| TBL1XR1-human | TBL1XR1-human | ENCFF002CPQ | 1569 | bed narrowPeak | optimal idr thresholded peaks | ENCSR000DYZ | ChIP-seq | GM12878 | cell line | 14/05/2012 | ENCODE | hg19 | archived |
| RB1-human | RB1-human | ENCFF601XVA | 1585 | bed narrowPeak | optimal idr thresholded peaks | ENCSR785OKZ | ChIP-seq | GM12878 | cell line | 07/06/2017 | ENCODE | hg19 | released |
| BCL3-human | BCL3-human | ENCFF625SHY | 1616 | bed narrowPeak | optimal idr thresholded peaks | ENCSR000BNQ | ChIP-seq | GM12878 | cell line | 18/07/2011 | ENCODE | hg19 | released |
| HCFC1-human | HCFC1-human | ENCFF208TRU | 1621 | bed narrowPeak | optimal idr thresholded peaks | ENCSR514VAY | ChIP-seq | GM12878 | cell line | 07/07/2014 | ENCODE | hg19 | released |
| KLF5-human | KLF5-human | ENCFF827ZLB | 1641 | bed narrowPeak | optimal idr thresholded peaks | ENCSR974OFJ | ChIP-seq | GM12878 | cell line | 07/07/2014 | ENCODE | hg19 | released |
| TAF1-human | TAF1-human | ENCFF544NRC | 1653 | bed narrowPeak | optimal idr thresholded peaks | ENCSR000BGS | ChIP-seq | GM12878 | cell line | 18/07/2011 | ENCODE | hg19 | released |
| NFATC1-human | NFATC1-human | ENCFF409QJU | 1667 | bed narrowPeak | optimal idr thresholded peaks | ENCSR000BQL | ChIP-seq | GM12878 | cell line | 29/02/2012 | ENCODE | hg19 | released |
| WRNIP1-human | WRNIP1-human | ENCFF002CPU | 1681 | bed narrowPeak | optimal idr thresholded peaks | ENCSR000EAA | ChIP-seq | GM12878 | cell line | 29/10/2011 | ENCODE | hg19 | archived |
| POLR2AphosphoS2-human | POLR2AphosphoS2-human | ENCFF630NFM | 1699 | bed narrowPeak | optimal idr thresholded peaks | ENCSR000DZK | ChIP-seq | GM12878 | cell line | 29/10/2011 | ENCODE | hg19 | released |
| HSF1-human | HSF1-human | ENCFF308ZGG | 1722 | bed narrowPeak | optimal idr thresholded peaks | ENCSR009MBP | ChIP-seq | GM12878 | cell line | 07/07/2014 | ENCODE | hg19 | released |
| UBTF-human | UBTF-human | ENCFF298UPI | 1729 | bed narrowPeak | optimal idr thresholded peaks | ENCSR459FTB | ChIP-seq | GM12878 | cell line | 07/07/2014 | ENCODE | hg19 | released |
| TCF7-human | TCF7-human | ENCFF305ICK | 1750 | bed narrowPeak | optimal idr thresholded peaks | ENCSR501DKS | ChIP-seq | GM12878 | cell line | 07/07/2014 | ENCODE | hg19 | released |
| CREM-human | CREM-human | ENCFF534CKB | 1760 | bed narrowPeak | optimal idr thresholded peaks | ENCSR839XZU | ChIP-seq | GM12878 | cell line | 07/07/2014 | ENCODE | hg19 | released |
| ZFP36-human | ZFP36-human | ENCFF144FCI | 1770 | bed narrowPeak | optimal idr thresholded peaks | ENCSR900XDB | ChIP-seq | GM12878 | cell line | 07/07/2014 | ENCODE | hg19 | released |
| POLR2AphosphoS5-human | POLR2AphosphoS5-human | ENCFF125SQT | 1812 | bed narrowPeak | optimal idr thresholded peaks | ENCSR000BIF | ChIP-seq | GM12878 | cell line | 18/07/2011 | ENCODE | hg19 | released |
