## Supplementary figures and images for "Analysis of long-range contacts across cell types outlines a core sequence determinant of 3D genome organisation"

### dm6_BG3_contact_maps_CfVsCII_chr2L2R3L3R4X.png

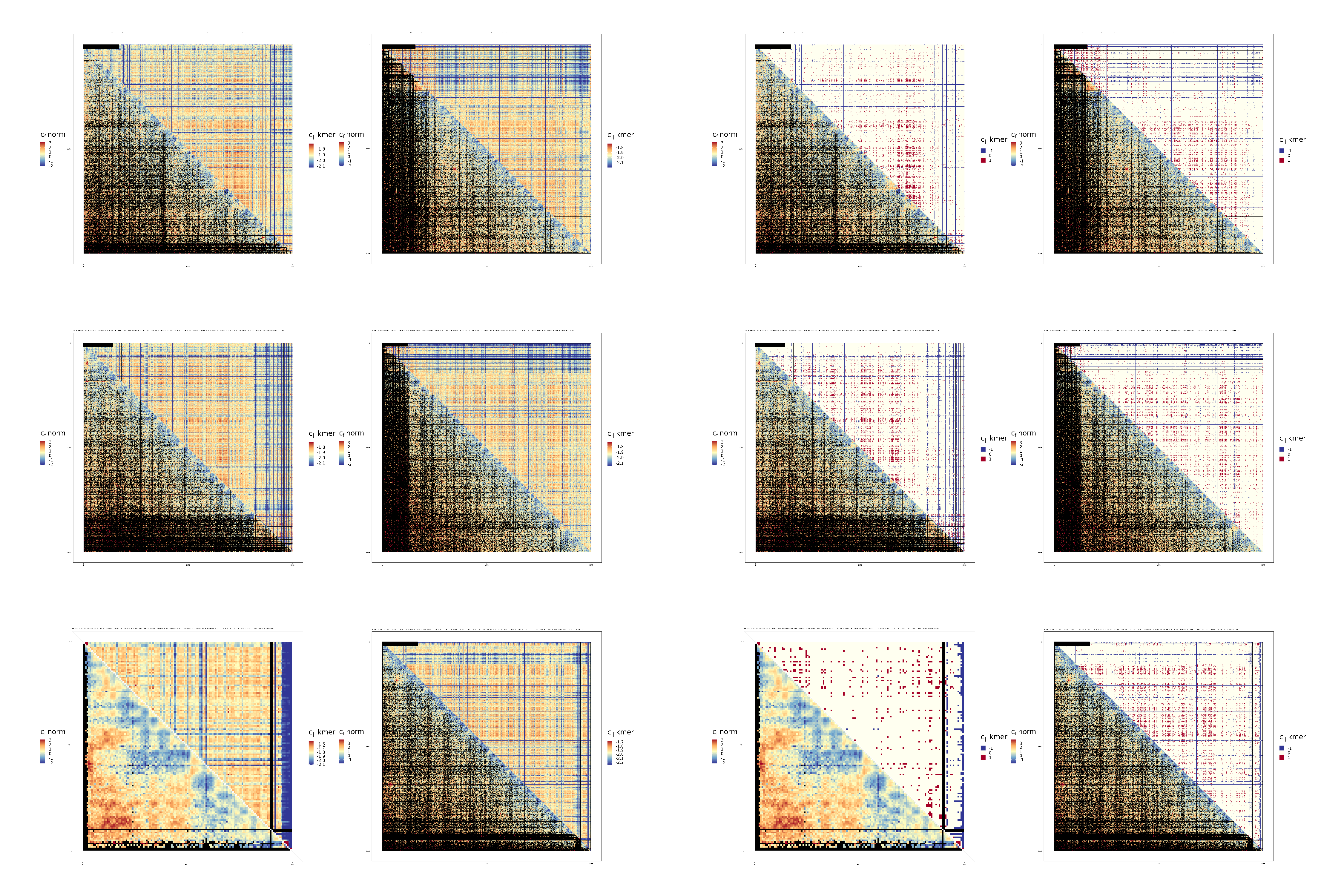

### dm6_Kc167_contact_maps_CfVsCII_chr2L2R3L3R4X.png

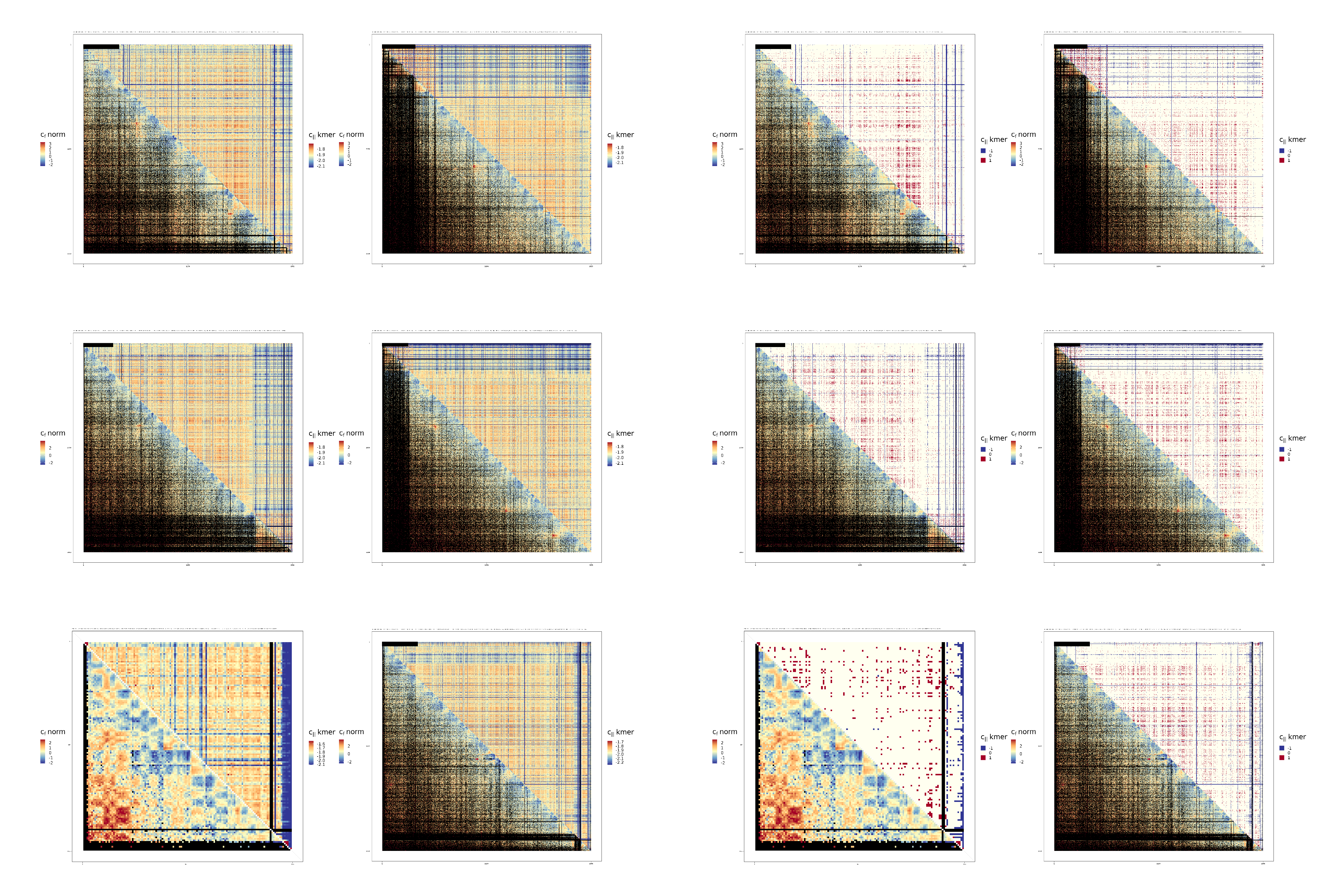

### hg37_contact_maps_CpVsCII_chr1To12.png

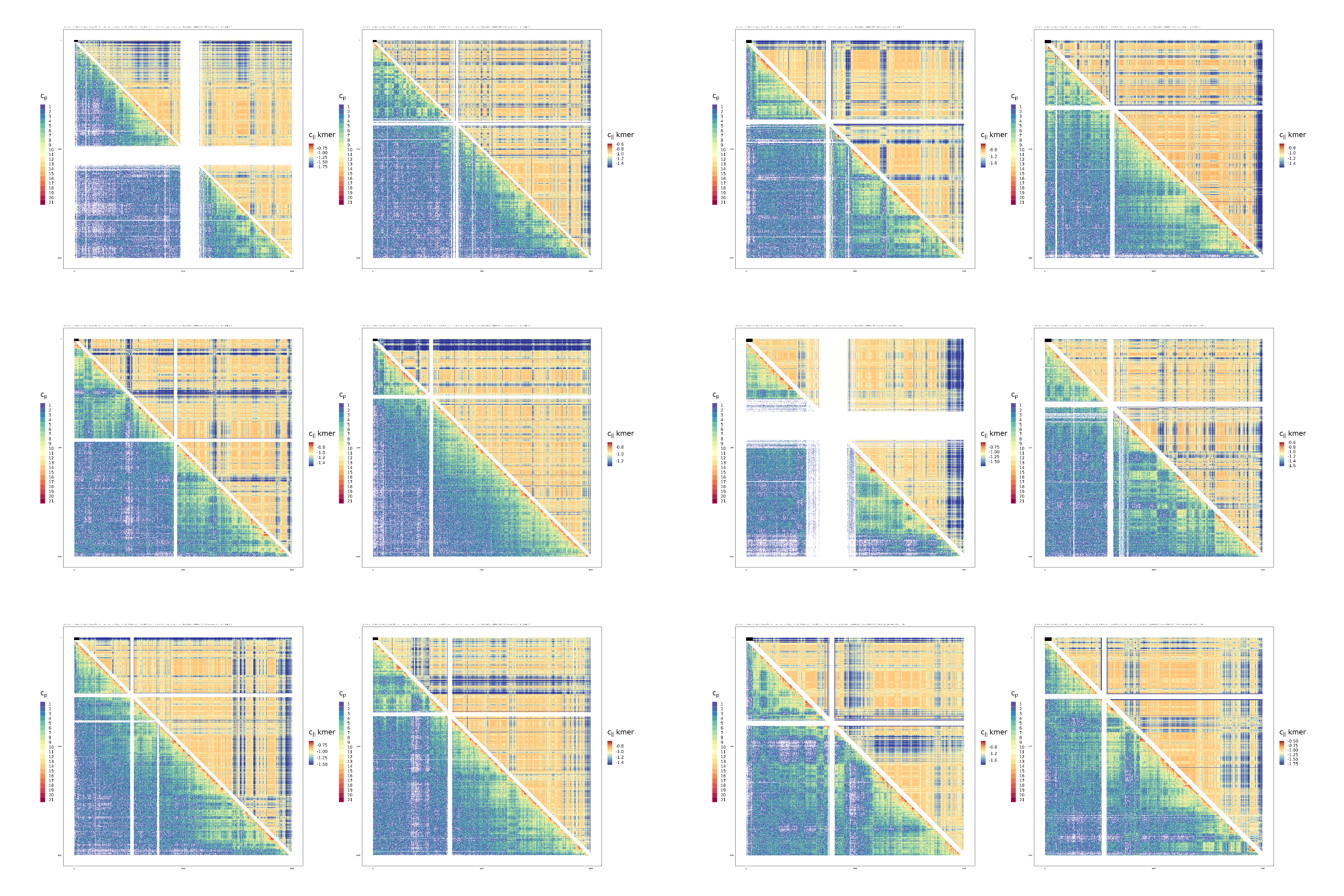

### hg37_contact_maps_CpVsCII_chr13ToX.png

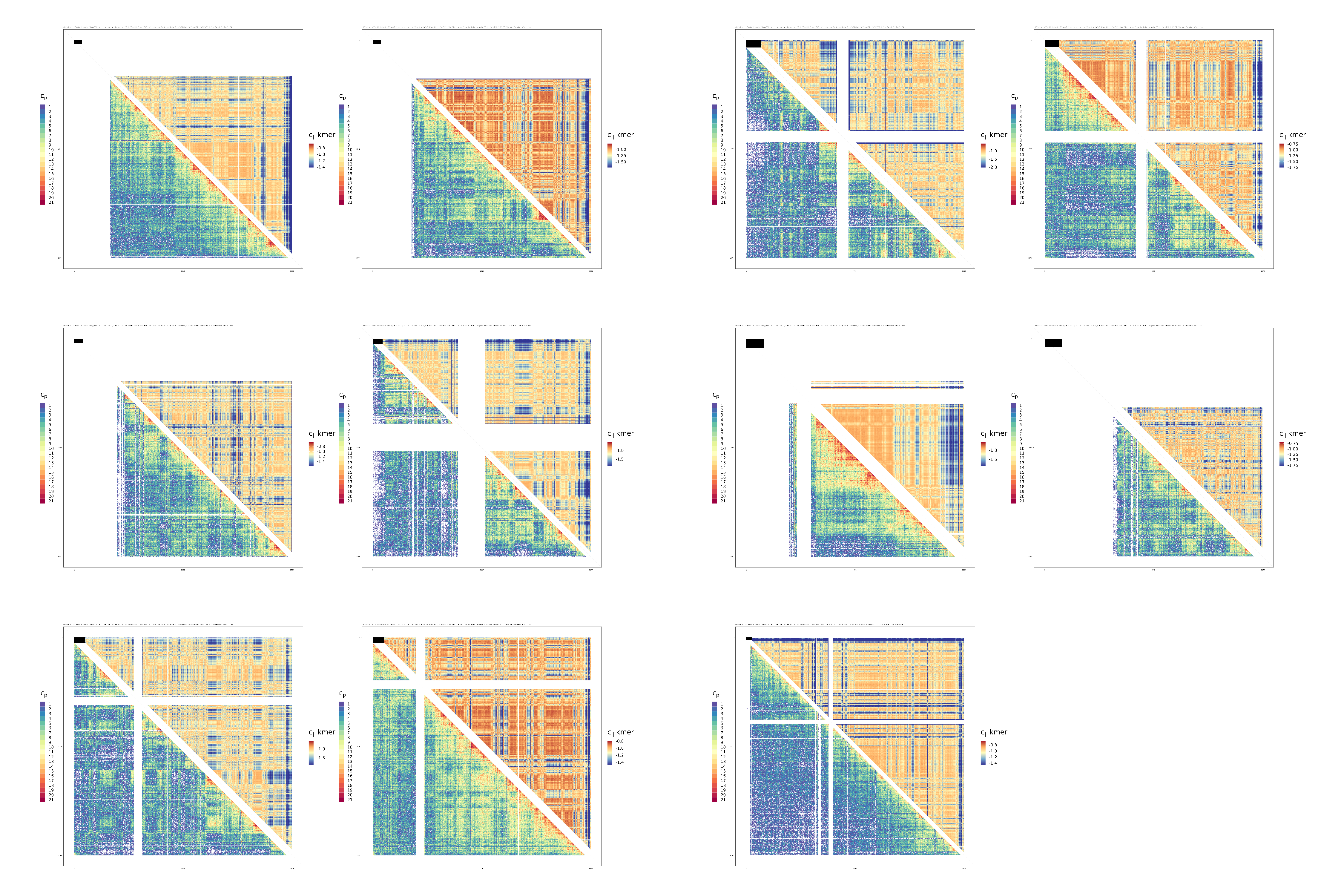

### hg38_ESCHiC_contact_maps_CfVsCII_chr1To12.png

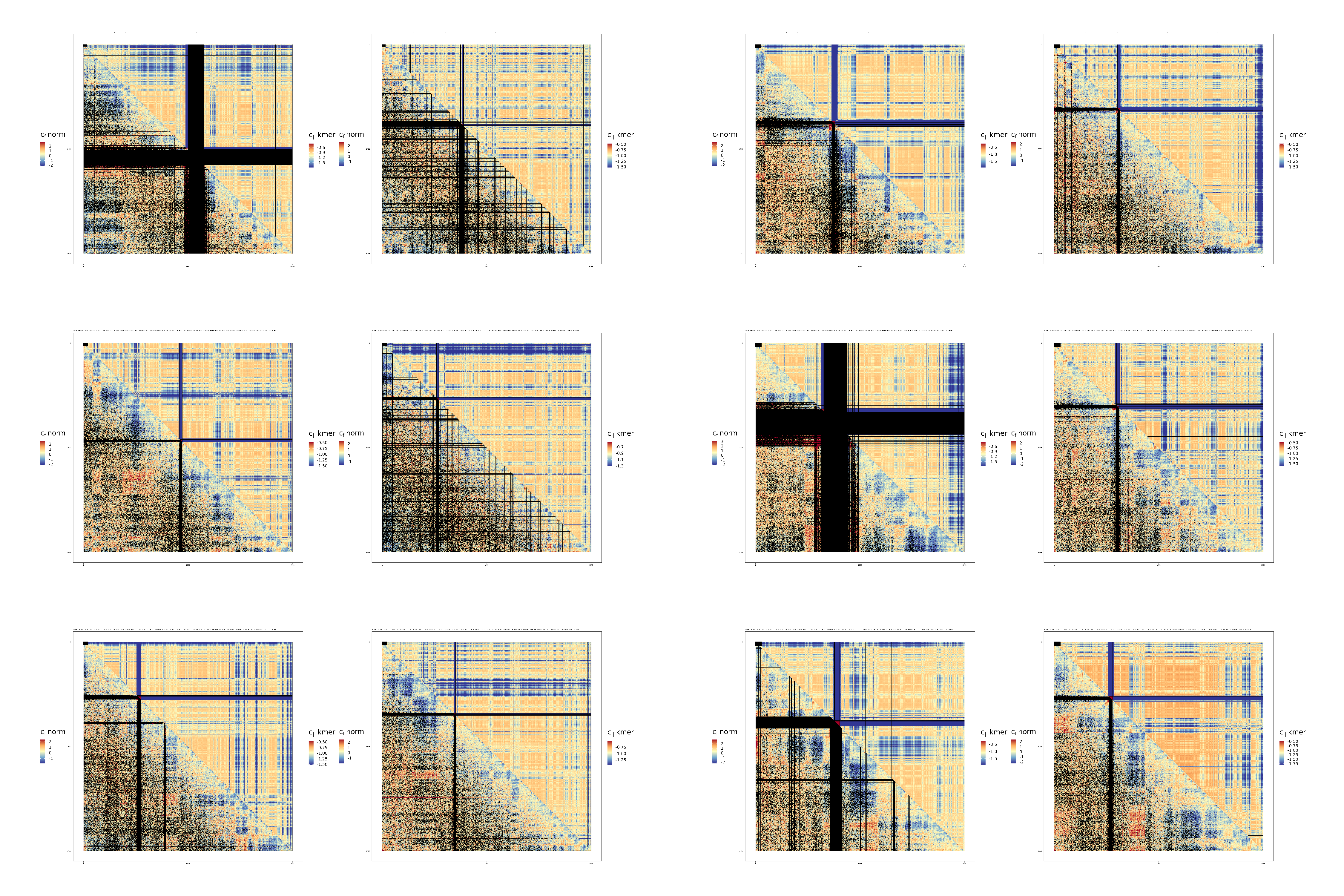

### hg38_ESCHiC_contact_maps_CfVsCII_chr13ToX.png

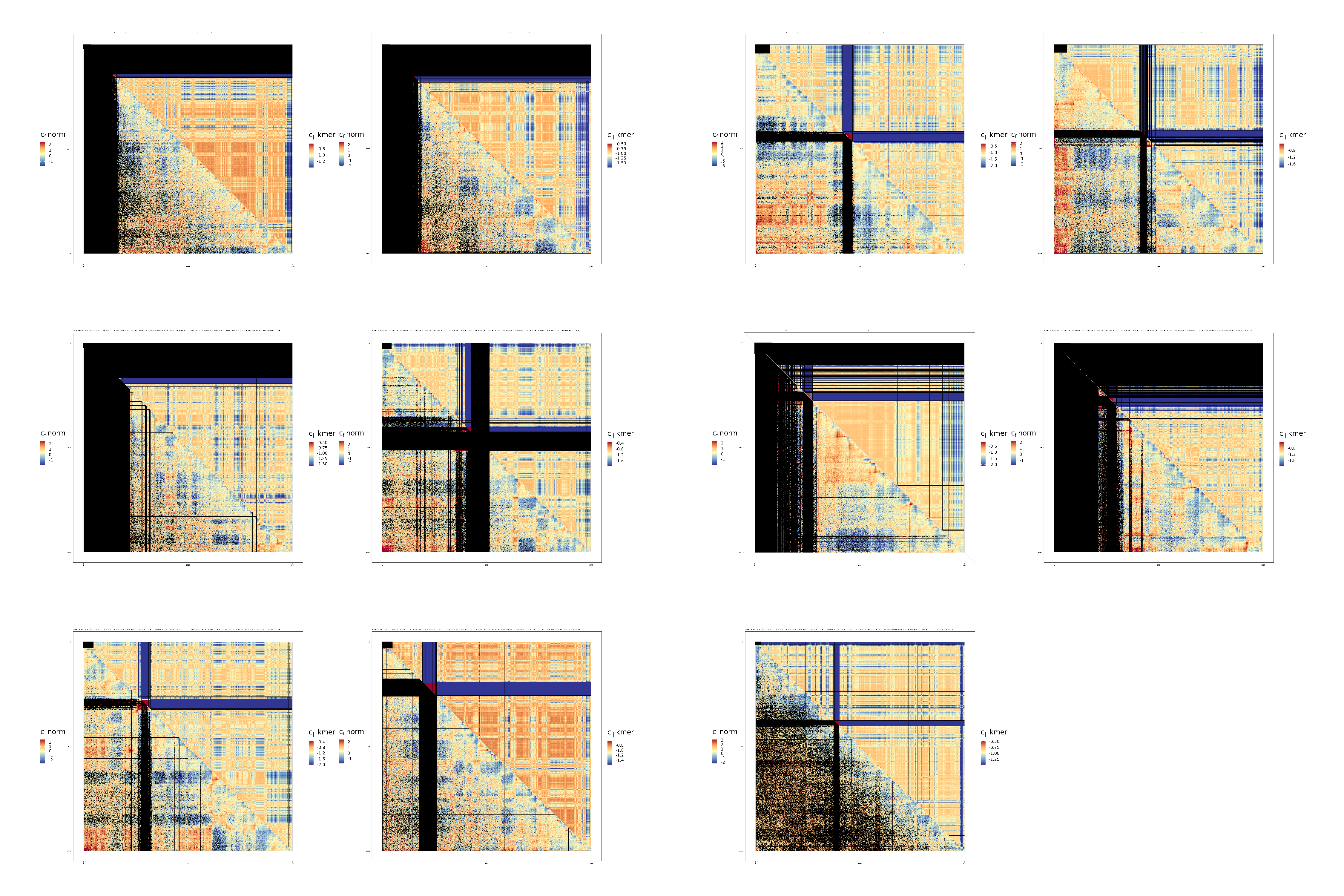

### hg38_ESCMicroC_contact_maps_CfVsCII_chr1To12.png

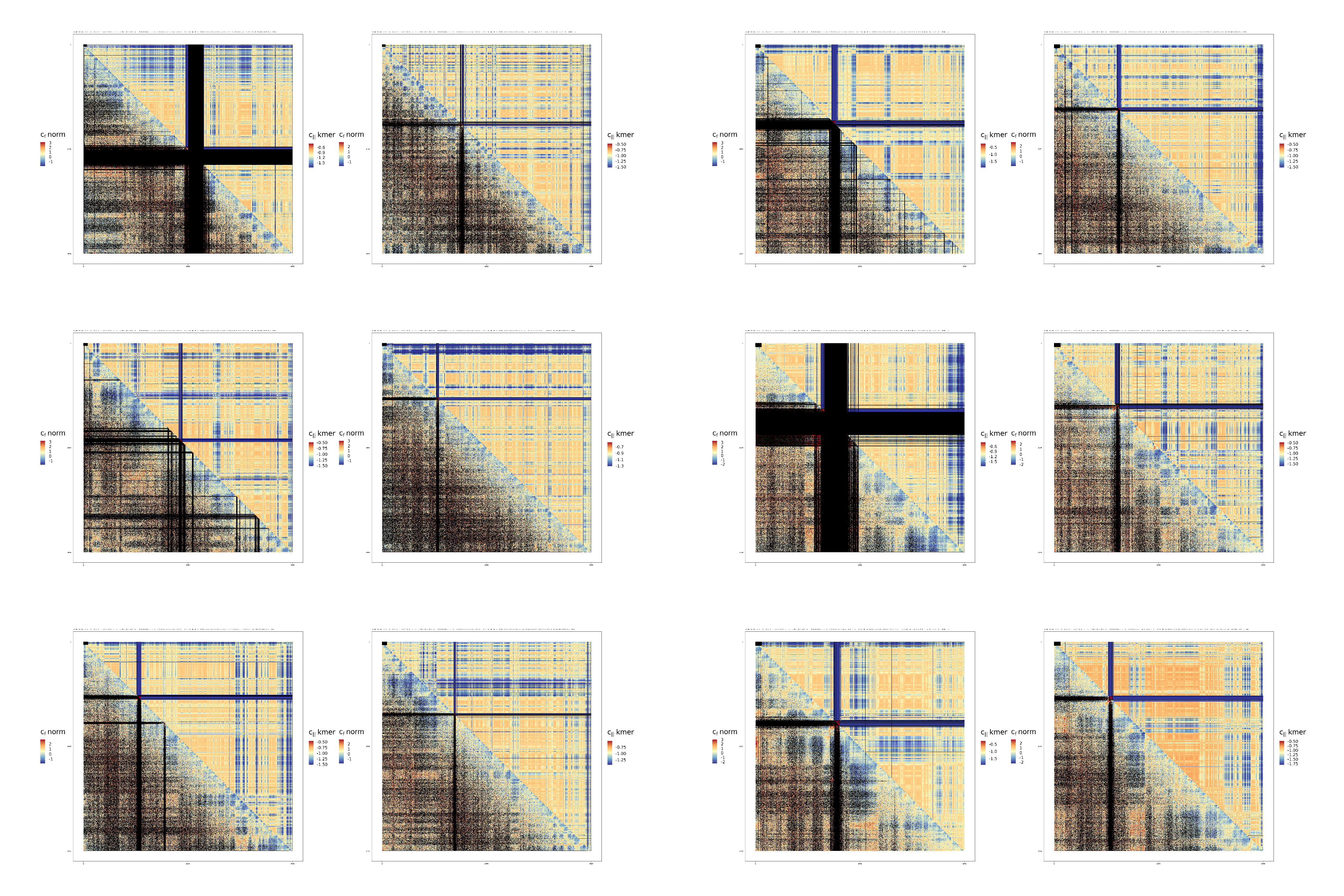

### hg38_ESCMicroC_contact_maps_CfVsCII_chr13ToX.png

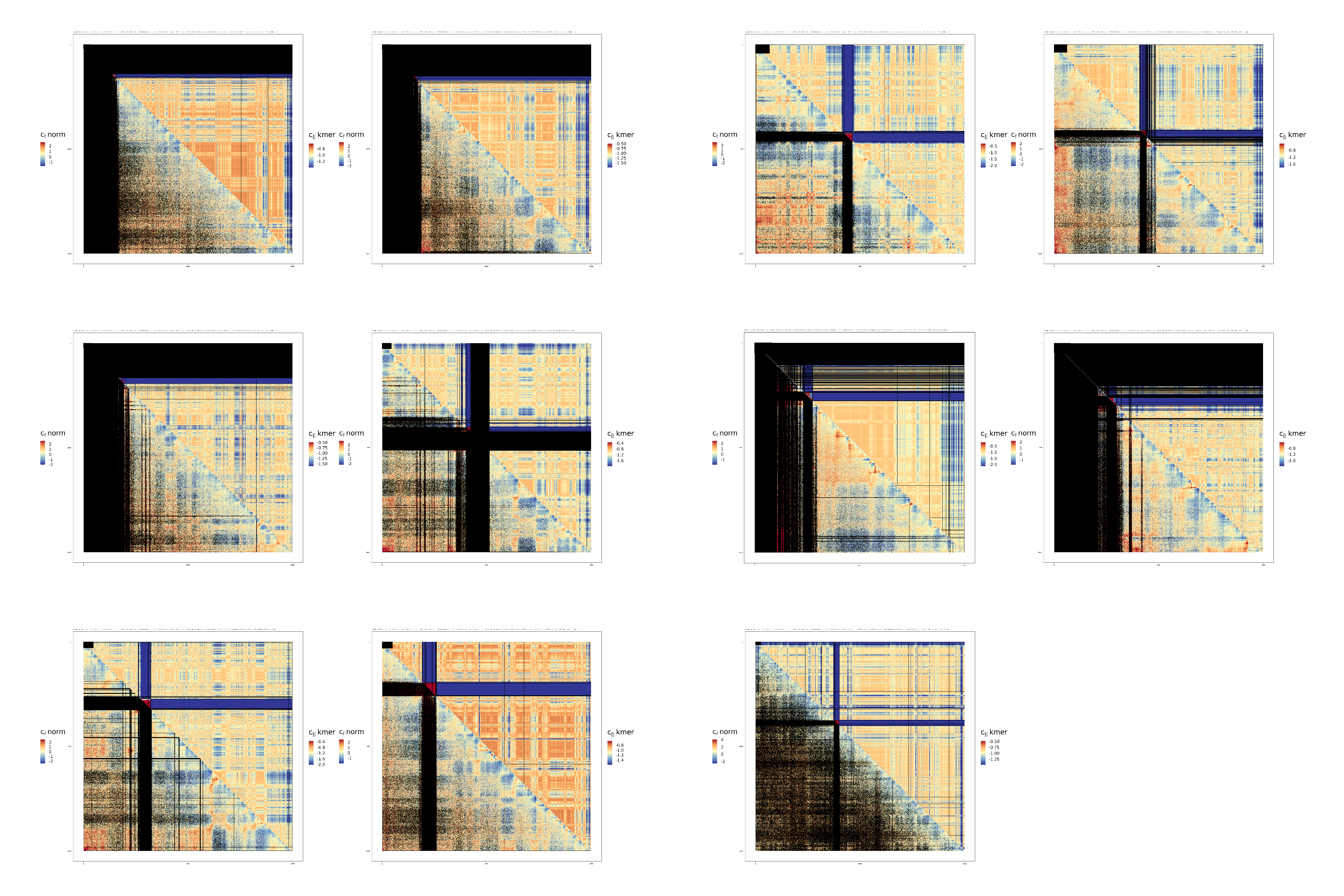

### hg38_LCHiC_contact_maps_CfVsCII_chr1To12.png

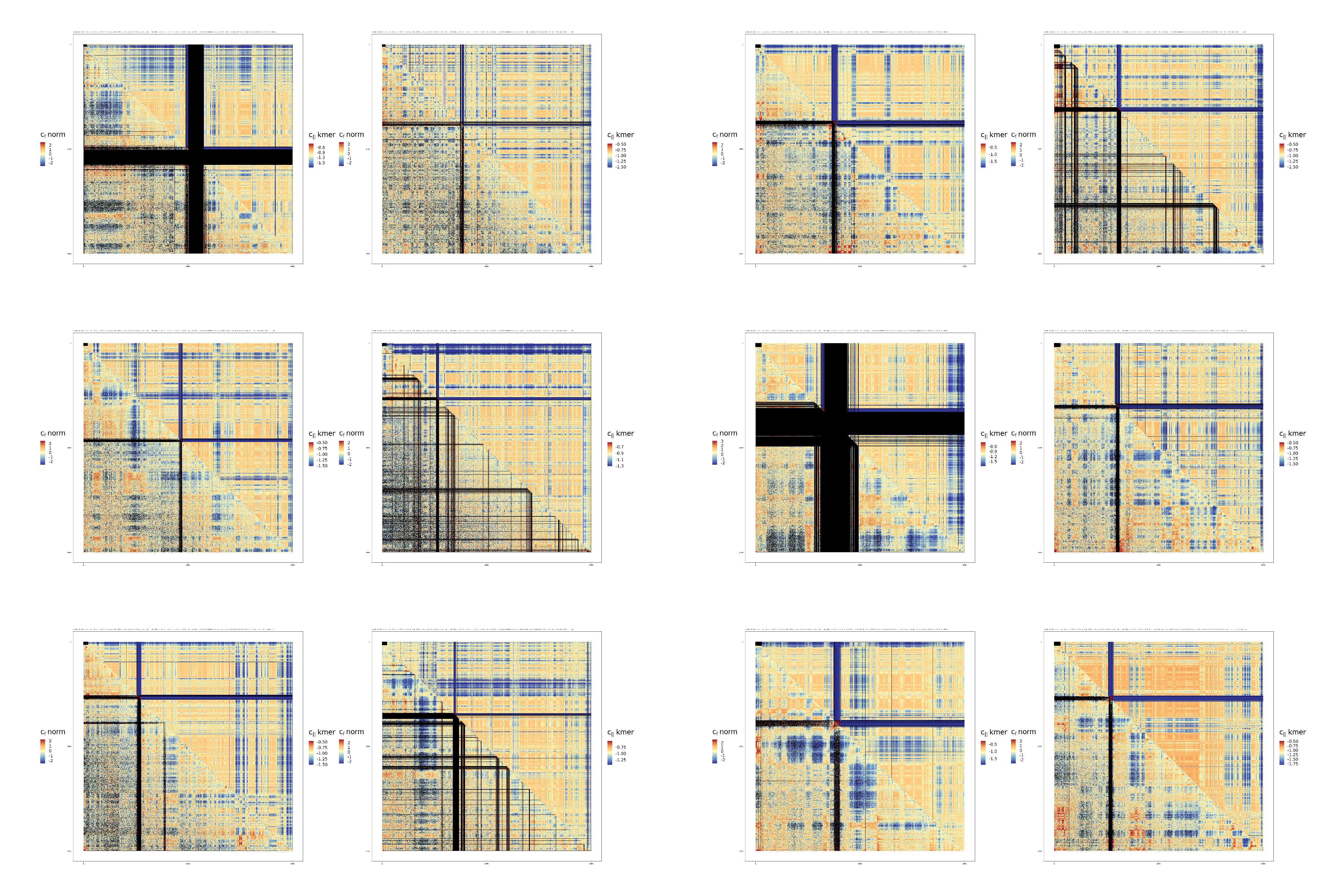

### hg38_LCHiC_contact_maps_CfVsCII_chr13ToX.png

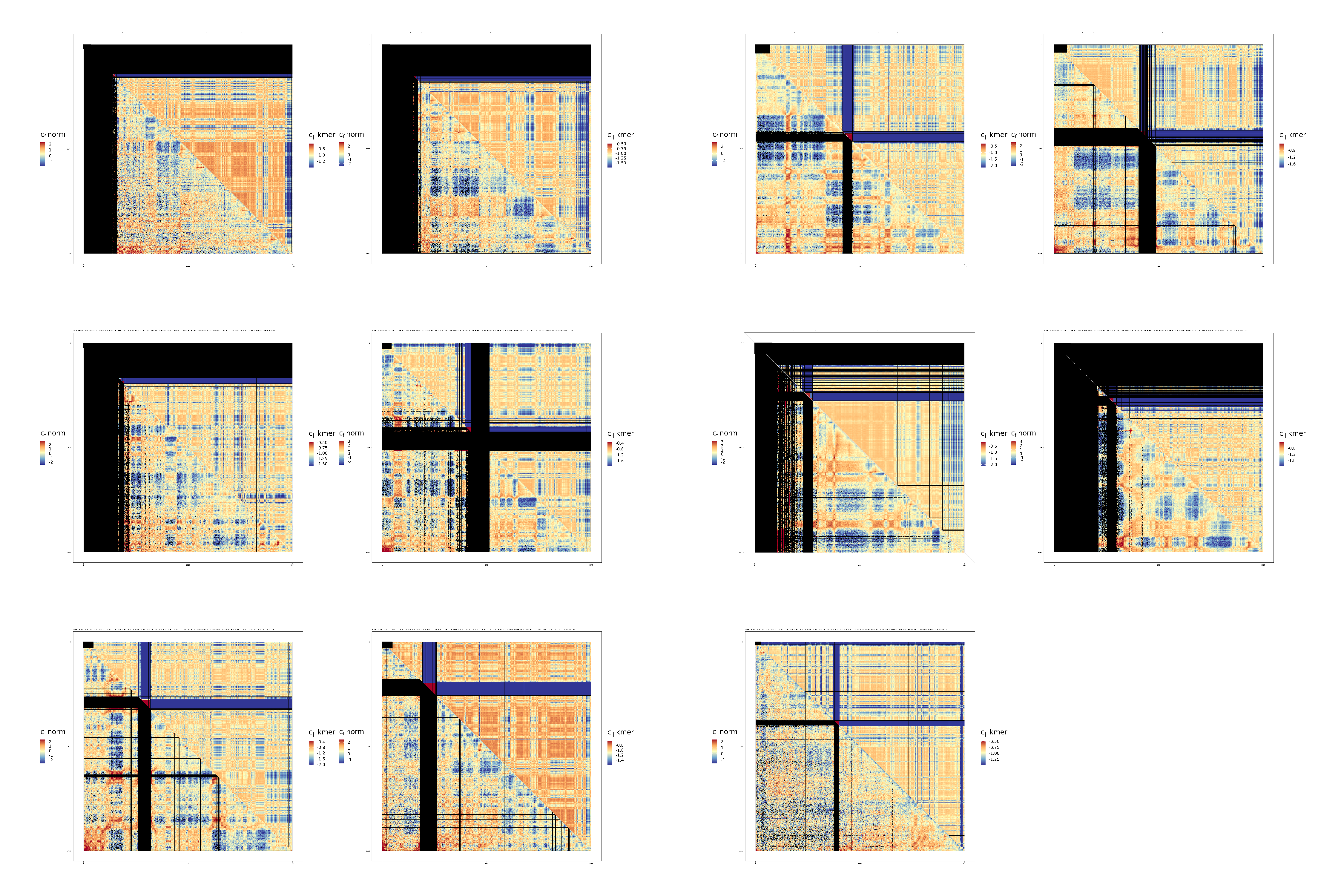
